# Mice discriminate odour source distance via sub-sniff temporal features of odour plumes

**DOI:** 10.1101/2025.05.14.653752

**Authors:** Alina Cristina Marin, Julia J. Harris, Debanjan Dasgupta, Andrew Erskine, Tom P.A. Warner, Andreas T. Schaefer, Tobias Ackels

## Abstract

Rodents rely on olfaction to navigate complex environments, particularly where visual cues are limited. Yet how they estimate the distance to an odour source remains unclear. The spatiotemporal dynamics of natural odour plumes, shaped by airflow turbulence, offer valuable cues for locating odour sources. Here, we show that mice can discriminate odour sources placed at different distances by extracting information from the sub-sniff temporal structure of naturalistic odour plumes. Using a wind tunnel and an olfactory virtual reality system, we generated dynamic plumes and demonstrated, through high-throughput automated behaviour, that mice distinguish near from far sources based on odour fluctuations operating faster than their respiratory cycle. Two-photon calcium imaging of olfactory bulb projection neurons revealed that distance-dependent responses are present in a small subset of mitral and tufted cells, and that population activity encodes source distance. Critically, neural responses correlated more strongly with high-frequency plume features than with mean odour concentration. Our results identify a neural basis for distance estimation from odour dynamics and highlight the importance of rapid temporal processing in mammalian olfaction.

## Introduction

Olfaction provides animals with the remarkable ability to remotely sample their environment under low-visibility conditions, a vital skill for crepuscular and nocturnal species, such as rodents. Rather than relying solely on direct contact, as taste does, mice detect airborne odours carried by complex, often turbulent airflows, enabling them to gather information about distant objects, such as potential food sources, predators, or mating opportunities (Marin et al., 2021; McKissick et al., 2024; Sunil et al., 2024). From an ecological standpoint, the ability to estimate how far away an odour source is can be crucial for a rodent’s survival. For instance, detecting the scent of a predator at close range prompts immediate evasive action, whereas sensing it from afar might allow continued foraging. In the same way, evaluating whether a food source is realistically within reach helps the animal to conserve energy and minimise exposure (Findley et al., 2021; Gire et al., 2016).

In natural environments, odour signals rarely spread as uniform gradients. Instead, they form spatiotemporally complex, dynamic plumes shaped by turbulent airflow (Crimaldi et al., 2022). Within these plumes, pockets and filaments of higher concentration are interspersed with intervals of relatively odour-free air (Celani et al., 2014; Crimaldi & Koseff, 2001; Moore & Atema, 1991), producing odour events on timescales of tens to hundreds of milliseconds, often exceeding typical sniffing frequencies of 4–12 Hz in rodents (Wachowiak, 2011; Welker, 1964). As a plume travels farther from its source, it expands spatially while changing its temporal statistics, including odour event parameters such as peak height, onset slope, and intermittency (Reddy et al., 2022; Schmuker et al., 2016; Vergassola et al., 2007). These features therefore carry crucial information about the distance to an odour source, offering cues to guide navigation and decision-making (Balkovsky & Shraiman, 2002; Mafra-Neto & Cardé, 1994; Rigolli et al., 2022). One key open question is whether and how rodents exploit these high-frequency fluctuations (Ackels et al., 2021; Dasgupta et al., 2022; Marin et al., 2021; Sunil et al., 2024; Tootoonian et al., 2025). If rodents can capitalise on this temporal information, it would not only enable them to pinpoint and distinguish different sources from the atmospheric background (Rokni et al., 2014), but also to construct a spatial map of their environment using purely olfactory cues (Jacobs, 2012; Poo et al., 2022; Sterrett et al., 2024). Critically, such a strategy would let them rapidly locate and identify odours without needing to physically move between sources to confirm their identity or position (Gire et al., 2016).

Behavioural evidence indicates that rodents might indeed exploit these transient cues. In certain odour localisation tasks, mice commit to a directional choice shortly after detecting the odour, well before they have traversed the entire distance to the source (Findley et al., 2021). This early commitment suggests that relevant spatial information is available in the odour signal itself, even at a distance. Moreover, when mice are trained in multi-choice arenas where multiple strategies such as memory-based foraging are possible, they sometimes switch between systematic searching and potentially more nuanced reliance on the spatiotemporal dynamics of the odour stimulus (Findley et al., 2021; Gire et al., 2016; Tariq et al., 2024). The fact that they do so under conditions of high plume complexity hints that rodents can integrate rapid temporal changes of the stimulus that might be missed by simpler gradient-following strategies (Gire et al., 2016; Jones & Urban, 2018; Khan et al., 2012).

Growing evidence suggests that each sniff captures not just a static snapshot of odour composition but can also convey intricate temporal details. In rodents, behavioural experiments with optogenetic stimulation have shown that mice can discriminate between inputs delivered as little as 10-20 ms apart (Li et al., 2014; Rebello et al., 2014; Smear et al., 2011, 2013), and asynchronous stimulation across the two olfactory bulbs can elicit different responses (Kuruppath et al., 2021). Systematic investigations of their temporal discrimination abilities have revealed that mice can discriminate odour correlation structures at frequencies up to 40 Hz, surpassing their typical sniffing rate (Ackels et al., 2021; Dasgupta et al., 2022; Warner et al., 2024). A head-restrained odour-counting task reveals that mice integrate sporadic, plume-like odour pulses across dozens of sniffs, giving greater weight to inhalation-phase stimuli which might ultimately constrain behavioural accuracy (Boero et al., 2025). Psychophysical work in humans shows that observers can distinguish the temporal order of odour pulses spaced by as little as 120 ms within a single sniff, indicating that fine-grained temporal coding extends across species (Wu et al., 2024). This capacity challenges earlier assumptions that the mammalian nasal cavity and slow transduction kinetics of olfactory receptor neurons (ORNs) would “wash out” high-frequency features of incoming odour plumes (Duchamp-Viret et al., 1999; Kepecs et al., 2006). Instead, recent physiological and modelling work indicates that the convergence of large populations of ORNs in the olfactory bulb (OB), allows access to fast-changing odour signals (Ackels et al., 2021). Correspondingly, OB projection neurons, mitral and tufted cells (MTCs), can encode odour correlation structure at frequencies up to 20 Hz (Ackels et al., 2021; Dasgupta et al., 2022). Further it has been shown that odour events such as onset, offset, whiffs, and blanks are tightly coupled to MTC population activity, with different MTC clusters exhibiting varying degrees of correlation (Lewis et al., 2021, 2024) and that mice can discriminate odour stimuli of low and high intermittency values (Gumaste et al., 2024). Such high-frequency processing confers a potential advantage for resolving spatial information from turbulent odour plumes. It could, for instance, enable rodents to perform source separation, correctly assigning odour molecules to their respective origins, and help tackle the “olfactory cocktail party problem” (Hopfield, 1991; Rokni et al., 2014; Tootoonian et al., 2025). An olfactory system with access to rapid fluctuations allows mammals to extract distance and direction cues without physically sampling multiple locations, thereby improving efficiency in odour-driven spatial tasks.

Here, we tested whether mice can indeed extract distance-related information from the spatiotemporal structure of odour plumes. By combining wind tunnel recordings in an automated behavioural setup (Erskine et al., 2019) and an “olfactory virtual reality” system (Ackels et al., 2021), we systematically explored how temporal features at various frequencies contribute to distance discrimination. Our results show that odour fluctuations in the sub-sniff frequency range are especially informative about space, providing more salient distance cues than slower timescale averages such as mean concentration. Mice learned to distinguish odour sources placed at different distances purely based on these high-frequency cues, underscoring the temporal bandwidth of rodent olfaction. Furthermore, two-photon Ca^2+^ imaging in the OB identified a subpopulation of MTCs that exhibit distance-dependent responses, suggesting a neural correlate for this spatial sensitivity. Overall, our work demonstrates that rodents can make use of rapid temporal signatures of odour plumes for spatial perception.

## Results

### Mice can discriminate between odours originating from near and far sources

Natural odour plumes are shaped by airflow turbulence, resulting in high-frequency odour intensity fluctuations that may contain information about odour source location (Gumaste et al., 2020, p. 20; Hopfield, 1991; Liu et al., 2020). Recent work has shown that mice can access high frequency components of odour stimuli (Ackels et al., 2021; Dasgupta et al., 2022; Warner et al., 2024), and suggested this could be used by mice to gather spatial information using olfaction (Ackels et al., 2021; Bhattacharyya & Singh Bhalla, 2015; Findley et al., 2021; Liu et al., 2020). In this work, we investigated whether mice can extract and use this information to discern odour source relative distance, and how this information is represented in the OB.

To this end, we first created a naturalistic but controlled environment that was robust enough to allow for long-term, high-throughput animal training, but complex enough to mimic naturally-occurring plumes (**Fig. 1A-C**). Odour stimuli were generated with custom-built distance Odour Delivery Devices (dODDs, Methods, **Fig. 1B**) placed inside a wind tunnel, using a passive-release mechanism (liquid odour picked up by passing airflow when the dODD is opened, **Fig. S1.1, S1.2 A**,**B**). This approach allowed for odour plumes to be generated anew for every trial, thus creating highly complex temporal structures (**Fig. 1C**) under reproducible conditions that nevertheless mimic the high variability occurring under natural conditions.

**Figure 1:**
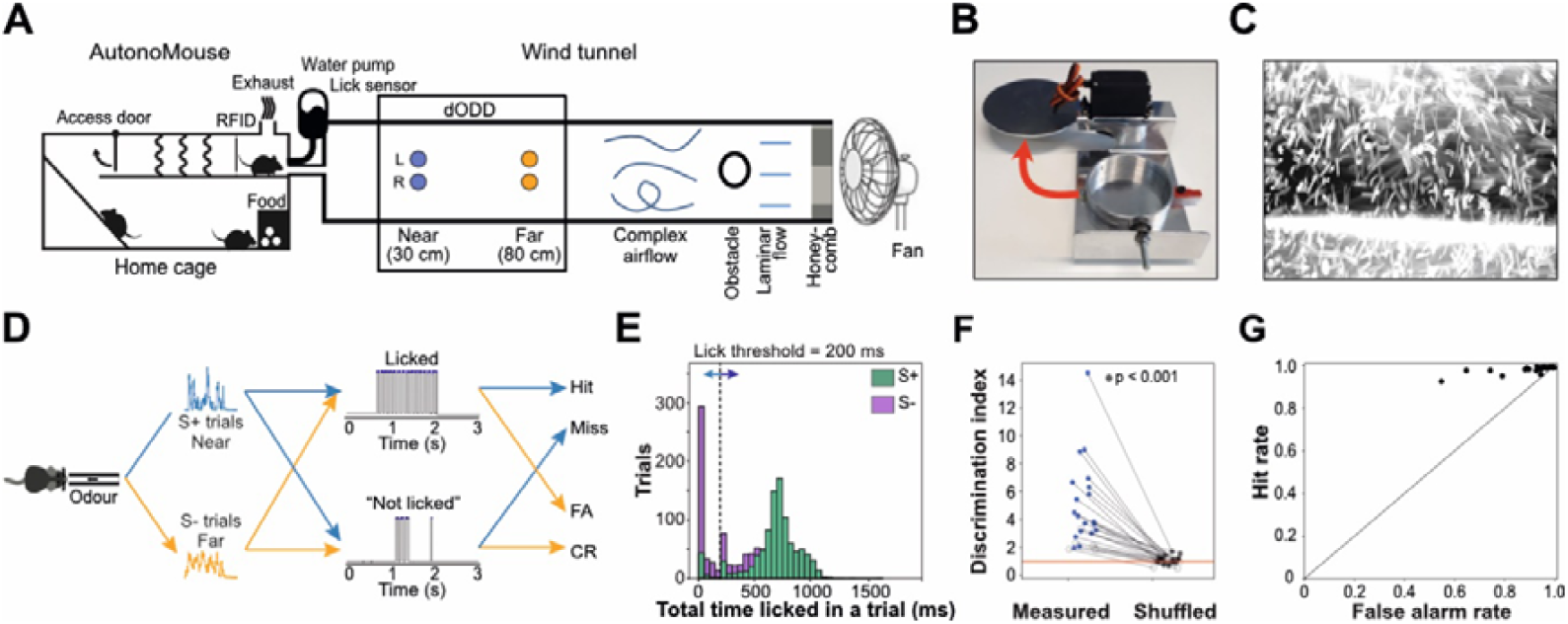
Mice can discriminate between near and far odour sources. **(A)** Complex airflow is generated inside a wind tunnel using a fan, a honeycomb structure and an obstacle. Odour sources are placed downstream of the fan and upstream of the detector (PID), forming the distance Odour Delivery Device (dODD). The dODD is formed of 8 odour boxes, 4 at each distance (near or far) from the PID, arranged symmetrically across the midline of the tunnel (Left and Right positions). Mice (n = 24) were housed in an automated operant conditioning system (AutonoMouse), where the only water source was the reward from the GNG task. **(B)** Odour presentation boxes consisting of a metal casing surrounding a glass dish. An Arduino controlled servo motor opens the lid to allow airflow to pick up odour molecules. The lid opens horizontally to the table for minimum airflow disturbance. **(C)** Complex airflow in the wind tunnel generated by a honeycomb structure and a cylindrical obstacle placed 25 cm downstream of the fan. Maximum intensity projection through a 10 s video of neutrally buoyant soap bubbles in the wind tunnel, filmed 30 cm downstream from the fan, immediately behind the obstacle. Bubbles appear elongated due to the exposure time of the camera. See also **Fig. S1.1 (D)** Schematic of task structure: Depicted is the reward structure for half of the animals (n = 13), where plumes originating from the near source (30 cm) were associated with a reward (S+ trials) and the plumes originating from the far source (80 cm) were not associated with a reward (S-trials). A lick response (top) would allow for early termination of the trial if the total time licked (shaded in blue on the lick trace) was >200 ms in a 2 s rolling window. The absence of a lick response (bottom) was defined as total time licked <200 ms in a 2 s rolling window, and the trial would continue until its maximum length was reached at 3s. These two categories of responses to the two types of trials (S+ or S-) produced the outcomes listed on the right-hand column (Hit, Miss, False Alarm, Correct Rejection). **(E)** Histograms of total time licked for an example mouse, as calculated by the AutonoMouse software to decide upon delivering a water reward, using time above threshold (dotted line) in the lick traces as shown in D (S+ in blue, S-in orange). **(F)** Discrimination index defined as the ratio of total time licked (S-/S+) for each mouse below the threshold (t = 200 ms), compared to the mean of shuffled values for that mouse. The measured value is significantly higher than shuffle (measured>shuffled in 99% of shuffles) in 19/24 mice (filled dots). **(G)** Hit rate (fraction of S+ trials licked) plotted against False alarm rate (fraction of S-trials licked) averaged across a period of 10 blocks of 100 trials each during a stable period of the experiment. Each dot is an individual mouse. A perfect block would be placed in the upper left corner, while a block with continuous licking would be placed in the upper right corner. A block where the mouse was not engaged in the task would be placed in the lower left corner. Indiscriminate licking while engaged in the task would be placed along the diagonal.

A home cage-based automated behavioural system (AutonoMouse (Erskine et al., 2019); Methods) was connected to the downwind end of the wind tunnel, allowing the animals to sample odour plumes created inside the wind tunnel (**Fig. 1A, Fig. S1.3**). This allowed us to simultaneously train a cohort of group-housed mice (n = 24) for a long period of time (up to 14 weeks), resulting in a large number of trials (4053.9 ± 660 trials/animal). Using this system, mice were trained to discriminate categories of odour plumes originating from different distances using an olfactory Go/No-Go (GNG) task paradigm (**Fig. 1A-D**). By harnessing the high number of trials that can be generated and presented to the mice, we were not limited to predetermined stimulus structures and can thus study discrimination ability in mice in a naturalistic context with ethological relevance, where the stimulus is highly variable, but also highly informative of odour source distance.

Mice were trained to respond with licking to the S+ and suppress licking to S-stimuli, where the odour source distance associated with a reward (S+) was “Near” for half of the animals, and “Far” for the other half (**Fig. S1.5**). The total duration of licking (total time licked) was calculated for each trial and used in the analysis of discrimination ability (**Fig. 1E, Fig. S1.6**). Using this measure, we found the distribution of total time licked to be different between S+ and S-trials for 16/24 mice (p < 0.05, Bonferroni-corrected Mann-Whitney U test). More specifically, mice showed a narrower distribution of lick durations during S+ trials compared to a wider distribution, often shifted to the left or with an additional peak below 200 ms for S-trials (**Fig. 1E** example mouse, more mice in **Fig. S1.6**). This suggests mice were more engaged in licking to S+ trials, compared to S-(they spent more time licking and licked more consistently over a large number of trials).

To quantify the difference between the distributions of lick durations in response to S+ or S-trials, we split the response window at a threshold of t = 200 ms and calculated a discrimination index defined as the ratio between the number of S-trials and the number of S+ trials with a lick duration smaller than the threshold t (**Fig. 1F**). A ratio >1 indicated a higher number of S-trials with lick durations shorter than t, indicating correct lick behaviour. Conversely, a ratio value <1 indicated a preference for S+ trials. For a perfectly trained mouse with no bias, the ratio value would tend towards infinity, while a ratio value close to 1 would indicate no preference (and therefore no difference between S+ and S-). To distinguish a real difference from one obtained by chance, while also taking the lick bias and the difficulty of the task into account, we compared the ratios obtained for individual mice to ratios calculated from their own responses, after shuffling the labels (**Fig. S1.6M**). All mice showed a Discrimination index >1 and the differences were significant for 19/24 mice (p < 0.001) when compared to the corresponding shuffled controls (**Fig. 1F**). Additionally, lick duration distributions remained robustly different for any threshold chosen between 50 and 600 ms (**Fig. S1.6N**,**O**). This analysis was necessary due to a clear bias for licking present in all mice. Thus, most mice displayed almost 100% accuracy in responding to S+ stimuli by licking (Hits). However, when mice did refrain from licking, they predominantly did so only in S-trials, as evidenced by the lower lick fraction recorded in S-trials compared to S+ trials (**Fig. S1.4**, fraction licked split by S+/S-), resulting in fewer False Alarms while still displaying a high Hit rate (**Fig. 1G, Fig S1.6P**). This is what allows mice to perform with above chance accuracy (**Fig. S1.4**). Overall, we conclude mice can discriminate odour sources placed at different distances based on the different lick behaviours expressed in response to S+ compared to S-.

The primary aim of the behaviour experiment described above was to determine whether mice are capable of discriminating between odour stimuli with the same odour identity, but delivered from two different locations in space, and thus arriving at the nose of the animal with different temporal structures. We found that mice were capable of this discrimination, but it remained unclear what information in the stimulus they were using to inform their responses. In order to answer this question, we further aimed to investigate what information is present in the odour plumes presented to the mouse, and what information from these plumes is accessible to the mouse olfactory system.

### What features of the odour plume allow for distance discrimination

To analyse what information is available in the plume structure that would allow distance discrimination, we recorded temporal structures of odour plumes using a photoionization detector (PID). We placed the detector in the same location where a mouse’s nose would be during a trial, and delivered odours as described above (**Fig. 2**). This was done during the same period as animal training (though not simultaneously), with the additional purpose of confirming stimulus stability throughout the training period. This large plume bank allowed us to get a handle on the large variability seen in naturalistic plumes, which is a key feature of our approach, as well as an experimental challenge. The temporal structure of recorded odour plumes was different between distances. **Figure 2B** shows example recorded plumes from a far source and near source, respectively. Additional examples (**Fig. S2.1**) further illustrate the differences between distances, as well as the variability within categories and across recorded plumes overall.

**Figure 2:**
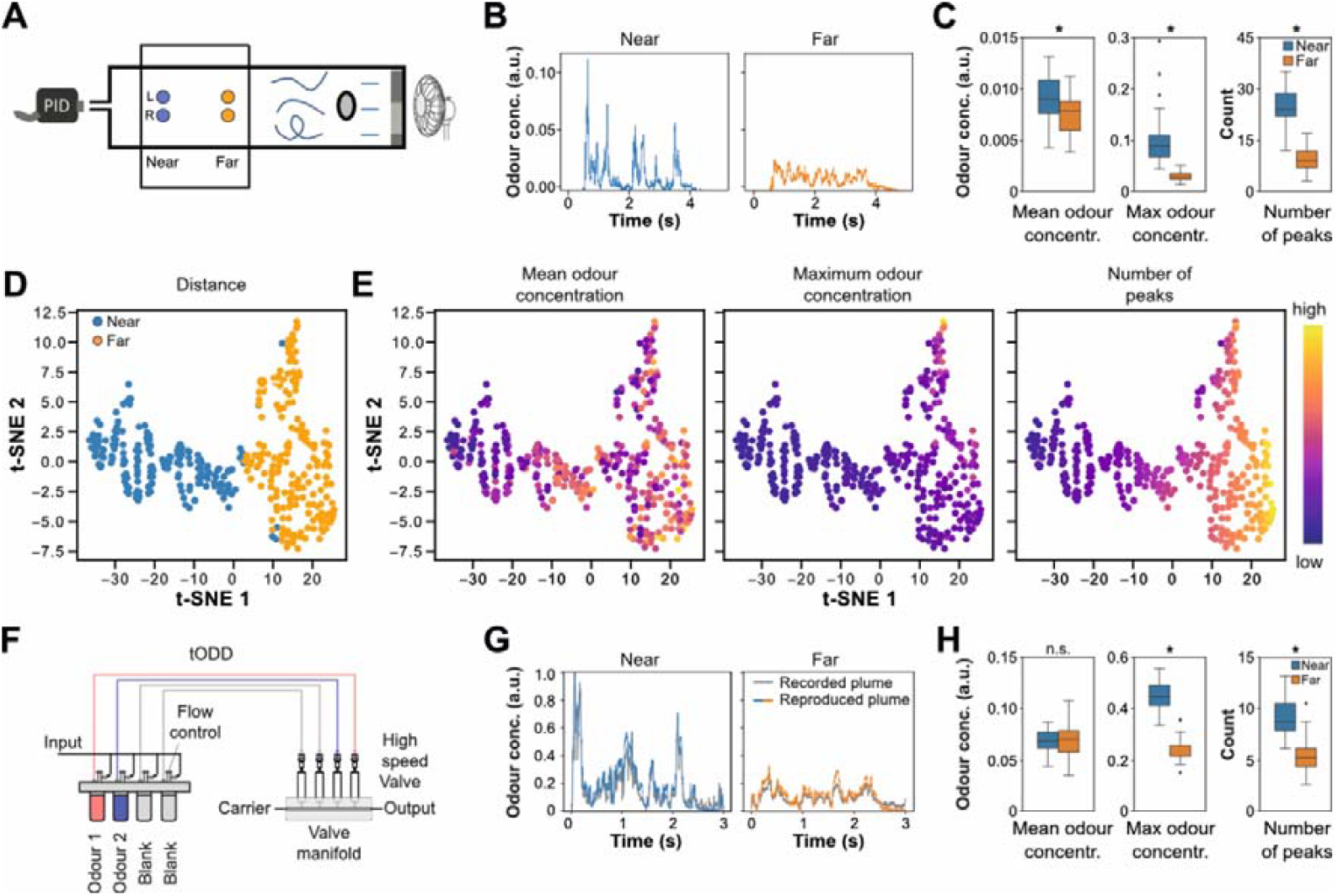
Odour plume structure varies reliably with distance and sub-sniff features are the most informative. **(A)** Schematic of the distance Odour Delivery Device (dODD) as in **Fig. 1**. Complex airflow is generated inside a wind tunnel using a fan, a honeycomb structure and an obstacle. Odour sources are placed downstream of the fan and upstream of the detector (PID) and arranged symmetrically across the midline of the tunnel (Left and Right positions). **(B)** Example plumes recorded from the Near (30 cm) and Far (80 cm) positions. **(C)** Quantification of Mean odour concentration (left), Maximum odour concentration (middle) and Number of peaks (right) for near and far plumes. Box indicates 25th-75th percentiles, thick line is median, whiskers are most extreme data points not considered outliers (see methods). **(D)** tSNE analysis of odour features (mean odour concentration, maximum odour concentration (largest peak height), number of peaks, maximum peak prominence, standard deviation, variance, kurtosis, skewness) calculated for a 5 s time window for each plume, showing clear separation by distance. Each dot is a trial (n = 360 plumes, see methods). **(E)** Same as (D) but for plume structure features. Dots are colour-coded the value of individual temporal structure features to visualise any gradients preserved in the two-dimensional space. **(F)** Schematic of temporal Olfactory Delivery Device (tODD). **(G)** Example plumes from near and far distances. Recorded plumes in grey, reproduced plumes in blue (near) or orange (far). **(H)** Same as in (C) but for plumes reproduced using tODD. Box indicates 25th-75th percentiles, thick line is median, whiskers are most extreme data points not considered outliers.

To quantify the differences in the temporal structure of odour plumes, we compared the two distance categories for eight features calculated for each plume instance (**Fig. 2B,C, Fig. S2.2, S2.3, S2.5**, Methods): mean odour concentration, maximum odour concentration (largest peak height), number of peaks, maximum peak prominence, standard deviation, variance, kurtosis, skewness (**Fig. 2C** and **Fig. S2.3** with all features). The most substantial differences were observed in temporal features of the plumes, such as number or height of peaks. These were also the features that allowed for the highest accuracy of a linear discriminant trained on individual features one at a time (**Fig. S2.4**). Moreover, combining different features allows for almost perfect classification of plumes into near and far categories (tSNE, eight features used, **Fig. 2D**) where the number of peaks is the most informative axis for the discrimination **(Fig. 2E**; see **Fig. S2.7** for complementary dimensionality reduction analysis using PCA). Thus, the temporal structure of odour plumes contains enough information to distinguish stimuli from sources at different distances, with some features allowing for higher discrimination accuracy than others.

In order to further investigate which features are most informative, we moved to a more controlled setting, in which we could readily simulate odour plumes from different distances using a high-speed odour delivery device: the temporal Odour Delivery Device (tODD (Ackels et al., 2021); **Fig. 2F**). Using this device, we were able to reproduce the recorded structures described above with high fidelity (**Fig. S2.8, S2.9, S2.10**). The reproduction strategy retained the discriminability between plume categories based on individual features by capturing the differences and variability in features observed in recorded plumes (**Fig. 2G,H, S2.11, S2.12**).

By capturing and replicating the spatiotemporal attributes that differentiate near from far odour sources, we have shown that temporal features can reliably classify plume distance categories. Precise playback of these recorded plumes preserves variability across individual stimuli. This sets the stage for tightly controlled physiology experiments under head-fixation, to link the observed discriminability in behavioural tasks to neural responses in the OB.

### Odour source distance is encoded in a subset of OB projection neurons

We have so far shown that mice can discriminate between odour sources placed at different distances, and that temporal information present in odour plumes would allow for this discrimination. We next examined whether these high-frequency cues are actually accessible to OB neurons. Historically, the brief, rapidly fluctuating concentration peaks found in turbulent plumes were assumed to exceed the temporal resolution of mammalian olfaction. However, recent work suggests otherwise (Ackels et al., 2021; Dasgupta et al., 2022; Warner et al., 2024; Wu et al., 2024). To investigate whether the OB can indeed extract distance-related information from naturalistic plume structures, we combined a method for reliably recreating plume dynamics with two-photon calcium imaging of dorsal OB projection neurons (MTCs) in head-fixed, anaesthetised mice (Tbet-cre:GCaMP6f, n=3), while simultaneously monitoring respiration (**Fig. 3A-C**). Rather than repeatedly presenting a single plume signature, we delivered 40 distinct “near” plumes and 40 distinct “far” plumes (**Fig. 3D**, outer columns), capturing a broad range of temporal features (**Fig. 2G-H**). This allowed us to assess whether information about distance can be extracted by the OB from highly variable, naturalistic stimuli. Additionally, the large number of distinct plumes, and the resulting wide range of different features and feature combinations covered by our stimulus space (**Fig. 2G-H**), allowed us to further probe into how different features of naturalistic odour plumes are represented in the OB.

**Figure 3:**
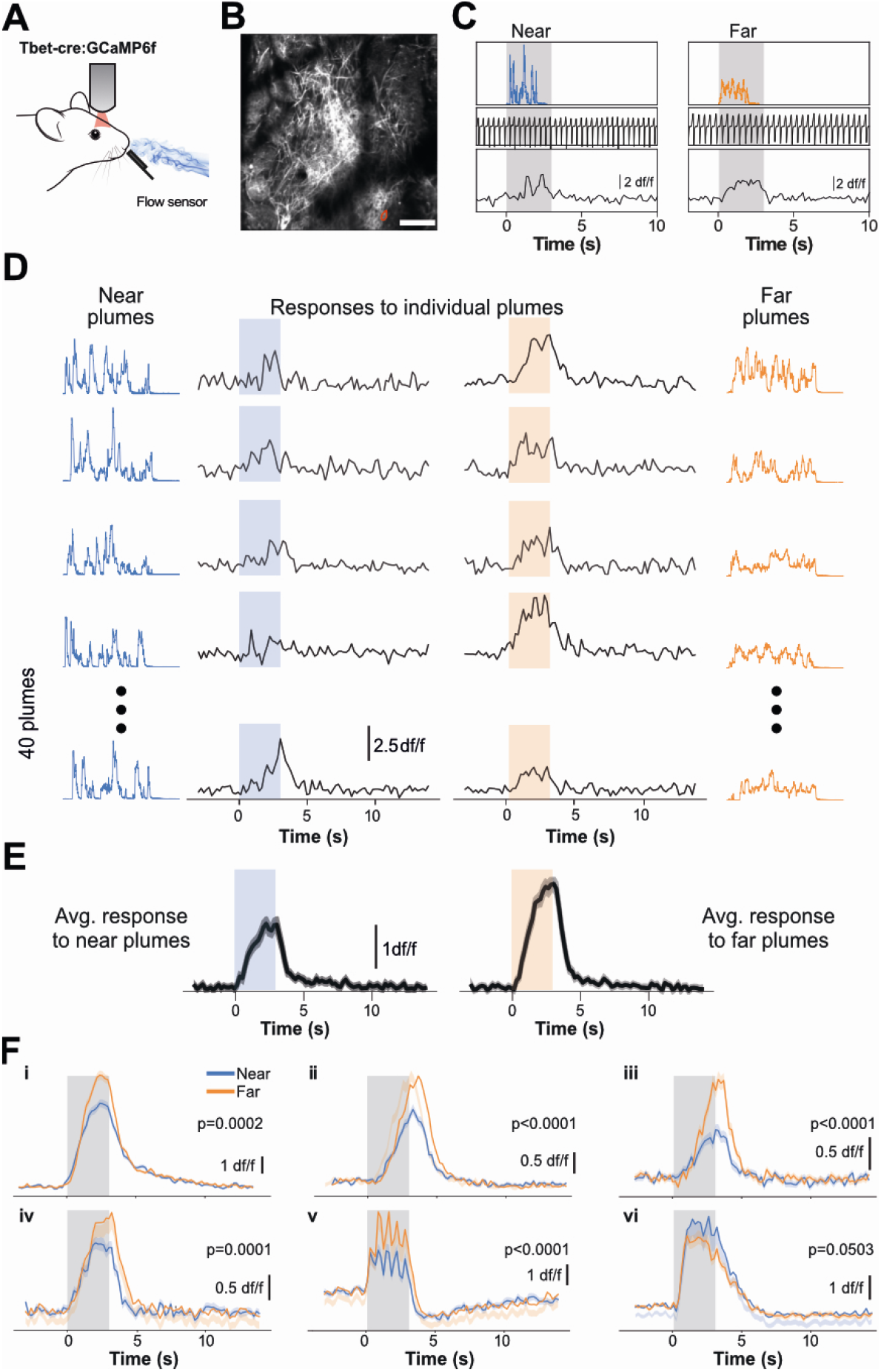
Responses of MTCs to temporally structured odour plumes recorded from different distances. **(A)** Schematic of the two-photon imaging approach. **(B)** GCaMP6f fluorescence from mitral and tufted cells (maximum projection of 8,000 frames). Responses from ROI marked in red are shown in (C). Scale bar, 100 μm **(C)** Example single trials in response to an odour stimulus from a near (30 cm) or far (80 cm) virtual odour source (top row). Respiration traces (inhalation pointing upwards) from the same trials (middle row). Example fluorescence traces (df/f) from the red circled ROI in (B). Shaded area represents 3 s odour stimulation window. **(D)** Individual near and far plumes (outer columns) and corresponding calcium responses (inner columns) of one ROI. **(E)** Averaged responses to all near and far plumes (n = 40 for each condition, mean ± s.e.m.). **(F)** Average fluorescence traces (mean ± s.e.m., n = 160 trials) for 6 example distance sensitive ROIs (near: blue, far: orange; ROIs i-iv: EB, v-vi: 2H).

To assess whether the OB is able to discriminate between plume structures originating from different distances, we recorded neuronal activity (as expressed by a change in GCaMP6f fluorescence, **Fig. 3D**, inner columns) in response to 40 different plume structures for each distance category (**Fig. 3D**, outer columns). Across all trials (∼90 per distance, two odours; ethyl butyrate and 2-heptanone), we identified a small subset (13 out of 1062 cell-odour pairs; 1.2%) that responded differently to near versus far plumes (**Fig. 3E**). These “distance-sensitive” cells showed stronger responses to far stimuli in the majority of cases (**Fig. 3F**), and each cell’s sensitivity was typically specific to one odour (**Fig. S3.1, S3.2**). In addition, a subset of plumes was presented multiple times to capture response variability (**Fig. S3.2**). An overview of responses of all ROIs to all stimuli (**Fig. S3.3**) and to intermediate distances (**Fig. S3.4**) and to a subset of plumes that were presented multiple times (**Fig S3.5)** illustrates the variety in their response strengths and kinetics. This indicates that, despite the broader heterogeneity in temporal plume features, a subset of MTCs can encode distance-related information.

This finding, while involving only a small number of MTCs, suggests that distance-specific signals may be accessible at the population level. It therefore led us to the question of whether these neural responses can be used to predict an odour source’s distance? In the following section, we address this by examining whether the patterns of OB activity can serve as a readout of plume distance.

### Odour plume distance can be predicted from MTC responses

We next assessed whether distance-dependent differences in OB activity extend beyond the small number of clearly distance-sensitive MTCs (**Fig. 3**). To visualise the responses underlying this classification, we plotted the average calcium signals in response to near and far plumes over time (**Fig. 4A**, left and middle). The difference map (**Fig. 4A**, right) highlights that a subset of cell-odour pairs exhibited higher activity in response to far plumes (blue) or near plumes (red). A linear classifier was trained on the population responses of all recorded cell-odour pairs to determine whether the stimulus originated from a near or far source, based on the ensemble activity (**Fig. 4B**). Classifier performance steadily increased with the number of cell-odour pairs included (purple line, maximum performance: 73.0 ± 9.0%), substantially exceeding the shuffled control accuracy (black line). This result indicates that while only a fraction of MTCs responded significantly differentially to distance, activity patterns across the population of cells carry significant information about plume origin. Further classification analyses for both odours individually (**Fig. S4.1, S4.2**) and a pairwise comparison between further intermediate odour source distances (**Fig. S4.3**) can be found in the supplement.

**Figure 4:**
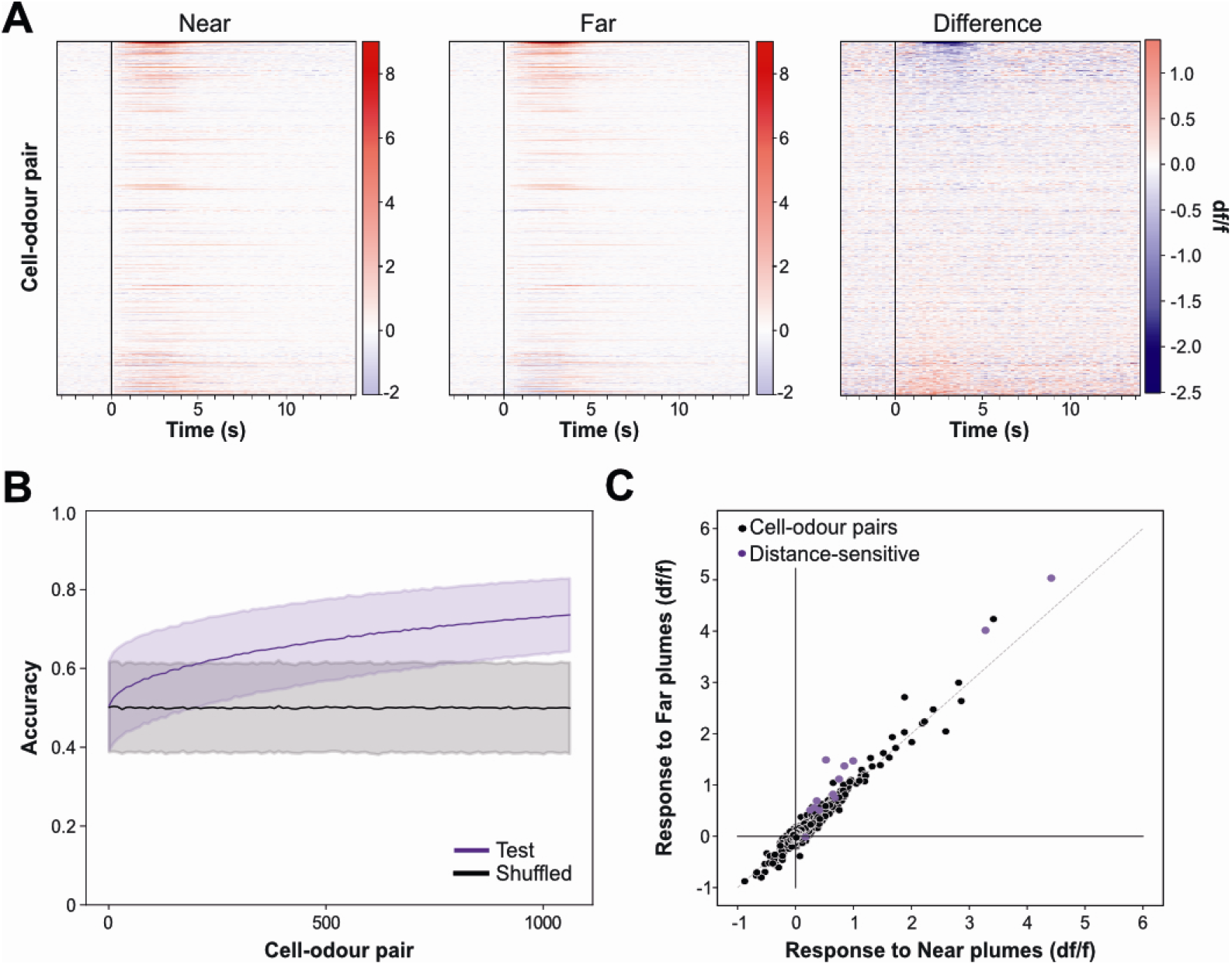
Population activity and linear discriminant analysis of MTC responses. **(A)** Calcium transients as colour maps for averaged responses to near (left) and far (middle) plumes and for the difference between the two (right) for all cell-odour pairs (n = 1062). **(B)** Accuracy of linear classifier trained on a 5 s response window (purple; black, shuffle control) to near versus far plumes (n = up to 1062 cell-odour pairs from 3 mice; mean ± s.d. of 50000 repetitions). **(C)** Responses to near vs responses to far plumes plotted as mean df/f response for all trials from those distances. Distance-sensitive cells shown in purple. Cells above diagonal respond more strongly to far plumes while cells below diagonal respond more strongly to near plumes.

When comparing mean responses of individual cells to far versus near conditions (**Fig. 4C**), the majority of responses showed similar amplitude for both plume types, but a subset (purple) fell above the unity line, showing stronger responses towards far stimuli. When we further segregated these cells’ activity by odour identity, it became clear that distance sensitivity was associated with the odour that elicited stronger responses (**Fig. S4.4, S4.5**). Overall, these results demonstrate that even modest differences in MTC response strength allow above-chance prediction of odour source distance from population activity.

In the final part of our results, we will examine which temporal features of naturalistic plumes underlie this distance-dependent coding, providing a direct link between the neural representation in the OB and the animal’s ability to discriminate odour sources at varying distances.

### Stimulus features informing distance-sensitivity

We first demonstrated that mice can discriminate between odour sources placed at different distances. Additionally, we showed distance-related information is also discernible from the activity of OB projection neurons. Next, we aimed to understand which features of the stimulus inform the OB (and consequently the animal) about distance. To this end, we took advantage of the wide range of plume structures we presented in the head-fixed setting. As described above (and Methods), we presented 40 different plume structures for each distance category, with each plume not only informing about distance, but also expressing many temporal features, in different combinations. Thus, this experimental design allowed us to analyse cell responses not only in terms of preference for a particular distance, but also preference for specific features of the odour plume. To further leverage diversity in the temporal features space, we introduced two intermediate source distances (40 cm and 60 cm) for a total of 160 different plume structures presented to the mouse (**Fig. 5A**). For each plume we defined extracted values for an array of different features. This allowed us to move beyond simple near vs far comparison (broad categories) and explore how the OB encodes specific temporal features across and within categories. To support consistent plume presentation and alignment, respiration was recorded during the experiments and remained stable across mice and conditions at ∼2 Hz (**Fig. S5.1**). Stimuli were aligned to inhalation onset, ensuring consistent timing across trials. We also verified that total and inhaled odour concentrations were broadly matched across distances (**Fig. S5.2**), ensuring that observed neural differences were not simply driven by concentration differences but solely by temporal plume structure. When examining odour exposure on a per-sniff basis, we found that the amount of odour inhaled during the first sniff was comparable across distances, while differences in the inhaled odour amount between sniffs emerged from the second sniff onwards, particularly for near versus far plumes (**Fig. S5.3**).

**Figure 5:**
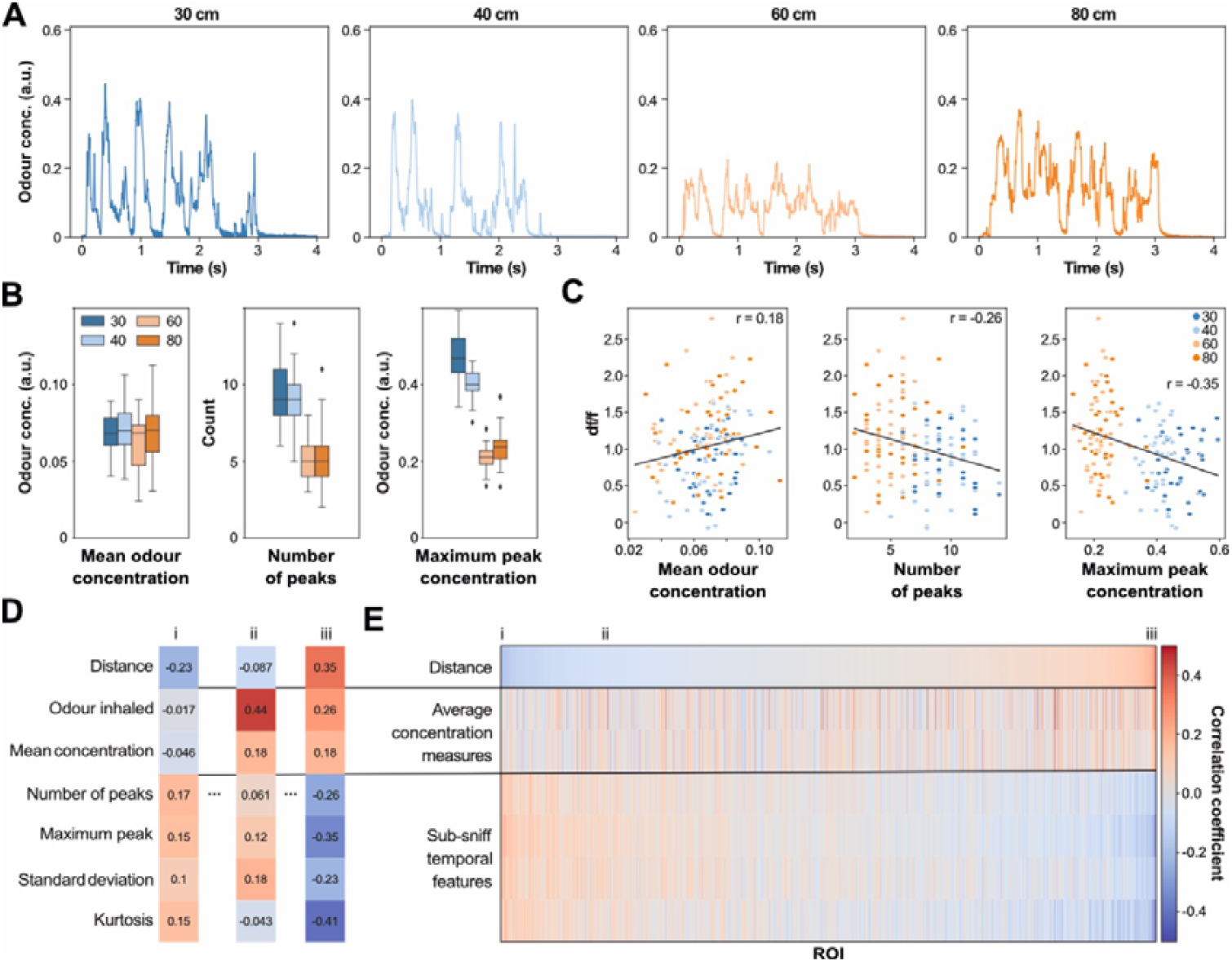
Features of the temporal structure of odour plumes involved in distance discrimination. **(A)** Example plumes recorded from four different distances (30 cm, 40 cm, 60 cm and 80 cm). **(B)** Quantification of mean odour concentration (left), number of peaks (middle) and maximum odour concentration (right) for all plumes from four distances. Box indicates 25th-75th percentiles, thick line is median, whiskers are most extreme data points not considered outliers (see methods). **(C)** Response of an example distance-sensitive ROI (df/f averaged over the 5 s response window) plotted against individual features of the odour stimulus (x-axis). Each dot is a trial (n = 160 trials), colour-coded by distance (80 cm orange, 60 cm pale orange, 40 cm pale blue, 30 cm blue). **(D)** Correlation coefficients to distance and other stimulus features for three ROIs (i-iii). **(E)** Heatmap of correlation coefficients as calculated in C for all cells (columns, n = 531 ROIs from 3 animals), sorted by their correlation to distance (top row). Labels i, ii, iii correspond to ROIs shown in (D).

To investigate which stimulus properties drive the responses of distance-sensitive cells, we leveraged the substantial variability in temporal features within each distance category (**Fig. 5B**). Specifically, we examined the correlation between trial-by-trial response amplitudes and individual plume features. **Figure 5C** illustrates the method for a representative distance-sensitive ROI: the calcium response is plotted against three plume descriptors (mean odour concentration, number of peaks and maximum peak concentration). This approach revealed cells whose activity is strongly correlated to certain plume features. For example, the response of Cell ii in **Fig. 5D** appears to be closely linked to the total odour inhaled. Notably, for distance-sensitive cells the strongest negative correlations appeared with the plume features that best indicate source distance. These features operate at sub-sniff timescales, such as the number of peaks and the maximum peak concentration (**Fig. 5C,D**). When we sorted all recorded cells by their correlation with distance, we observed that correlation between response and distance could be inversely mapped to the correlation between response and sub-sniff temporal features (**Fig. 5E**, bottom part of the heatmap). In contrast, no pattern emerged between correlation with distance and average correlation measures (**Fig. 5E**).

Collectively, these findings underscore that sub-sniff temporal dynamics, rather than simple concentration averages, serve as key indicators of source distance for OB neurons. This provides critical insight into how rodents might use the complex temporal dynamics of odour plumes under natural conditions.

## Discussion

Rodents depend on olfactory cues to navigate their surroundings, making the ability to discern whether an odour source is nearby or distant critical for survival decisions such as foraging or predator avoidance. To investigate how rodents might perform such discrimination, we developed an experimental setup that reliably produces temporally complex odour plumes from different distances, striking a balance between naturalistic complexity and experimental control. With high-throughput behavioural assays and *in vivo* physiological recordings, we examined whether mice can discriminate the distance of different odour sources and how distance-related temporal plume features are encoded within the OB.

Our results demonstrate that mice can discriminate odour sources placed at different spatial distances, showing for the first time that information within odour plumes provides sufficient cues for distance discrimination. We observed that plume features at sub-sniff timescales, rather than broader measures such as mean odour concentration, carried more information to discriminate odour source distances. Particularly the number of peaks and maximum peak concentration provided higher accuracy in distinguishing between near and far sources. This aligns with odour bout count and stimulus intermittency measures proposed in previous research (Gumaste et al., 2024; Schmuker et al., 2016). We conducted plume feature analysis at high sampling frequencies (1 kHz) which likely exceeds the temporal resolution of mouse olfactory perception (approximately 40 Hz; (Ackels et al., 2021)) and thereby potentially inflates peak counts and magnitude values. Future studies should systematically test plume feature reliability and informativeness at various sampling frequencies to align with rodent sensory capabilities (Boero et al., 2025; Gumaste et al., 2024; Lewis et al., 2024). The distances we chose for our experiments provided clear and discriminable statistical differences in plume features, effectively capturing the natural variability rodents might encounter in their environments (Reddy et al., 2022; Rigolli et al., 2022). Interestingly, we observed that the four distances studied grouped into two distinct categories, suggesting a near versus far categorisation as opposed to a gradient mapping of source distance. This categorical distinction may reflect an intrinsic boundary in rodent olfactory perception (Liu et al., 2020; Rigolli et al., 2022). Yet, whether this categorization generalises beyond our experimental settings remains to be determined. Expanding the range of distances and systematically varying environmental parameters, such as airflow dynamics and obstacle configuration, would clarify how robustly specific plume features inform distance discrimination across diverse contexts.

Our findings show mice consistently displayed differential licking responses to odour sources at varying distances, although achieving consistently high discrimination accuracy proved challenging. Several factors likely contributed to this moderate performance. Notably, the inherent asymmetry of the Go/No-Go task, where an impulsive action (licking) is contrasted against inhibitory control (withholding licking), might have influenced overall accuracy (Carandini & Churchland, 2013). Detailed analysis of licking patterns revealed that mice were more consistently responsive during rewarded (S+) trials, suggesting clear reward anticipation, whereas variability in responses during non-rewarded (S−) trials might reflect fluctuations in motivation rather than genuine perceptual errors (Berditchevskaia et al., 2016). Additionally, efforts to mitigate predictable differences in odour arrival times between distances, by incorporating variable onset delays and flexible trial termination, inadvertently intensified task asymmetry. Although designed to maintain engagement, these adjustments allowed mice to conclude rewarded trials prematurely while requiring them to endure the full duration of non-rewarded trials for a correct response. Consequently, mice may have strategically licked simply to reduce trial durations, potentially decoupling their licking decisions from actual stimulus discrimination. Taking these behavioural and task-design considerations into account, it seems likely that mice possess greater discrimination capability than was observed under our current experimental conditions. Refining the training paradigm, for instance by implementing a two-alternative forced-choice (2AFC) task (Bitzenhofer et al., 2022; Boero et al., 2025; Nakayama et al., 2022) to mitigate motivational biases, may help shed further light onto the limits of sensory discrimination capabilities.

A key question we asked was: How is odour source distance represented in the OB? To ensure naturalistic variability of temporal patterns, we exposed animals to a wide range of plume structures recorded from near and far sources. When decoding odour source distance from the population of MTC calcium responses, we found that accuracy exceeded behavioural performance of mice proficient in the discrimination task. This suggests that distance information carried by odour plumes could, in principle, support even more precise discrimination. Strikingly, a small subset of individual MTCs exhibited clear differential responses to near versus far plume sources, with most distance-selective MTCs responding more strongly to far than to near stimuli in most cases. Notably, this tuning was already present in naïve animals that had never been trained to judge source distance. To understand what might be driving this selectivity, we presented a large and diverse set of plume structures, including intermediate distances (40 cm and 60 cm). When correlating neural responses in a trial-by-trial fashion with specific stimulus features, we observed that their activity tended to correlate inversely with sub-sniff plume features most predictive of source distance. Furthermore, when sorting cells by their correlation with distance, we found that their response tuning could be almost perfectly mapped to their tuning to these fast temporal features. In contrast, correlations with broader, average plume statistics did not show such a pattern. These findings suggest that fast fluctuations in odour intensity, rather than slow integrative cues, may play a central role in how OB neurons encode spatial information. We acknowledge, however, that this observation is correlative, and future work is needed to determine whether these features are used directly by the OB circuitry to support behavioural discrimination.

What cellular or circuit mechanisms could account for these distance-dependent responses? One possibility is that MTCs are not “counting peaks” per se, but instead integrate the constancy of odour concentration (Boero et al., 2025; Lewis et al., 2024). Less-peaky, more continuous plumes (typical of far sources) could drive a larger, temporally summated input from ORNs than highly intermittent near plumes (Lewis et al., 2024). Since ORNs of a given receptor type are scattered across the epithelium before converging onto the same glomerulus (Malnic et al., 1999; Mombaerts et al., 1996), pockets of odorised air will likely recruit partially non-overlapping ORN ensembles over time. A smoother plume might therefore activate a larger part of that ensemble, whereas a smoother plume with low intermittency may engage fewer ORNs within brief time windows. Additional temporal summation could occur at the level of individual ORNs or of their postsynaptic MTCs, as ORN spike fidelity drops at high stimulation frequencies or high concentrations (Ghatpande & Reisert, 2011). Taken together, this could transform fine-scale plume fluctuations into population codes that emphasise the more stable temporal signature of distant sources. Consistent with this view, the distance-sensitive MTCs we identified were generally more narrowly tuned to odour identity. This raises the testable idea that the most reliable responders within a glomerular column jointly encode both chemical identity and temporally derived spatial information (Arneodo et al., 2018).

Methodologically, our calcium imaging approach is well suited for identifying and targeting specific neurons, but its temporal resolution limits capturing millisecond-scale plume dynamics. Future work using high-density electrophysiology, faster GCaMP variants, or genetically encoded voltage indicators will be essential to resolve the precise timing of ORN and MTC firing relative to fine temporal plume features (Lewis et al., 2024; Storace et al., 2015). Combining such recordings with selective activation of a single, genetically labelled glomerulus (Arneodo et al., 2018; Schwarz et al., 2018), and multiplexed imaging of both ORN input and MTC output (Martelli & Storace, 2021; Storace & Cohen, 2017) could reveal how lateral inhibition, sister-cell diversity (Zhang et al., 2025), and molecular specialisations together give rise to distance-sensitive signalling in the OB.

Our results show that mice can discriminate the distance of an odour source by tracking rapid, sub-sniff changes in a plume, and that a subset of MTCs in the OB reflects this information. By tying together plume measurements, behaviour, and single-cell activity, we add a strong link between the physics of natural odours, animal behaviour and neural coding. Future studies could further clarify which temporal plume features rodents utilise for spatial discrimination in different contexts by employing an olfactory virtual reality coupled with precise plume control, or even in unrestrained mice. Such experiments will help to pinpoint precisely, how specific olfactory bulb circuits transform natural chemical signals into reliable cues for navigation.

## Supporting information

Main figures (high resolution)

Supplementary Figures (high resolution)

## Methods

### Behaviour experiments

#### Animals

All animal procedures performed in this study were approved by the UK government (Home Office PPL number PA2F6DA12) and by Institutional Animal Welfare Ethical Review Panel. All mice used were C57/Bl6 males (n = 24 mice), evenly distributed across the different groups. Long-term group housing precluded us from using mixed-sex cohorts.

All mice in a cohort (n = 24) were group-housed from weaning (21 days of age) to avoid disruption of social hierarchy and aggression later in the experiment (Van Loo et al., 2001; Van Loo et al., 2003). They were housed into a suitably large cage, with chew blocks, a variety of houses and hiding places and multiple sources of water and food, until ready to be implanted with the RFID. All mice were kept on a 12h light cycle with the light phase between 7:00 and 19:00. The change in light occurred gradually over a period of 15 minutes. Room temperature and humidity were checked daily and kept around 21°C and 50% respectively. The group cage (and later the AutonoMouse cage) was the only cage in the room, to avoid conspecific odours disturbing the social hierarchy and causing aggression within the cage.

#### RFID implantation

The RFID implant surgery occurred around 6 weeks of age, as previously described (Erskine et al., 2019). Briefly, Mice were anaesthetised under isoflurane (induction: 5% in O_2_ 2 l/min, maintenance: 2%) and placed on a heat pad for maintenance of body temperature during the surgery. The fur around the base of the neck and scruff was shaved away and the skin cleaned with chlorhexidine (1%) and then dried with a sterile swab. A pre-sterilised needle (IM-200, RFID Systems Ltd., Yorkshire, UK) containing an RFID chip (ID-100B, RFID Systems Ltd., Yorkshire, UK, in Chapter 3, ID-162B/1.4, RFID Systems Ltd., Yorkshire, UK in Chapter 5) was then loaded onto a plunger and inserted into the loose skin at the base of the neck, facing towards the tail of the mouse. The plunger was used to push the chip out of the needle before removing the needle, leaving the RFID chip implanted under the skin. Forceps were then used to pinch shut the incision made by the needle and medical superglue (Vetbond, 3M) was applied to seal the wound. Animals were returned to an individual cage for 10 minutes following the surgery to recover from anaesthesia and for the superglue to dry. Once the righting reflex was regained and the wound was confirmed as properly sealed the mouse was returned to the group cage with its. Following at least 5 days recovery from the implant surgery, the mice were transferred to the AutonoMouse system and kept for up to 18 months.

#### Automated Behaviour setup

In AutonoMouse, groups of mice (up to 24) implanted with an RFID chip are housed in a common home cage (for detailed description see Erskine et al., 2019; Ackels et al., 2021). Within the common home cage of AutonoMouse, mice have free access to food, social interaction and environmental enrichment (including nesting material, chew blocks, climbing frames, cardboard tubes, cardboard and red plastic houses). Water is not freely available in the system, but can be gained at any time by completion of an operant conditioning Go/No-Go task. To access these behavioural tasks, mice must leave the home cage and enter a behavioural area. This behavioural area contains the odour port and a lick port through which water rewards can be released. The lick port is also connected to a lick sensor, which registers the animal’s response (time spent in contact with the sensor) in response to the task stimuli. As animals can only gain their daily water intake by completing behavioural tasks, mice are motivated to complete long sequences of trials without manual water restriction. However, there is no limit on how many times mice can trigger a trial, or how much time they can spend in the odour port, which could add additional constraints on using timeouts for negative reinforcements.

#### Acclimatisation to the AutonoMouse system

Animals were weighed and their weight recorded on the day they were transferred to the AutonoMouse and then every day for 2 weeks.

The first phase of the behavioural task assigned to all animals was a ‘pre-training’ phase, designed to train animals to reliably gain their water intake from the behavioural port. This phase was itself split into multiple tasks. At first water was delivered as soon as an animal was detected in the behaviour port (20 trials). Next, water was delivered only if the mouse licked at least once after activating the behaviour port (50 trials). Following from that, the percentage of total trial time (2 s) that the animal must lick to gain a water reward was increased in 50 ms increments up to 10% of trial length, or 200 ms (50 trials per step). Each water reward was initially 15 µl. This was adjusted to 10-30 µl depending on animal performance (to ensure all mice performed roughly the same number of trials per day).

For the first two weeks of any AutonoMouse experiment, animal weights were checked daily to ensure health status of the cohort. After two weeks, weight was checked more infrequently (once a week) but total trials performed were monitored daily to ensure animals had acquired sufficient water rewards (e.g. by performing >100 trials, 50% rewarded) in the last 24 hours. Any animal not meeting this criterion or consistently dropping in weight (more than 2 days in a row) was isolated in the behaviour port and manually given water rewards from the lick port. All animals successfully learnt to acquire water by engaging the task.

Once reliable licking was achieved, the S+ stimulus was introduced. This often caused a drop in lick reliability, but mice quickly got accustomed to the new stimuli. Once lick reliability was re-established, the mice were progressed to the first stage of the Go/No-Go odour discrimination task (EB vs IAA).

#### Distance discrimination stimuli

To probe whether mice could discriminate between odours originating from two different sources, odours were placed at different distances to the mouse in the wind tunnel. All tasks followed a standard Go/No-Go training paradigm, where one distance was associated with a water reward (S+ trials), while the other distance was not (S-trials), and licking during these trials would trigger a time-out of 7 s. Reward was reversed for roughly half the experimental group (13 mice trained to lick for Near stimuli, 11 mice trained to lick for Far stimuli). Before being introduced to the distance discrimination tasks, mice underwent a series of pre-training stages that gradually increased in difficulty:

##### Stage A - Introduction to the AutonoMouse system – 1-7 days

Briefly, animals were first trained to lick the waterspout for a certain amount of time (the lick threshold, usually 10% of total trial length, or 200 ms for trials of variable length) in the presence of an odour. Further reinforcement was offered for animals performing fewer than 200 trials/day, to ensure they could successfully use the system to meet their daily water requirement.

##### Stage B - Odour discrimination – 3 weeks

Next followed a simple odour discrimination task (EB vs IAA) where animals learnt to lick to the Go cue and refrain from licking in response to the No Go cue. Half of the animals trained to lick in response to EB, and half in response to IAA. Once stable performance above 80% was achieved, the onset of the odour delivery was varied to train the mice to wait for odour arrival, in preparation for the variable odour arrival of the plumes from the wind tunnel in the distance discrimination stage. The delay was chosen to be between 100-950 ms, randomised for each trial and starting with shorter delays before progressing to the full range of delays. A switch valve control was performed at this stage, by introducing two new valves that had not been used before in the task. This assessed whether animals were using odour information and no other environmental cues.

##### Stage C - Valve distance discrimination – introduction to the wind tunnel – 4 weeks

Next, the animals performed the same odour discrimination (EB vs IAA), but this time delivered from the wind tunnel. EB was delivered from the distance odour delivery devices, forming odour plumes, while IAA was delivered from a pressurised airflow valve. The concentration of IAA was decreased in small steps (5% per day starting from 100%) over 4 weeks, such that by the end of this stage the animals were effectively performing a distance discrimination task between the near EB source(s) and the far EB source(s). For roughly half of all animals, the near source was S+ (rewarded) and the far source was S-(unrewarded); in the other half of the group, this reward valence was reversed.

##### Stage D - Distance discrimination task – 4 weeks

Finally, mice were then progressed to the next stage of the experiment, the full distance discrimination task. This was split across 4 experimental stages (27 days excluding breaks, or 3981 ± 550 trials) with short pauses between stages for equipment maintenance (10 days between 2 and 3, and 3 days between 3 and 4). Additionally, the experiment was paused for ∼30 min every 12h to replenish the odours, shuffle the spatial arrangement of the dODD, and replace any servo motors that were at risk of failure due to overheating.

The algorithm for generating distance discrimination trials was as follows:

1. Choose whether the stimulus will be Near or Far.

2. Randomly select an onset delay of 100-950 ms.

3. Randomly select an odour dODD position (Left or Right) from the corresponding distance.

4. Randomly select a blank dODD position (Left or Right) from opposite distance.

5. Randomly select two mineral oil valves from the tODD to create a blank plume and anti-plume structure, to create a noise decoy, using a 3s long plume template from the distance plume bank as described elsewhere.

6. Trigger the trial to have valves in (5) open from time 0 until the trial is terminated, and the 2 dODD positions open from time 0 + onset delay decided in (2) + 2 s.

### Data analysis

AutonoMouse data was first extracted into MATLAB using the corresponding AutonoMouse software feature (Erskine et al., 2019, https://github.com/RoboDoig/autonomouse-control), extracting all task parameters and lick data as total time licked (sum of all “on” times) and individual lick times (times of threshold crossing). Full lick sensor traces were also saved as separate files sampled at 20 kHz for future reference. First analysis steps were performed in MATLAB using custom scripts and further analysis was performed using Python 3 in Jupyter Lab.

### Imaging experiments

#### Animals

Three female mice aged 20 weeks expressing the calcium indicator GCaMP6f in MTCs were used for this experiment. The mice were Tbet-Cre (Haddad et al., 2013) crossed with GCaMP6f reporter line Ai95(RCL-GCaMP6f)-D (Madisen et al., 2015).

#### Surgical procedure

Prior to surgery all utilised surfaces and apparatus were sterilised with 1% trigene. Mice were anaesthetised using a mixture of fentanyl/midazolam/medetomidine (0.05, 5, 0.5 mg/kg respectively). Depth of anaesthesia was monitored throughout the procedure by testing the toe-pinch reflex. The fur over the skull and at the base of the neck was shaved away and the skin cleaned with 1% chlorhexidine scrub. Mice were then placed on a thermoregulator (DC Temperature Controller, FHC, ME USA) heat pad controlled by a temperature probe inserted rectally. While on the heat pad, the head of the animal was held in place with a set of ear bars. The scalp was incised and pulled away from the skull with four arterial clamps at each corner of the incision. A custom head-fixation implant was attached to the base of the skull with medical super glue (Vetbond, 3M, Maplewood MN, USA) such that its most anterior point rested approximately 0.5 mm posterior to the bregma line. Dental cement (Paladur, Heraeus Kulzer GmbH, Hanau, Germany; Simplex Rapid Liquid, Associated Dental Products Ltd., Swindon, UK) was then applied around the edges of the implant to ensure firm adhesion to the skull. A craniotomy over the left olfactory bulb (approximately 2 × 2 mm) was made with a dental drill (Success 40, Osada, Tokyo, Japan) and then immersed in ACSF (NaCl (125 mM), KCl (5 mM), HEPES (10 mM), pH adjusted to 7.4 with NaOH, MgSO4.7H2O (2 mM), CaCl2.2H2O (2 mM), glucose (10 mM)) before removing the skull with forceps. The dura was then peeled back using fine forceps. A layer of 2% low-melt agarose diluted in ACSF was applied over the exposed brain surface before placing a glass window cut from a cover slip (borosilicate glass #1 thickness [150 μm]) using a diamond scalpel (Sigma-Aldrich) over the craniotomy. The edges of the window were then glued with medical super glue (Vetbond, 3M, Maplewood MN, USA) to the skull. Following surgery, mice were placed in a custom head-fixation apparatus and transferred to a two-photon microscope rig along with the heat pad.

#### Imaging parameters

The microscope (Scientifica Multiphoton VivoScope) was coupled with a MaiTai DeepSee laser (Spectra Physics, Santa Clara, CA) tuned to 940 nm (∼30 mW average power on the sample) for imaging. Images (512 × 512 pixels, field of view 550 × 550 μm) were acquired with a resonant scanner at a frame rate of 30 Hz using a 16 × 0.8 NA water-immersion objective (Nikon). Using a piezo motor (PI Instruments, UK) connected to the objective, a volume of ∼300 μm was divided into 6 planes resulting in an effective volume repetition rate of ∼5 Hz. The odour port was adjusted to approximately 0.5 cm away from the ipsilateral nostril to the imaging window, and a flow sensor (A3100, Honeywell, NC, USA) was placed to the contralateral nostril for continuous respiration recording and digitised with a Power 1401 ADC board (CED, Cambridge, UK).

#### Odour stimuli

For this experiment a total of 752 trials were planned to be presented for each mouse. The plumes used were recorded in the wind tunnel during the same session and were picked from the centre of the distributions of plume features as shown in Fig. S2.3. We presented 40 plumes for each of the 4 distances using 2 main odours (EB and 2H), resulting in 320 “core” trials that were randomised before being added to the trial bank. The next 384 trials consisted of the same core 320 trials, together with an additional 8 more repeats of one plume/distance/odour, which were again independently randomised and added to the trial bank. Finally, the last 48 trials were 3 repeats of one plume/distance/odour for an additional 4 odours, also randomised. This trial order was then presented to each mouse, in blocks of 40 trials, in order to facilitate data acquisition. Odour delivery was triggered by inhalation using a threshold-crossing algorithm. The delay between trigger and odour delivery was the same for all plumes at 65ms.

#### Imaging data analysis

For MTC imaging, motion correction, segmentation and trace extraction were performed using the Suite2p package (https://github.com/MouseLand/suite2p and (Pachitariu et al., 2016). Putative neuronal somata were automatically identified by segmentation and curated manually. Fluorescence signal from all pixels within each ROI was averaged and extracted as a time series. ΔF/F = (F − F0)/F0, in which F is raw fluorescence and F0 is the median of the fluorescence signal distribution. Soma and neuropil fluorescence traces were extracted and neuropil fluorescence was subtracted from the corresponding soma trace. All further analysis with custom written scripts in Python 3 and JupyterLab.

*Responsive cells / cell-odour pairs* were defined as cells/cell-odour pairs for which the mean fluorescence during the response window for all trials of an individual odour was above mean + 3*standard deviation of the period 3 s before stimulus onset.

*Distance sensitive cells / cell-odour pairs* were defined as cells showing a significantly different (one way ANOVA) response to one distance category of the four presented. Multiple comparisons arising from these tests were controlled using the Benjamini-Hochberg false-discovery-rate (FDR) procedure with α = 0.05.

Unless otherwise specified, linear classifiers used to understand the ability of the olfactory bulb to represent distance-dependent plumes were all LinearSVC, as implemented in the scikit-learn package (https://scikit-learn.org/). Data was split into a training set and a test set using a 75:25 split stratified by category (distance). Values were then scaled using Standard Scaler. C parameter was set to 0.001.

### Respiration data analysis

Respiration data was extracted using Spike 2 (CED, Cambridge, UK) and further analysed in Python 3. Inhalation was performed using a threshold crossing approach. Traces were normalised within one experiment to the maximum value over the length of the experiment, then segmented into inhalation (above threshold) and non-inhalation (below threshold) phases. This segmentation was used to create a binary signal that was overlaid on the plume traces to calculate the “inhaled plume” and features that took account of inhalation.

### Odour delivery

#### Odours used

All odours were obtained in their pure form from Sigma-Aldrich, St. Louis MO, USA. Odours used: isoamyl acetate (IAA), 1-heptanal (1H), nonanoic acid (NA), α-terpinene (AT), ethyl butyrate (EB), 2-heptanone (2H), benzyl acetate (BA). Odours used in the wind tunnel were presented in their pure form (EB, IAA). For imaging experiments, odours were diluted 1:50 in mineral oil and a further 1:2 in air, resulting in ∼1:100 or 1% concentration reaching the animal.

#### Wind tunnel

To produce olfactory stimuli with similar temporal structures as those measured outdoors (Ackels et al., 2021), a wind tunnel with near-laminar flow was developed. An obstacle placed in the airflow introduced controlled complexity to the airflow. The wind tunnel was square in cross-section and measured 165 cm (L) x 65 cm (W) x 62 cm (H). The frame was constructed from aluminium profiles (MayTec Aluminium Systemtechnik GmbH, Dachau, Germany) and walled with clear acrylic panels. The floor of the tunnel was a laboratory bench placed 5cm lower than the odour port that mice used for sampling. The obstacle was a plastic cylinder (height: 12.7 cm, diameter: 7.6 cm), positioned on the midline 15 cm downstream of the fan, inspired by the routine use of cylindrical obstacles to generate airflow turbulence in fluid dynamics research (Béra et al., 2000).

Wind generation was achieved by a centrifugal blower (D2E133-AM47-23, ebm-papst) placed in the inlet section of the tunnel, with the rear end being open for air to exit the tunnel. The room ventilation system had its outlet placed at the back of the room, above and slightly behind the mouse/PID end of the wind tunnel. To better control the airflow, we used a honeycomb structure (Hexagonal Aluminium W/CRIII Coating 3.0 3/8 .002N 5052, 10 cm in thickness, Texas Almet) placed between the fan and the tunnel, to create a largely laminar airflow. This ensured that the airflow complexity we observed was primarily created by the obstacle. Neutrally buoyant helium-filled soap bubbles (Sage Action, Inc.) were used to visualise airflow, as filmed with a high-speed camera (Basler).

#### Distance Odour Delivery Device (dODD)

The distance Odour Delivery Device (dODD) was used to deliver odours into the airflow in a passive fashion. The device of choice was a custom-built ensemble of an Arduino controlled servo motor (TowerPro SG-5010, Adafruit) that rotates an aluminium lid covering a glass petri dish (5cm diameter). Eight such devices were placed inside the wind tunnel, and controlled via a compact data acquisition (cDAQ) device (National Instruments) and either PulseBoy or the AutonoMouse software (Ackels et al., 2021; Erskine et al., 2019, https://github.com/RoboDoig/PulseBoy and https://github.com/autonomouse-control). This also allowed for integration with the temporal ODD (Ackels et al., 2021), which was adapted and integrated into the system for the imaging experiments. Before each recording session the petri dishes were filled with 10 ml ethyl butyrate (EB), which was the odour used for all plume experiments.

#### Temporal Odour Delivery Device (tODD)

Reproduction of recorded odour plumes was achieved with a temporal Odour Delivery Device (tODD) and custom control software, as previously described in Ackels et al., 2021. In brief, the odour delivery device was based on a modular design of four separate odour channels, and consisted of an odour manifold for odour storage, a valve manifold for control of odour release and hardware for controlling and directing airflow through the system. We used high-speed micro-dispense valves with custom electronics for pulse-width modulation to maximise bandwidth. Two manifolds of four valves were joined together using a three-way connector (TMMA3203950Z, The Lee Company) to create an eight-channel device. Pulse profiles for calibration and stimulus production were generated with custom Python software, allowing us to define pulse parameters across multiple valves using a graphical user interface. The drivers themselves were passed valve opening times via a 5 V TTL pulse from digital I/O controls via a compact data acquisition (cDAQ) device (National Instruments).

#### Odour plume recordings

The temporal structure of odour plumes was recorded using a mini-PID photoionization detector with a bandwidth of ∼330 Hz (200B miniPID, Aurora Scientific, Aurora ON, Canada) and Spike2 software (Cambridge Electronic Design), sampled at 1 kHz. The tip of the detector was placed in the behaviour chamber at the centre of the odour port, to mimic the position of the nose of a mouse performing the task. For each set of recordings, the odour source was alternated between 4 different positions (near or far, left or right) inside the wind tunnel. The PID was calibrated periodically by adjusting the gain and baseline such that it would produce the same response to a series of square pulses of known amplitude (1000 ppm, 100 ppm, 10 ppm isobutylene). Thus, we could compare broad ranges of concentration over different recording sessions

#### Reproduction of recorded plumes

Reproduction of recorded odour plumes was achieved using pulse-width modulation (PWM) generated with custom Python software (PyPulse, PulseBoy, daqface; https://github.com/RoboDoig and https://github.com/warnerwarner) as previously described in Ackels et al., 2021. Briefly, this was achieved by mapping odour signal amplitudes from recorded plume data to PWM duties, converting a time series of amplitudes to a time series of PWM duties. These duties could then be used as the input to an odour valve to reproduce the original signal. First, each PID trace was corrected to baseline drift by performing baseline subtraction, as compared to a 1 s period before each trial. The trace was then normalised to between 0 and 1, as set by the *target_max* variable in PulseBoy, and then converted into a series of binary opening and closing times, at a frequency of 500 Hz. To reproduce the plume using the whole dynamic range of the olfactometer, *target_max* would be set to 1. The length of the openings and closings relate directly to the value of the normalised signal, a value of 1 translates to a continuous opening, and a value of 0 translates to continuously closed. In an offline assignment, *target_max* values were adjusted to group ratios and within group ratios to match mean concentration over the whole plume. Thus, there was variability in the overall concentration presented, but that variability was largely within groups of plumes from the same distance, not between different distances. Generally, wherever an odour position is inactivated a blank position was activated to compensate for flow change. Thus, an inverse sequence of commands as used for the plume generation, termed “anti-plume”, was simultaneously fed to a valve connected to a mineral oil (blank) channel to produce the compensatory flow. This reproduction strategy allowed to precisely control stimulus onset and to fully equalise mean concentration, though, crucially, preserving variability within groups (**Fig. 2H, Fig S2.11, S2.12**).

#### Plume analysis

Plumes were recorded using Spike2 (Cambridge Electronic Design), sampled at 1 kHz. Trial onset was calculated using a TTL pulse passed through an additional channel on the acquisition board (Cambridge Electronic Design). Further analysis was then performed initially in MATLAB and Python 3 using Jupyter Lab. Briefly, traces were segmented using the trial onsets and baseline subtraction performed using the 1 s prior to trial onset. Plume features were defined and calculated as follows: *Correlation* was calculated using the corr function in MATLAB or DataFrame.corr(method=‘pearson’) in Python 3 using the pandas package. *Plume arrival time* was calculated as the time of the first point above 3*SD of the baseline (1s interval before trial onset). *Mean odour concentration* was calculated as the average PID signal over a certain time window as specified. *Maximum peak* was calculated as the maximum value of the PID signal over a certain time window as specified. *Number of peaks* was calculated using the find_peaks function (scipy.signal.find_peaks) in Python 3, using the parameters height = 0.05 and prominence = 0.1 for the reproduced plumes, and height = 0.01 and prominence = 0.01 for the recorded plumes. *Maximum prominence* was extracted from the outputs of the find_peaks function above. *Standard deviation, Variance, Kurtosis and Skewness* were calculated using the corresponding functions in the scipy.stats library for different time windows as specified. *Reproduced plume integral* and *Odour inhaled* were calculated using the integrate.trapz function in scipy. *Cumulative odour presented* and *Cumulative odour inhaled* were calculated using the cumsum function in numpy.

### Statistical analysis and data display

To test for statistical significance between individual groups where appropriate we used either paired or unpaired student t-tests or, for non-parametric data, the Mann–Whitney U test, or the Kolmogorov-Smirnov test for the equality of probability distributions. Statistical test details and P values are provided in figures and/or legends.

Data figures were plotted using Jupyter Lab using seaborn, pandas, sci-kit learn or matplotlib functions in Python 3. Unless specified otherwise, boxplots depict the median as a thick line and default maximal whisker length of 1.5 × (q3 − q1) where q3 and q1 indicate the 75th and 25th percentile, respectively. If points were located outside this whisker range, they were displayed individually as outliers. Unless otherwise specified, line plots depicting animal performance show mean (thick line) ± SD (shaded area) for a rolling window of blocks of 100 trials.

## Data availability

Source data supporting the graphs presented in the main figures are provided as a repository from https://zenodo.org/records/15390336.

## Code availability

Analysis code to recreate the main figures is available from https://github.com/ackels-lab/Odour_distance_paper. All custom analysis code supporting the findings of this study will be made publicly available through the same code repository upon publication.

## Acknowledgements

This work was supported by the Francis Crick Institute which receives its core funding from Cancer Research UK (FC001153), the UK Medical Research Council (FC001153), and the Wellcome Trust (FC001153); a Wellcome Trust Investigator (110174/Z/15/Z) grant and the NeuroNex program “From Odor to Action” to A.T.S., a BIF doctoral fellowship to A.C.M., and a DFG postdoctoral fellowship to T.A. It was further supported by the German Research Foundation (FOR5424 “Modolfor”, A.T.S and T.A) and the European Union (ERC, “TempCOdE”, 101077017, T.A.). Views and opinions expressed are, however, those of the author(s) only and do not necessarily reflect those of the European Union or the European Research Council. Neither the European Union nor the granting authority can be held responsible for them.

