## Supplementary material for "Mice discriminate odour source distance via sub-sniff temporal features of odour plumes": Main figures (high resolution)

### Figure 1

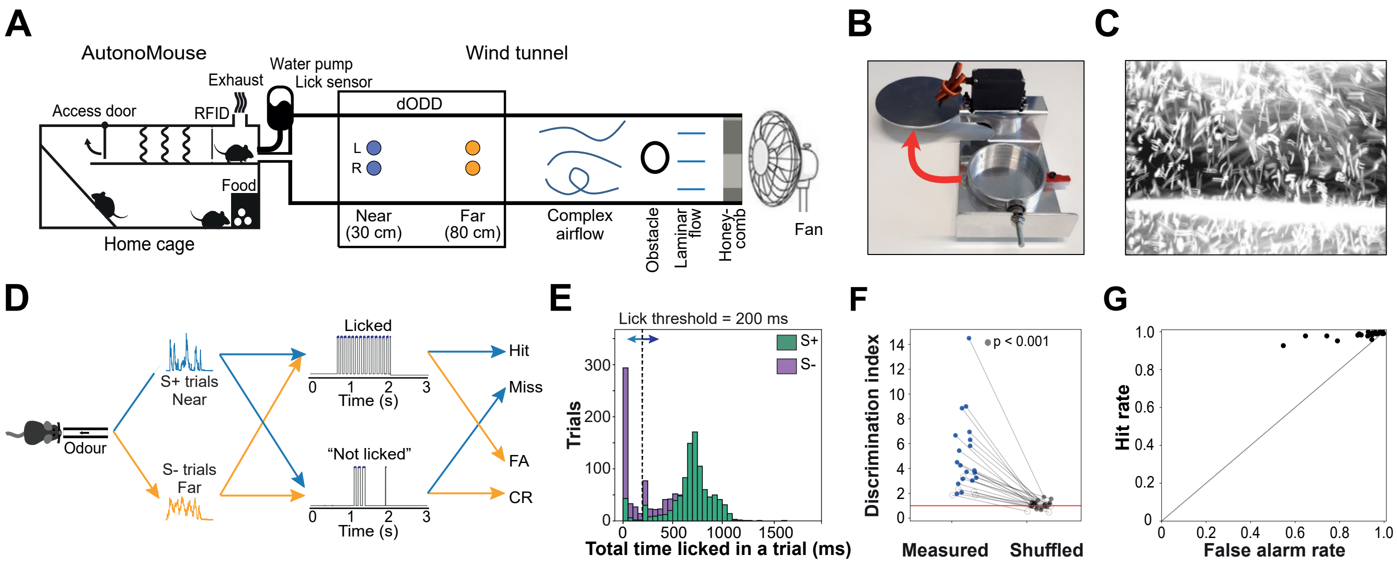


**Figure 1: Mice can discriminate between near and far odour sources. (A)** Complex airflow is generated inside a wind tunnel using a fan, a honeycomb structure and an obstacle. Odour sources are placed downstream of the fan and upstream of the detector (PID), forming the distance odour delivery device (dODD). The dODD is formed of 8 odour boxes, 4 at each distance (near or far) from the PID, arranged symmetrically across the midline of the tunnel (Left and Right positions). Mice (n = 24) were housed in an automated operant conditioning system (AutonoMouse), where the only water source was the reward from the GNG task. **(B)** Odour presentation boxes consisting of a metal casing surrounding a glass dish. An Arduino controlled servo motor opens the lid to allow airflow to pick up odour molecules. The lid opens horizontally to the table for minimum airflow disturbance. **(C)** Complex airflow in the wind tunnel generated by a honeycomb structure and a cylindrical obstacle placed 25 cm downstream of the fan. **(D)** Schematic of task structure: Depicted is the reward structure for half of the animals (n = 13), where plumes originating from the near source (30 cm) were associated with a reward (S+ trials) and the plumes originating from the far source (80 cm) were not associated with a reward (S- trials). A lick response (top) would allow for early termination of the trial if the total time licked (shaded in blue on the lick trace) was >200 ms in a 2 s rolling window. The absence of a lick response (bottom) was defined as total time licked <200 ms in a 2 s rolling window, and the trial would continue until its maximum length was reached at 3s. These two categories of responses to the two types of trials (S+ or S-) produced the outcomes listed on the right-hand column (Hit, Miss, False Alarm, Correct Rejection). **(E)** Histograms of total time licked for an example mouse, as calculated by the AutonoMouse software to decide upon delivering a water reward, using time above threshold (dotted line) in the lick traces as shown in D (S+ in blue, S- in orange). **(F)** Discrimination index defined as the ratio of total time licked (S-/S+) for each mouse below the threshold (t = 200 ms), compared to the mean of shuffled values for that mouse. The measured value is significantly higher than shuffle (measured>shuffled in 99% of shuffles) in 19/24 mice (filled dots). **(G)** Hit rate (fraction of S+ trials licked) plotted against False alarm rate (fraction of S- trials licked) averaged across a period of 10 blocks of 100 trials each during a stable period of the experiment. Each dot is an individual mouse. A perfect block would be placed in the upper left corner, while a block with continuous licking would be placed in the upper right corner. A block where the mouse was not engaged in the task would be placed in the lower left corner. Indiscriminate licking while engaged in the task would be placed along the diagonal.

### Figure 2

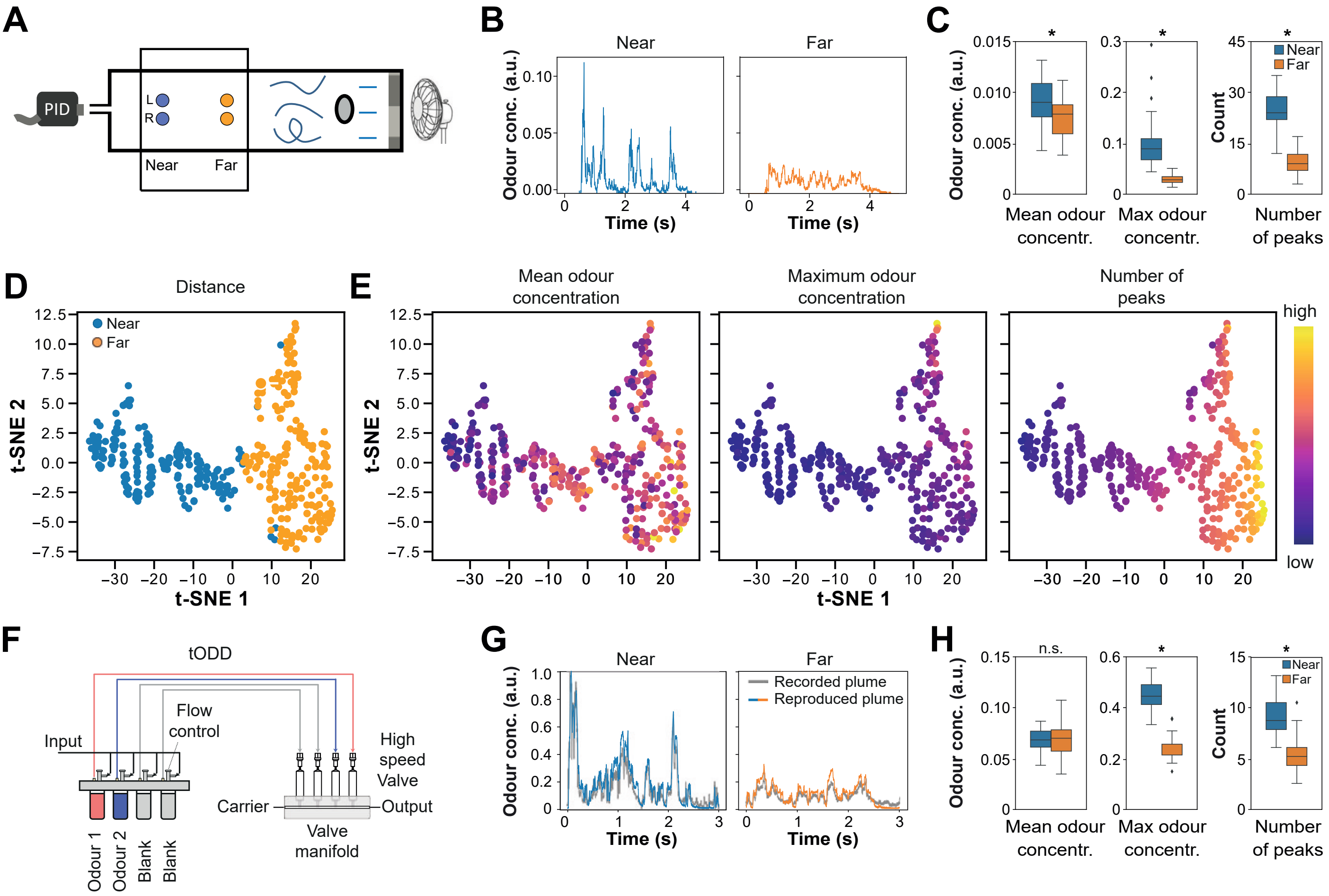


**Figure 2: Odour plume structure varies reliably with distance and sub-sniff features are the most informative. A)** Schematic of the distance Odour Delivery Device (dODD). Complex airflow is generated inside a wind tunnel using a fan, a honeycomb structure and an obstacle. Odour sources are placed downstream of the fan and upstream of the detector (PID) and arranged symmetrically across the midline of the tunnel (Left and Right positions). **(B)** Example plumes recorded from the Near (30 cm) and Far (80 cm) positions. **(C)** Quantification of Mean odour concentration (left), Maximum odour concentration (middle) and Number of peaks (right) for near and far plumes. Box indicates 25th-75th percentiles, thick line is median, whiskers are most extreme data points not considered outliers. **(D)** tSNE analysis of odour distance calculated for a 5 s time window for each plume. Each dot is a trial (n = 360; 180 near and 180 far plumes). **(E)** Same as (D) but for plume temporal structure features. Dots are colour-coded the value of individual temporal structure features to visualise any gradients preserved in the two-dimensional space. **(F)** Schematic of temporal Olfactory Delivery Device (tODD). **(G)** Example plumes from near and far distances. Recorded plumes in grey, reproduced plumes in blue (near) or orange (far). **(H)** Same as in (C) but for plumes reproduced using tODD. Box indicates 25th-75th percentiles, thick line is median, whiskers are most extreme data points not considered outliers.

### Figure 3

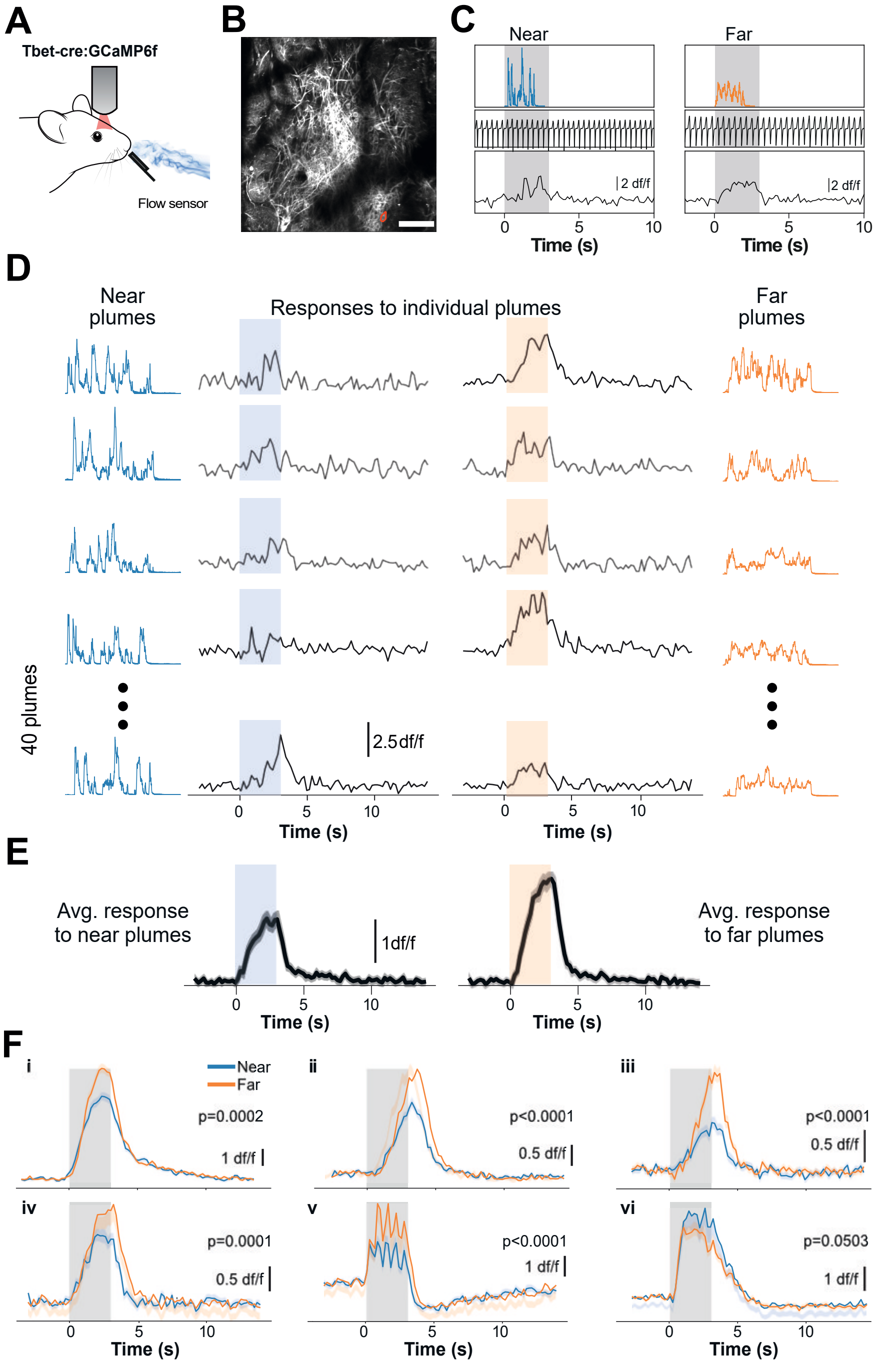


**Figure 3: Responses of MTCs to temporally structured odour plumes recorded from different distances. (A)** Schematic of the two-photon imaging approach. **(B)** GCaMP6f fluorescence from mitral and tufted cells (maximum projection of 8,000 frames). Responses from ROI marked in red are shown in (C). Scale bar, 100 µm **(C)** Example single trials in response to an odour stimulus from a near (30 cm) or far (80 cm) virtual odour source (top row). Respiration traces (inhalation pointing upwards) from the same trials (middle row). Example fluorescence traces (df/f) from the red circled ROI in (B). Shaded area represents 3 s odour stimulation window. **(D)** Individual near and far plumes (outer columns) and corresponding calcium responses (inner columns) of one ROI. **(E)** Averaged responses to all near and far plumes (n = 40 for each condition, mean ± s.e.m.). **(F)** Average fluorescence traces (mean ± s.e.m., n = 160 trials) for 6 example distance sensitive ROIs (near: blue, far: orange; ROIs i-iv: EB, v-vi: 2H).

### Figure 4

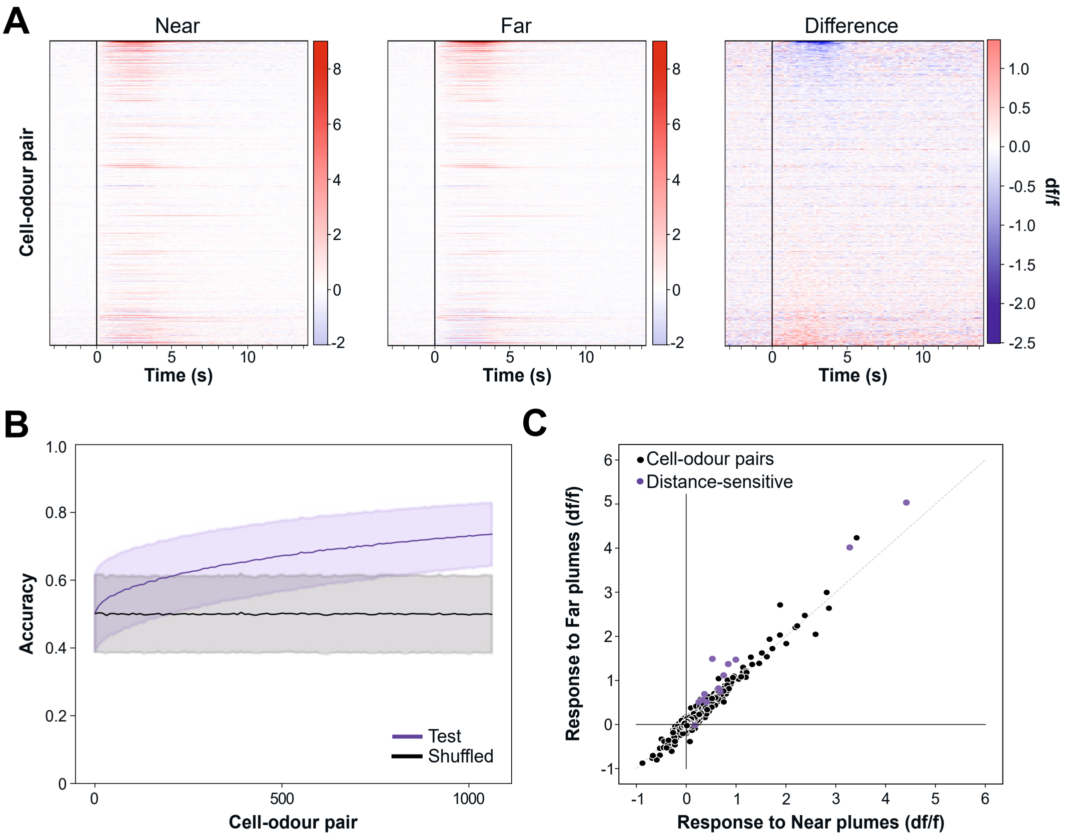


**Figure 4: Population activity and linear discriminant analysis of MTC responses. (A)** Calcium transients as colour maps for averaged responses to near (left) and far (middle) plumes and for the difference between the two (right) for all cell-odour pairs (n = 1062). **(B)** Accuracy of linear classifier trained on a 5 s response window (purple; black, shuffle control) to near versus far plumes (n = up to 1062 cell-odour pairs from 3 mice; mean ± s.d. of 50000 repetitions). **(C)** Responses to near vs responses to far plumes plotted as mean df/f response for all trials from those distances. Distance-sensitive cells shown in purple. Cells above diagonal respond more strongly to far plumes while cells below diagonal respond more strongly to near plumes.

### Figure 5

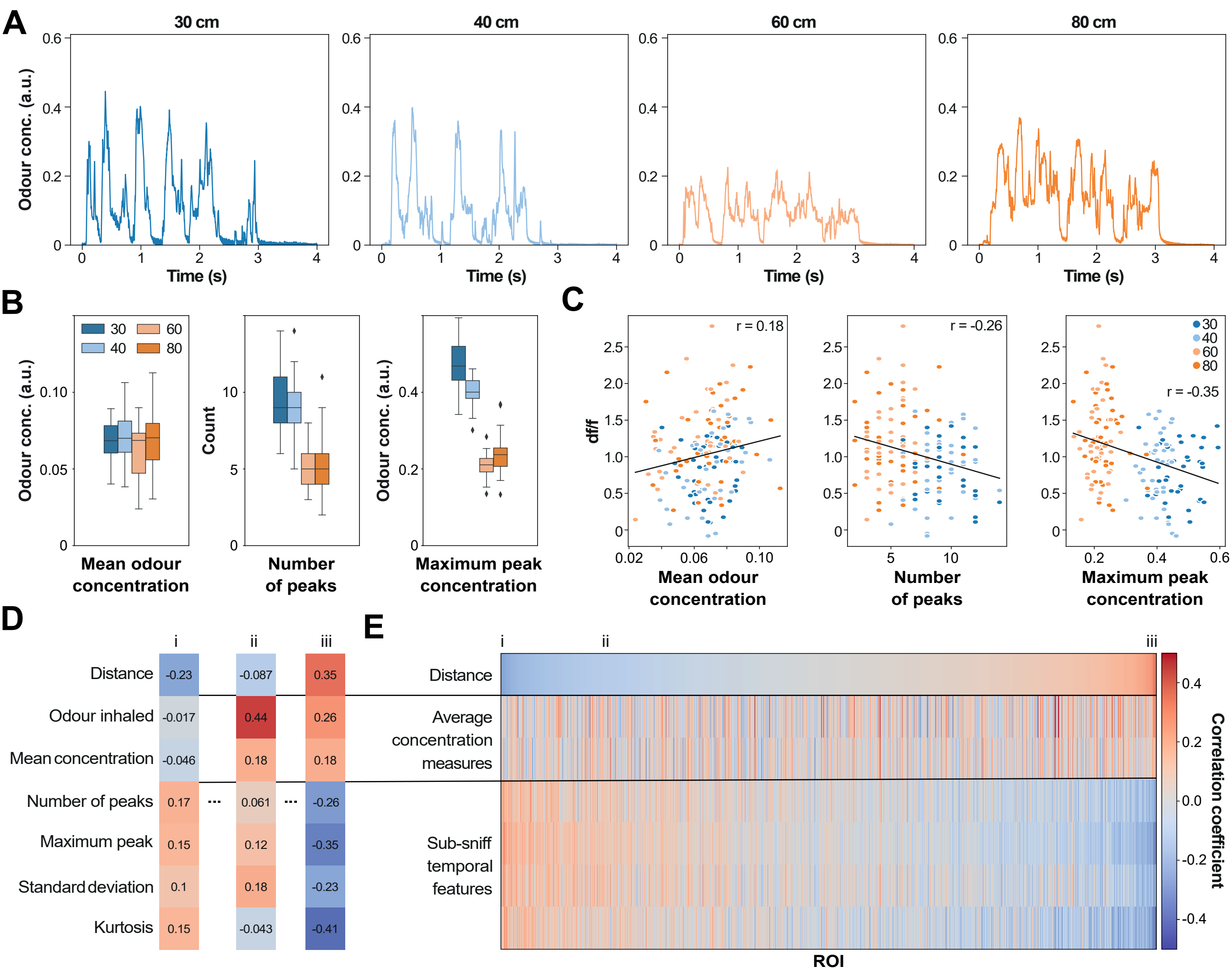


**Figure 5: Features of the temporal structure of odour plumes involved in distance discrimination. (A)** Example plumes recorded from four different distances (30 cm, 40 cm, 60 cm and 80 cm) positions. **(B)** Quantification of mean odour concentration (left), number of peaks (middle) and maximum odour concentration (right) for all plumes from four distances. Box indicates 25th-75th percentiles, thick line is median, whiskers are most extreme data points not considered outliers. **(C)** Response of an example distance-sensitive ROI (df/f averaged over the 5 s response window) plotted against individual features of the odour stimulus (x-axis). Each dot is a trial (n = 160 trials), colour-coded by distance (80 cm orange, 60 cm pale orange, 40 cm pale blue, 30 cm blue). **(D)** Correlation coefficients to distance and other stimulus features for three ROIs (i-iii). **(E)** Heatmap of correlation coefficients as calculated in C for all cells (columns, n = 531 ROIs from 3 animals), sorted by their correlation to distance (top row). Labels correspond to ROIs shown in (D).
