## Supplementary Figures (high resolution) for "Mice discriminate odour source distance via sub-sniff temporal features of odour plumes"

### Figure S1.1

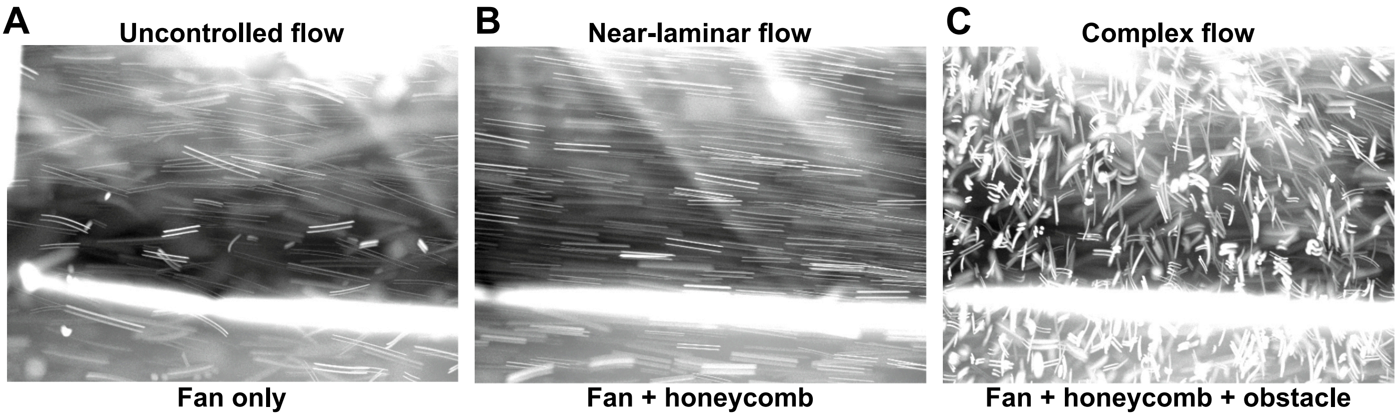


**Figure S1.1: Complex airflow in the wind tunnel generated by a known obstacle.** Maximum intensity projection through a 10 s video of neutrally buoyant soap bubbles in the wind tunnel, filmed 30 cm downstream from the fan, immediately behind the obstacle. Bubbles appear elongated due to the exposure time of the camera. Straight streaks of equal lengths are an indication of laminar airflow. **(A)** Turbulence in the air coming directly from the fan indicates unknown and uncontrolled sources of turbulence. **(B)** When a honeycomb structure is added to straighten the airflow, a near laminar pattern is observed. **(C)** A cylindrical obstacle placed 25 cm downstream of the fan introduces reproducible turbulence to the airflow.

### Figure S1.2

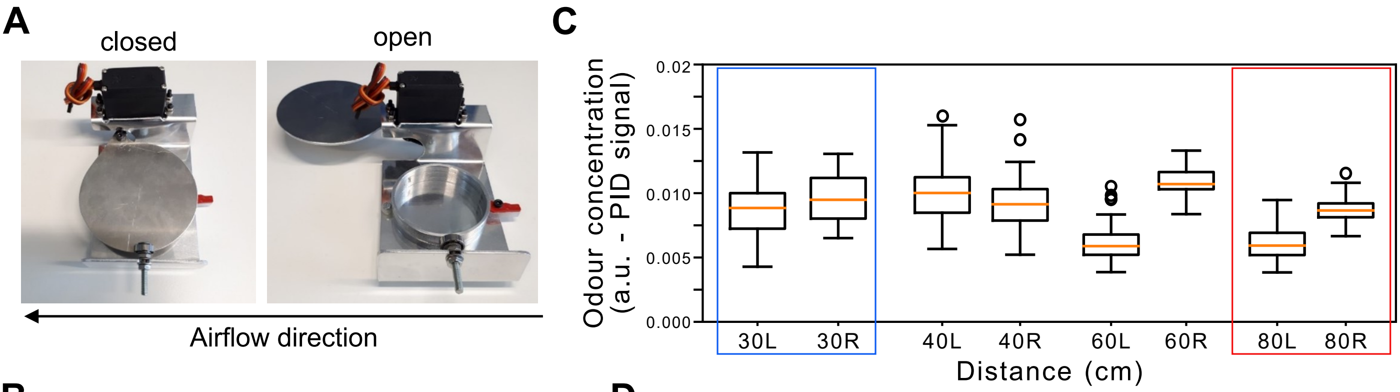


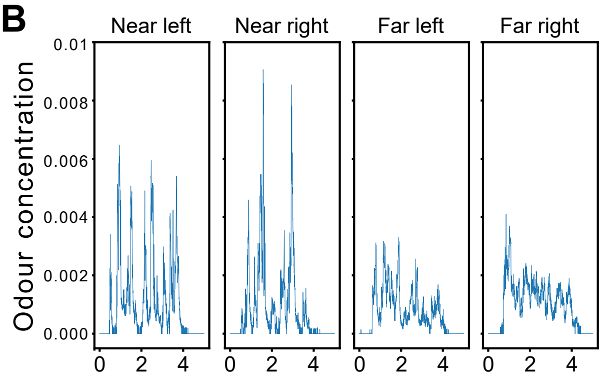


D

| **Concentration ($x10^3$)** | **30** | **40** | **60** | **80** |
|---|---|---|---|---|
| Left | 8.8±2.0 | 10.1±2.2 | 6.2±1.6 | 6.1±1.4 |
| Right | 9.6±1.9 | 9.4±2.0 | 10.9±1.1 | 8.7±0.8 |
| L vs R | P=0.083 | P=0.099 | P<0.005 | P<0.005 |
| Both | 9.2±2.0 | 9.7±2.1 | 8.5±2.7 | 7.4±1.7 |

**Figure S1.2: Plume generation in the wind tunnel and statistics. (A)** Individual odour boxes consist of a metal casing surrounding a glass dish where liquid odour can be added. An Arduino controlled servo motor can then open the lid to allow airflow to pick up odour molecules. The lid opens horizontally to the table for minimum airflow disturbance. **(B)** Example plumes recorded from the positions marked in A: Near (30 cm) Left and Right, Far (80 cm) Left and Right, as viewed from the detector/odour sampling port. **(C)** Mean odour concentration over a 5 s period for plumes recorded from 4 distances, both left and right. Plumes from B belong to the categories highlighted in blue (near) and red (far). Box indicates 25th–75th percentiles, thick line is median, whiskers are most extreme data points not considered outliers. **(D)** Quantification of C as mean ± s.d. for each position and comparison between Left and Right for each distance (unpaired t test). Bottom row means ± s.d. for each distance.

### Figure S1.3

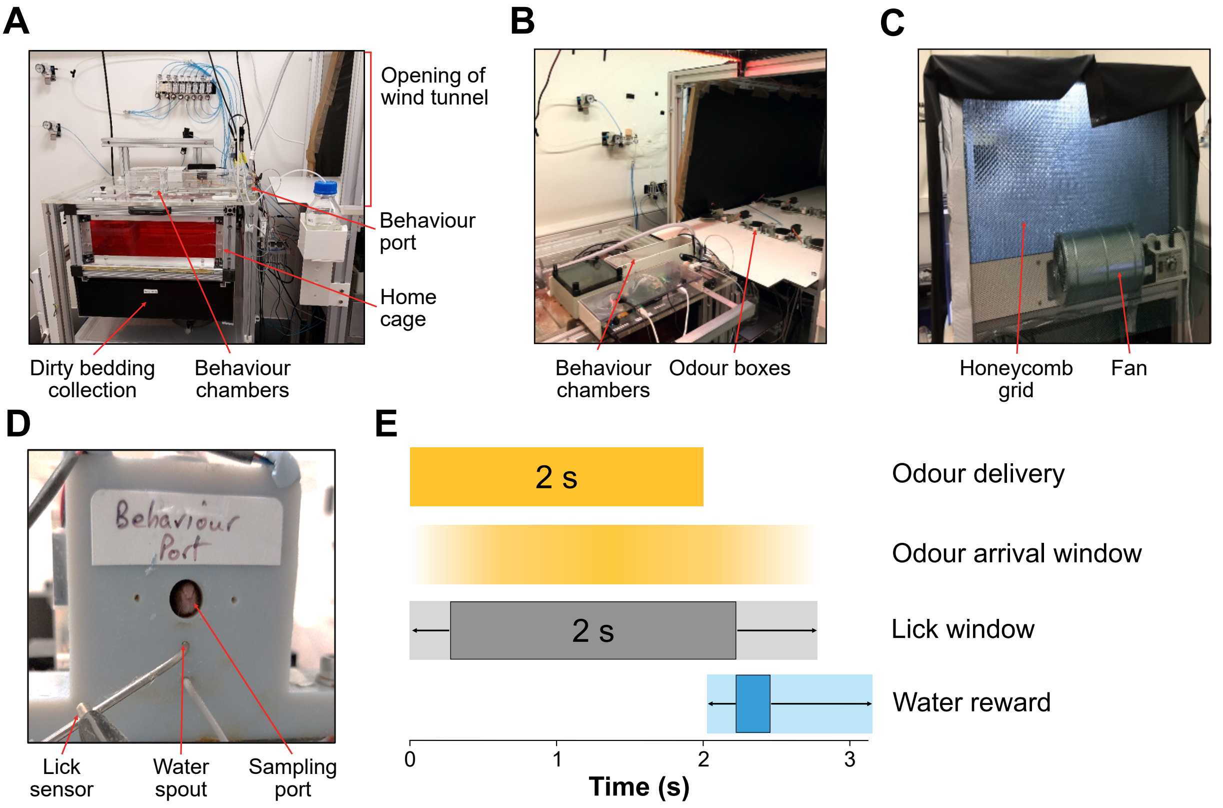


**Figure S1.3: AutonoMouse system for training cohorts of mice. (A)** Picture of the AutonoMouse training system. **(B)** Picture of the wind tunnel opening and odour boxes. **(C)** Rear view of the wind tunnel showing the fan and honeycomb grid. **(D)** Picture of the modified odour port that allows mice to sample odours originating from the wind tunnel. The opening is 1cm in diameter and is placed above the lick spout, which is connected to a lick sensor and also delivers the water rewards. **(E)** Timeline of the Go/No-Go task paradigm with adaptive/variable lick window.

### Figure S1.4

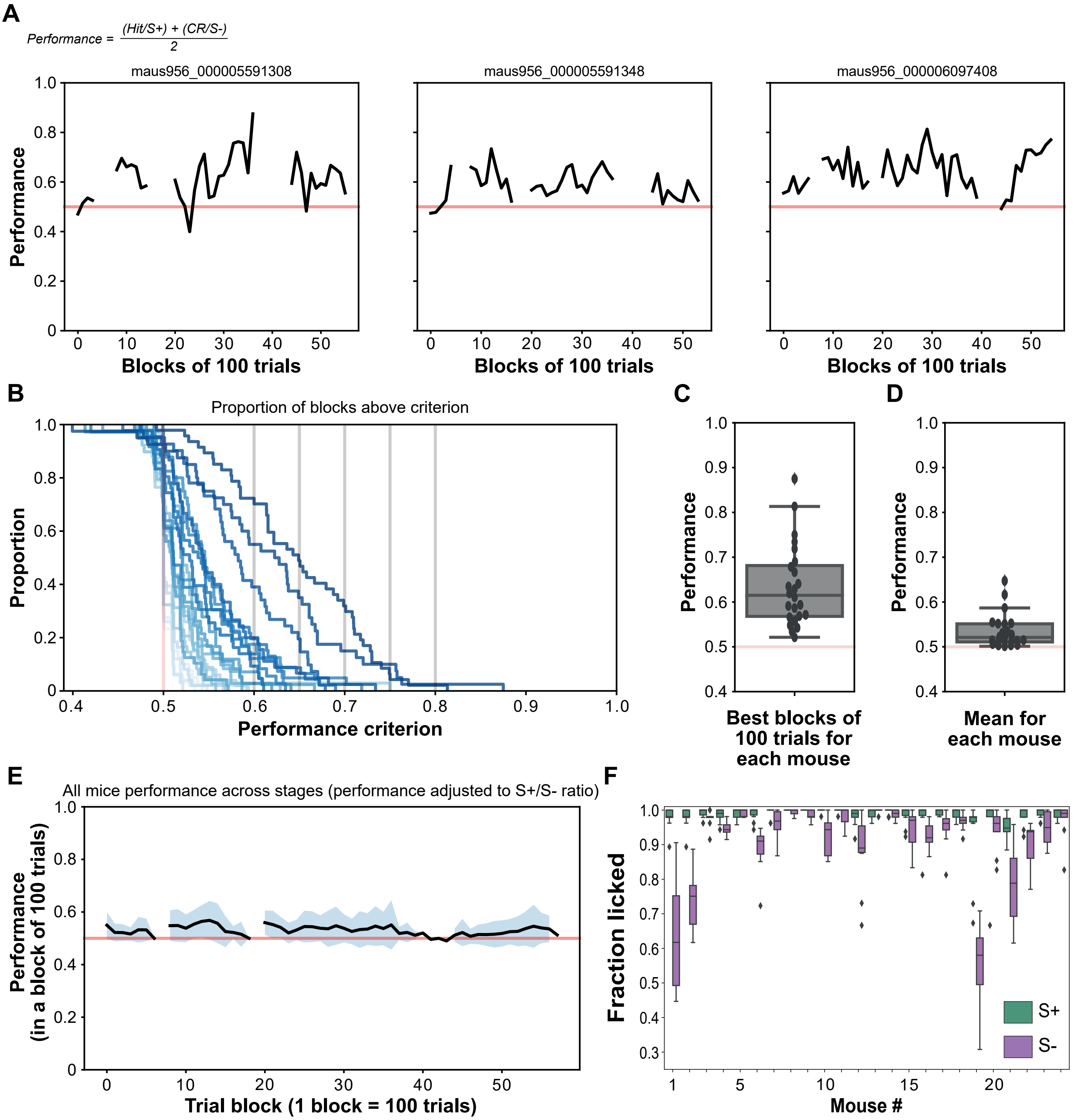


**Figure S1.4: Learning performance of the distance-discrimination task. (A)** Example learning curve for three mice trained to perform the distance discrimination task. Two mice were trained to lick in response to the Far source and refrain from licking in response to the Near source (Group 1, left and middle), while the third mouse was trained on the reverse valence (Group 2, right). Performance was calculated for blocks of 100 trials, taking into account the split of S+ and S- trials in that block. The gaps illustrate periods when the experiment was interrupted for maintenance work and the length of the gaps is such that block number could be realigned between different mice when the experiment restarted, despite different mice performing different numbers of trials during the experiment. Red line indicates chance performance at 50%. **(B)** Proportion of blocks (100 trials/block) with a performance above criterion (x axis). Each line represents a mouse, colour-coded by mean performance. Darker shade indicates higher performance. The intersection between each curve and the x axis represents the highest performance reached by each mouse in an individual block. **(C)** Best performance over a block (100 trials) reached by each mouse. Box indicates 25th-75th percentiles, thick line is median, whiskers are most extreme data points not considered outliers, dots are individual mice. **(D)** Mean performance for each mouse calculated as the mean of all blocks performed by that mouse. **(E)** Average learning curve for all mice trained to perform the distance discrimination task (mean±s.d.). Performance was calculated over blocks of 100 trials, taking into account the split of S+ and S- trials in that block. Gaps illustrate periods when the experiment was interrupted for maintenance work. Data for individual mice was realigned to the start of each stage. **(F)** Fraction of trials in which the mice responded by licking plotted for individual mice split by S+ (green) or S- (magenta). A perfectly performing mouse would score 1 for S+ trials and 0 for S- trials. A mouse licking all the time would score 1 for both, while a mouse not engaged in the task would score 0 for both. A mouse licking indiscriminately would score the same for both S+ and S- trials. Box indicates 25th–75th percentiles, thick line is median, whiskers are most extreme data points not considered outliers. Analysis performed on a period of 10 blocks of 100 trials each during a stable period of the experiment.

### Figure S1.5

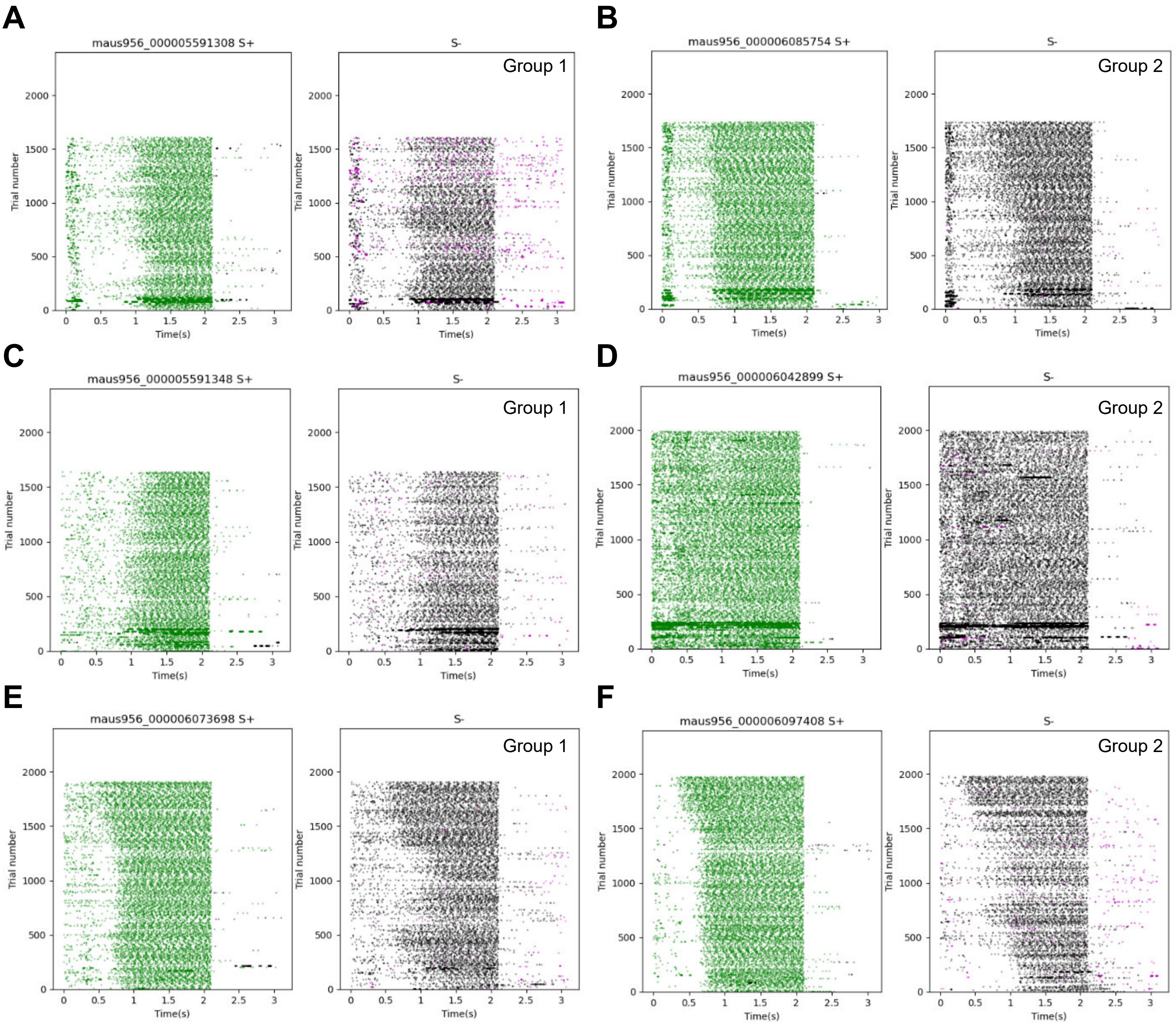


**Figure S1.5: Example lick responses.** Lick responses plotted for 6 example mice **(A-F)** for S+ trials (left) and S- trials (right) during one experimental period. Each dot is a lick event, calculated through a threshold crossing method from lick traces as shown in Figure 1, each row is a trial. Green dots indicate Hits, magenta dots indicate Correct Rejections, while black dots indicate Misses (left) or False Alarm (right). Example mice picked from both Groups 1 and 2 as indicated.

### Figure S1.6

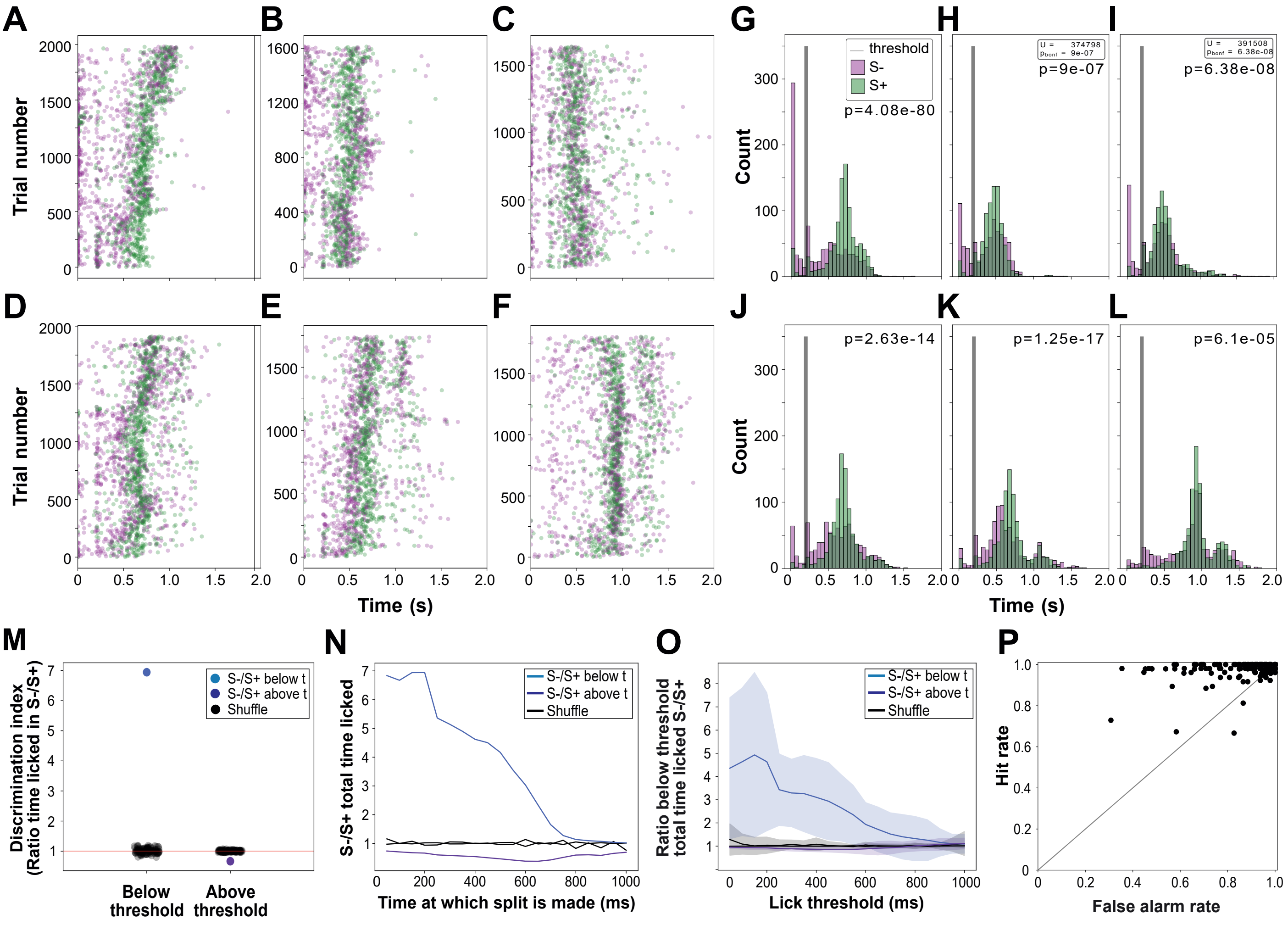


**Figure S1.6: Example lick durations. (A-F)** Total time licked for the same mice as in Figure S1.5. Durations was calculated by the AutonoMouse software to decide upon delivering a water reward, using time above threshold in the lick traces as shown in Figure 1. Each line is a trial. Each dot is the total time licked for that trial colour-coded by trial type: S+ in green, S- in magenta. **(G-L)** Histograms of total time licked for A-F. Vertical line is the threshold (200 ms) used by the software to determine whether a trial was licked or not (p-values determined by Bonferroni-corrected Mann-Whitney U test). **(M)** Discrimination index defined as the ratio between number of S- trials and number of S+ trials with a lick duration smaller than a 200ms threshold (blue), or larger than the threshold (violet) for one example mouse. Grey dots are similarly calculated for each mouse while shuffling the reward. A value >1 indicates more S- trials; a value <1 indicates a preference for S+ trials. A value close to 1 signifies no preference. A perfectly trained mouse with no bias would tend towards infinity for below threshold and 0 for above threshold. **(N)** Same as M but for multiple time thresholds (x axis). **(O)** Same as N but for all mice grouped together (mean ± s.d., shaded area). **(P)** Hit rate (fraction of S+ trials licked) plotted against False alarm rate (fraction of S- trials licked) during a period of 10 blocks of 100 trials each during a stable period of the experiment. Each dot is a block of 100 trials. A perfect block would be placed in the upper left corner, while a block with continuous licking would be placed in the upper right corner. A block where the mouse was not engaged in the task would be placed in the lower left corner. Indiscriminate licking while engaged in the task would be placed along the diagonal.

### Figure S2.1

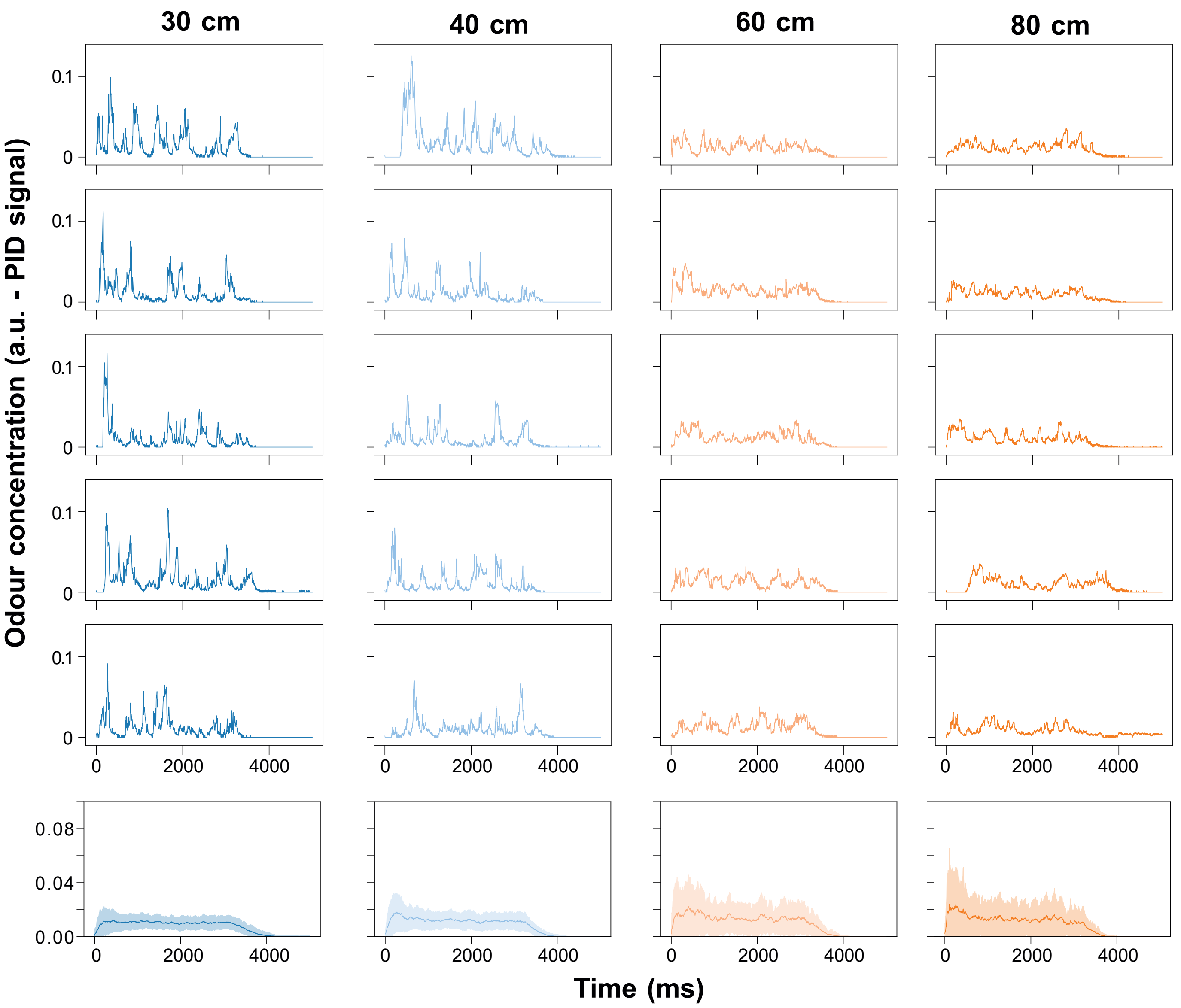


**Figure S2.1: Example plumes from different distances.** Temporal structures of individual odour plumes for each distance (top 5 rows, each column is one distance). Bottom row: average plume structure (mean ± s.d. for all plumes recorded for each distance). Colours indicate source distance (80 cm orange, 60 cm pale orange, 40 cm pale blue, 30 cm blue).

### Figure S2.2

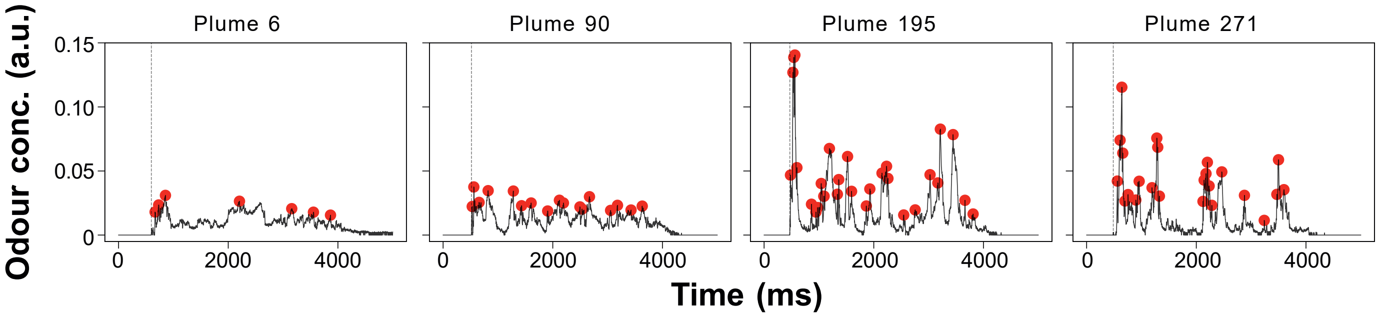


**Figure S2.2: Calculation of plume features.** Example temporal structures of four plumes. Red dots indicate identified peaks. Maximum peak is defined as the height of the highest peak (first red arrow). The peak finding function also provides the prominence for each peak, in addition to the position and the height of the peak. The second red arrow shows the prominence for the first identified peak. Maximum prominence is then defined as the largest prominence value for all peaks in a specified time window.

### Figure S2.3

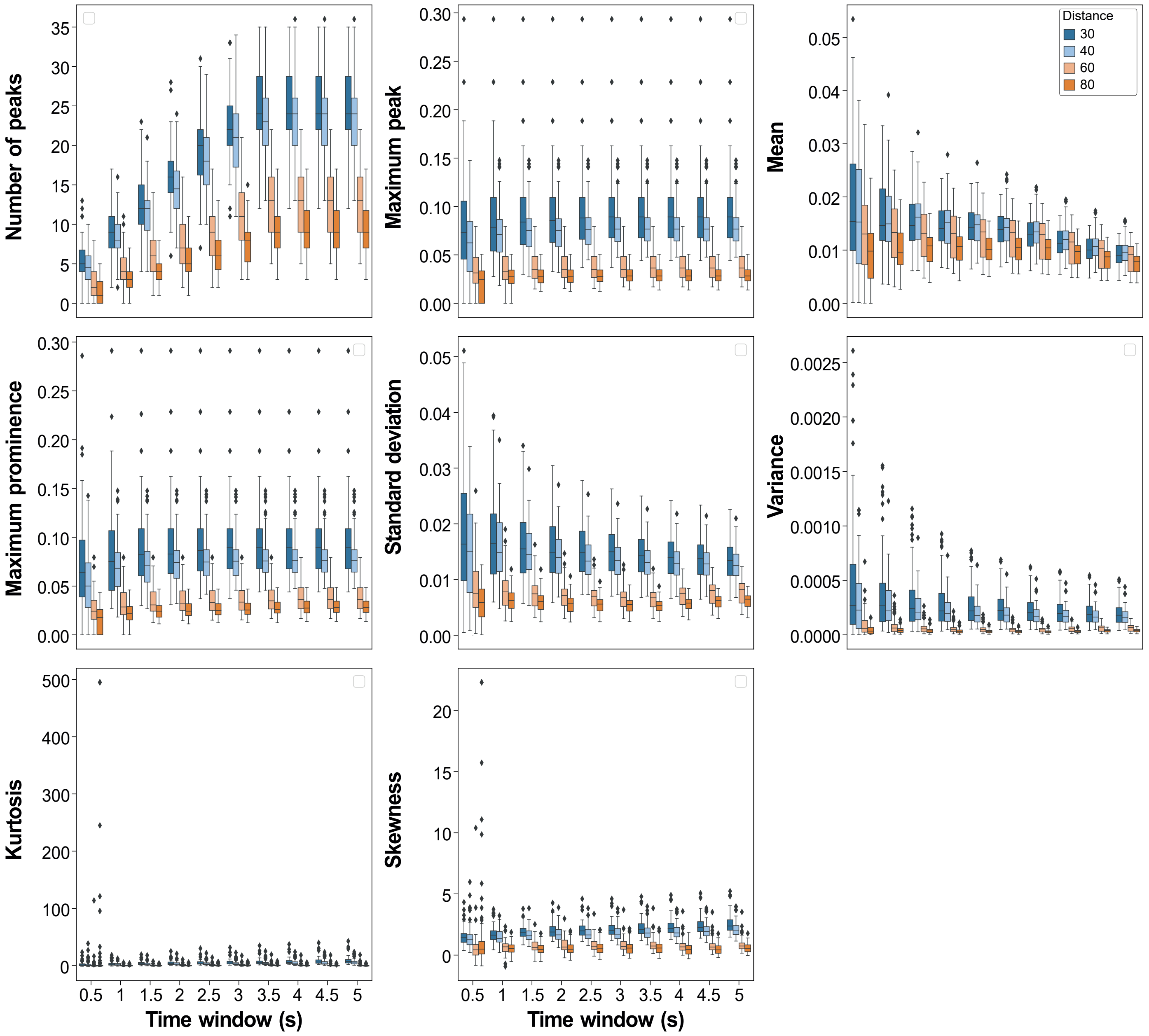


**Figure S2.3: Temporal structure features of plumes from different distances.** Box plots of plume features for intervals of increasing length between time = 0ms, as defined by odour arrival, and different time points as shown on the x axis. Colours indicate source distance (80 cm orange, 60 cm pale orange, 40 cm pale blue, 30 cm blue). Box indicates 25th-75th percentiles, thick line is median, whiskers are most extreme data points not considered outliers. All features are different between 30 cm and 80 cm for all time windows (Kruskal-Wallis test with Dunn correction).

### Figure S2.4

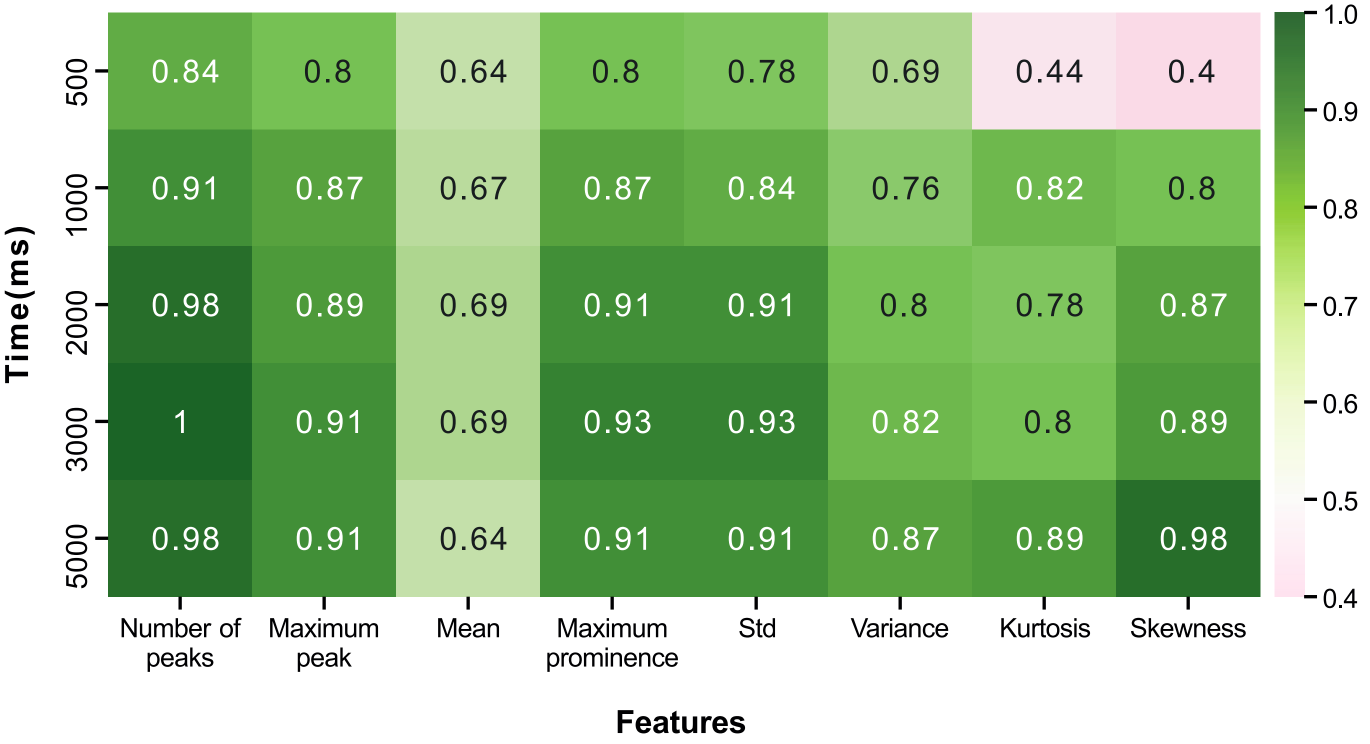


**Figure S2.4: Accuracy of 30 cm vs 80 cm discrimination using different temporal structure features over different time windows.** Classification accuracy on a test set of 45 plumes for a linear discriminant trained on 135 plumes, using only one feature (columns) and one time window (rows) at a time. The split was stratified by distance such that equal numbers of 30 cm and 80 cm plumes were present in the training and the test set.

### Figure S2.5

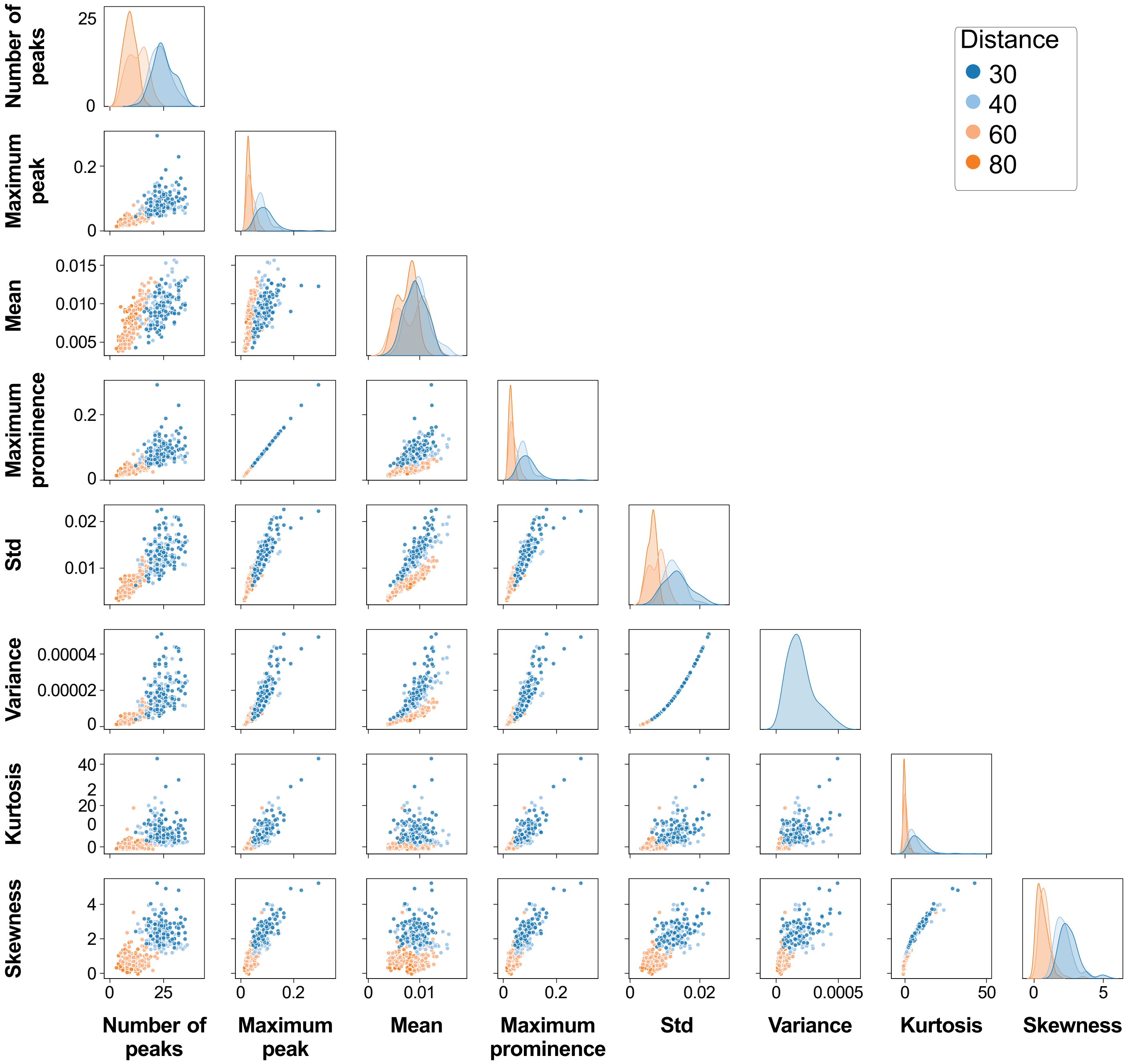


**Figure S2.5: Pairwise comparisons of temporal structure features calculated over a 5 s trial.** Each off-diagonal square is a scatter plot of two features, where every dot is a trial, colour-coded by distance (80 cm orange, 60 cm pale orange, 40 cm pale blue, 30 cm blue). The diagonal squares are distribution plots drawn to show the marginal distribution of data in each column, colour-coded for each distance

### Figure S2.6

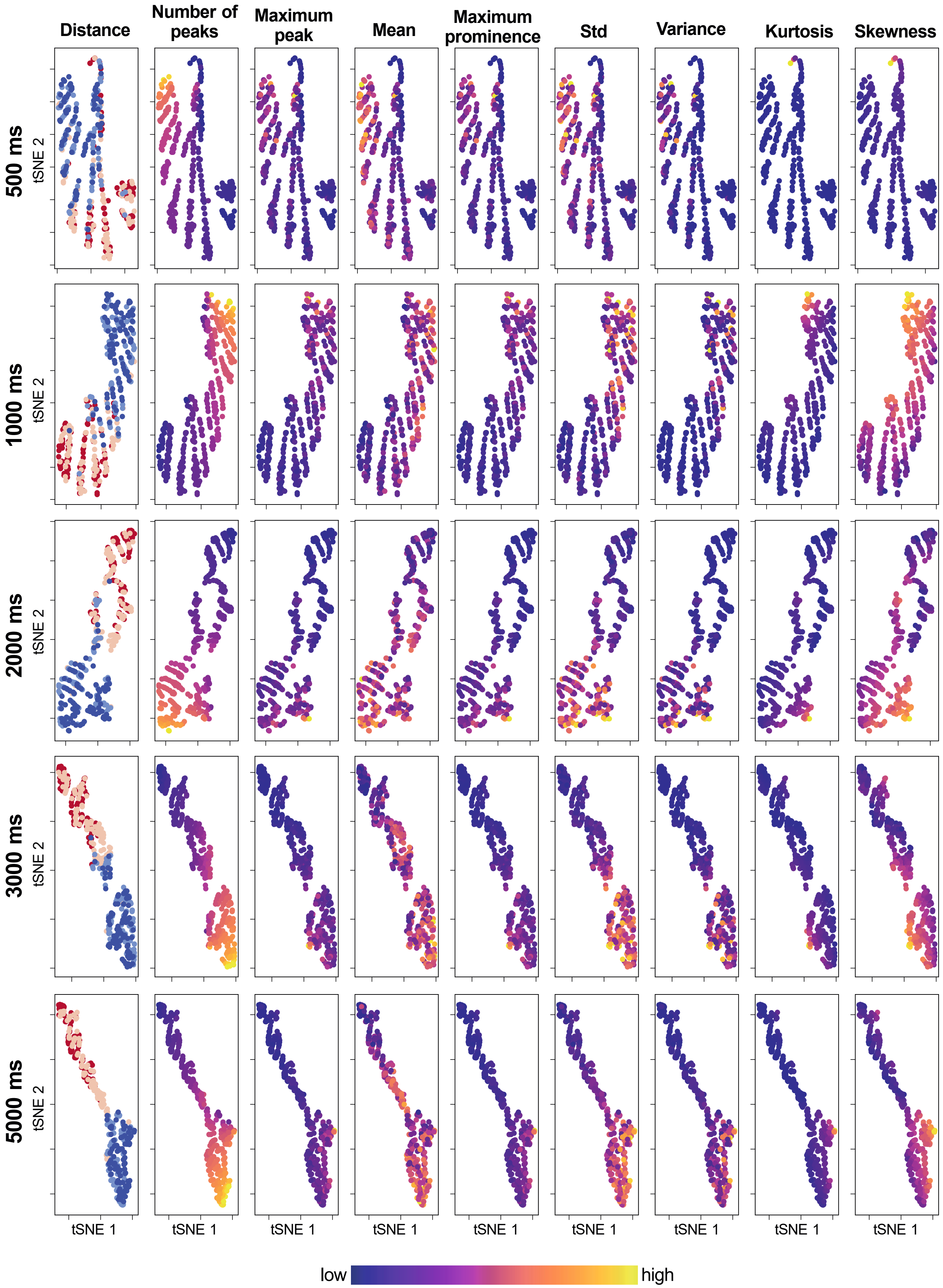


**Figure S2.6: Two-dimensional representation of distance plumes based on temporal structure features, highlighting the near and far discrimination.** t-distributed stochastic neighbour embedding of 8 temporal structure features into a two-dimensional space for better visualisation. Each dot is a trial (n = 360, 90 for each distance, colours as in Fig. S2.5: blue = 30 cm, pale blue 40 cm (both "Near" in Fig 2), pale red 60 cm, red 80 cm (both "Far"). tSNE analysis was performed separately for different time windows (rows) for which the temporal structure features were calculated. The resulting two-dimensional representation is then shown multiple times (columns), where each dot is colour-coded by distance (1st column) or the value of individual temporal structure features (columns 2-9), to visualise any gradients preserved in the two-dimensional space. This would indicate a greater contribution of that feature to building this representation.

### Figure S2.7

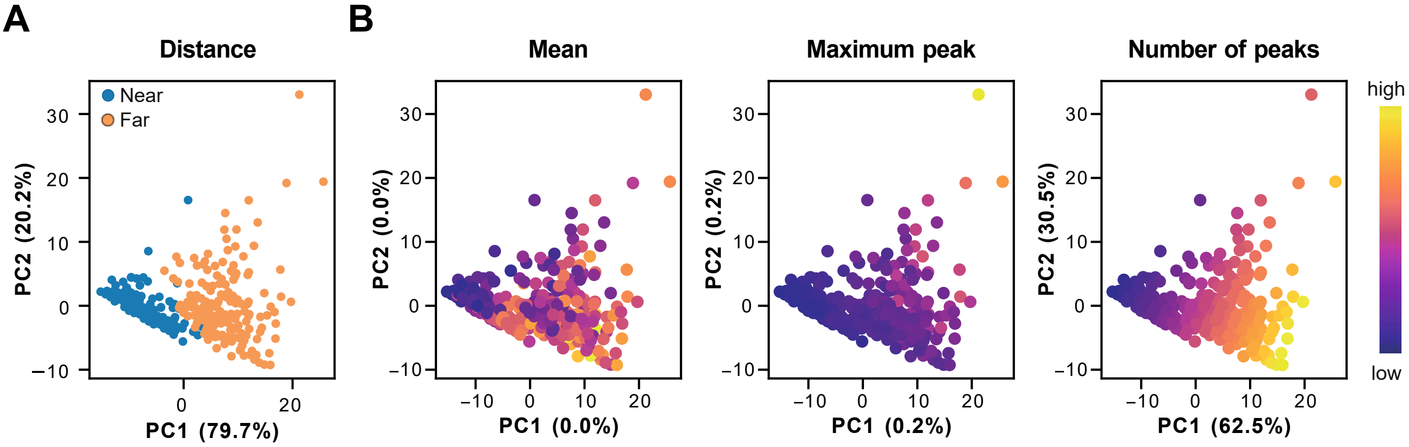


**Figure S2.7: Two-dimensional representation of distance plumes based on temporal structure features. (A)** Principal component analysis of odour distance calculated for a 5 s time window for each plume. Each dot is a trial (n = 360; 180 near and 180 far plumes). **(B)** Same as (A) but for plume structure features. Dots are colour-coded by the value of individual temporal structure features to visualise any gradients preserved in the two-dimensional space. Explained variance for individual principal components shown on the axis labels.

### Figure S2.8

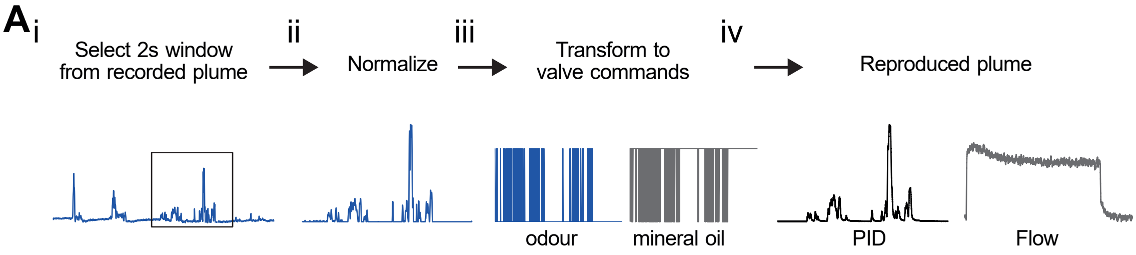


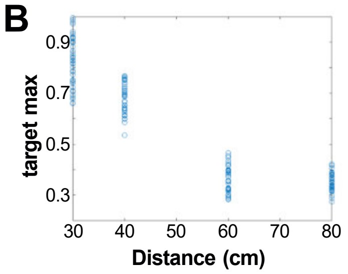


C

| Distance (cm) | 30 | 40 | 60 | 80 |
|---|---|---|---|---|
| **Group ratios** | 0.43 | 0.47 | 0.77 | 1.00 |

**Figure S2.8: Schematic of plume reproduction and normalisation strategy. (A) (i)** In order to deliver temporally structured odours, a 2 s window was first selected from the PID recording. When the recording was longer than the intended reproduced stimulus, the window was selected from the middle of the trace and such that odour was present during the first 500 ms. **(ii)** Second, the trace was normalized between 0 and 1. **(iii)** Third, the trace was converted into a series of binary opening and closing commands directly related to the value of the normalized signal. A value of 1 translated to a continuous opening, and a value of 0 translated to continuously closed. This series of commands was relayed to an odour valve and an inverted version of the commands was relayed to a mineral oil valve to generate a compensatory airflow. **(iv)** The resulting output resembled the original plume, as measured with a PID, and there was constant airflow throughout the trial, as measured with a flow meter. If multiple odours were used, the same procedure was then applied to the accompanying odour, to create both plumes. **(B)** Final *target_max* values used to reproduce the plumes from different distances, which effectively set the maximum height for each trial. Each blue circle is the *target_ max* for each plume (n = 40 plumes / distance). Each value was calculated by multiplying a group ratio and a within group ratio. The values for within groups were obtained for each plume by normalising each plume within a group (distance) to the highest peak within that group (distance), thus preserving relationships within groups. **(C)** The group ratios were calculated by comparing the mean plume integral between groups if *target*_max had been set to 1. The final values used for target_max were all above 0.25, within the range that odour release is linearly related to the PWM duty.

### Figure S2.9

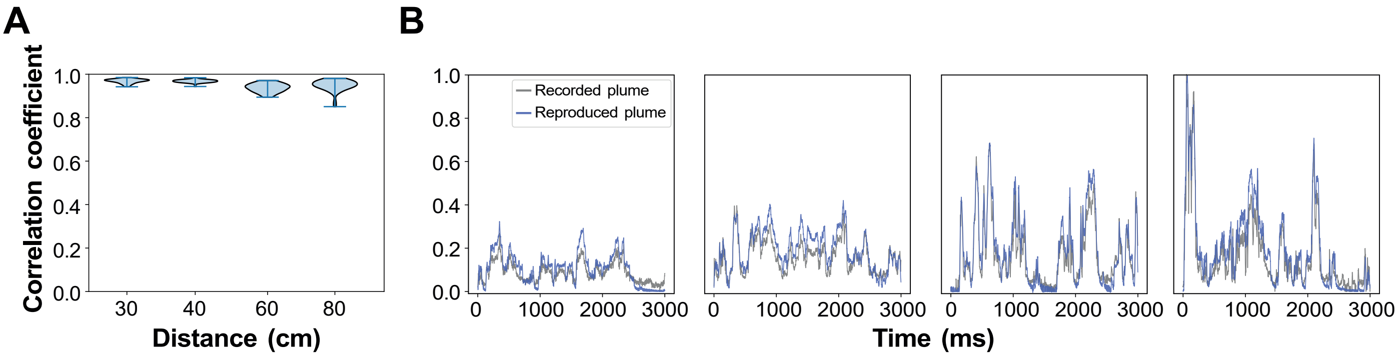


**Figure S2.9: Plume reproduction fidelity. (A)** Correlation coefficients of recorded plumes and reproduced plumes presented in the imaging experiment. Violin plot showing kernel density estimation of the underlying distribution, lines represent minimum and maximum (n = 40 plumes/distance). **(B)** Example plumes from different distances (L-R: 80 cm, 60 cm, 40 cm, 30 cm), recorded plumes in grey, reproduced plumes in blue. Traces normalised to the maximum of the 4 traces to aid the visualisation

### Figure S2.10

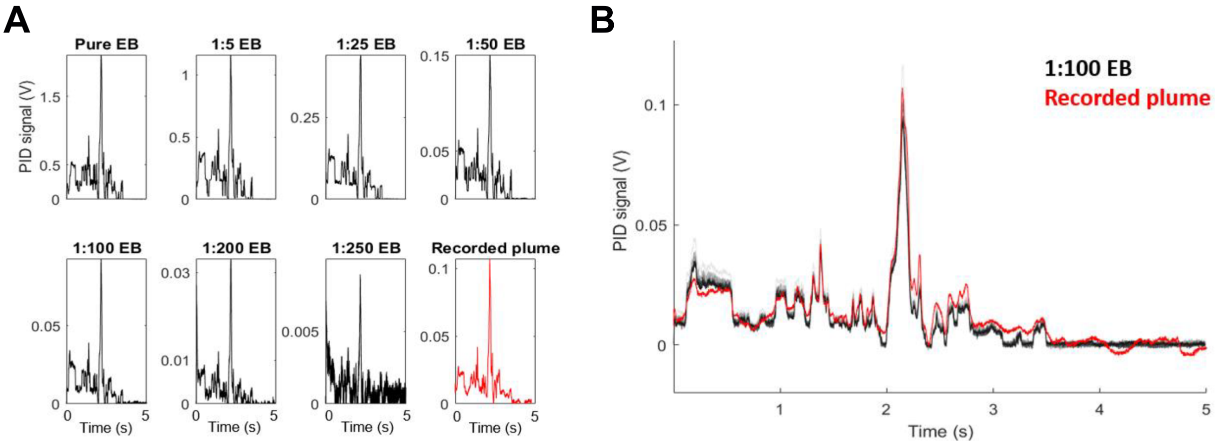


**Figure S2.10: Replicating recorded plume structure at different concentrations. (A)** Examples of replicated plumes at different concentrations of odour loaded into the tODD, Ethyl Butyrate diluted in Mineral Oil (black traces) and the recording used to produce them (red trace). **(B)** Plume structure can be reliably replicated over multiple trials (n = 10) at 1:100 EB (light grey = first trial, black = last trial), which produces a signal similar in amplitude to the recorded plume (red trace). Different levels of shading indicate signal evolution over time, the black trace being the most recent trace, approximately 2 min between repeats.

### Figure S2.11

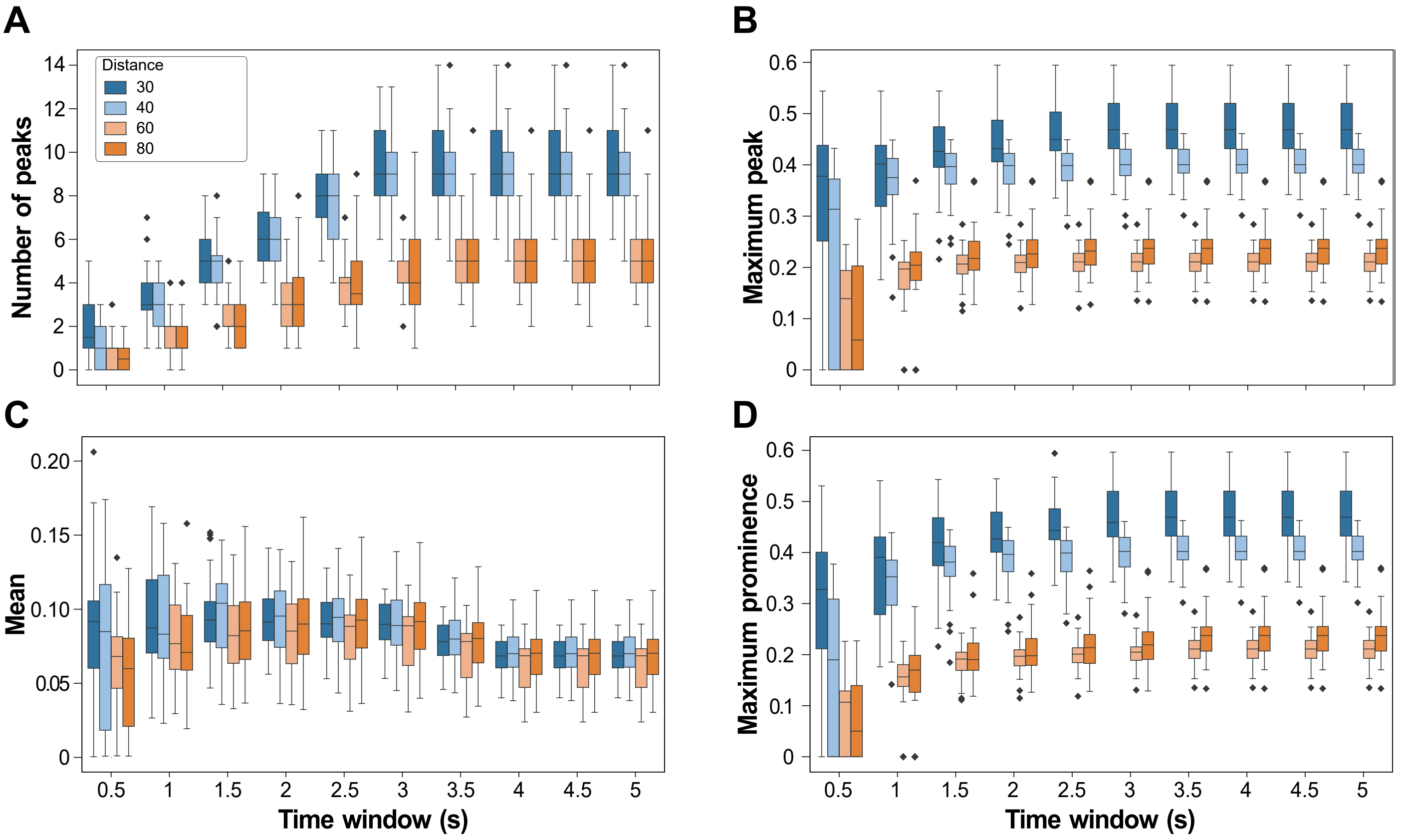


**Figure S2.11: Temporal structure features of reproduced plumes from different distances.** Box plots of plume features for intervals of increasing length between time = 0 s, as defined by odour arrival, and different time points as shown on the x axis for the entire length of the response window considered for imaging analysis (5 s). Colours indicate source distance (80 cm orange, 60 cm pale orange, 40 cm pale blue, 30 cm blue). Box indicates 25th-75th percentiles, thick line is median, whiskers are most extreme data points not considered outliers. Number of peaks **(A)**, maximum peak **(B)** and maximum prominence **(D)** are different between 30 cm and 80 cm for all time windows ($p < 0.05$, Bonferroni-corrected Mann-Whitney U test). Mean concentration **(C)** is significantly different between 30 cm and 80 cm only at times 0.5 s and 1 s.

### Figure S2.12

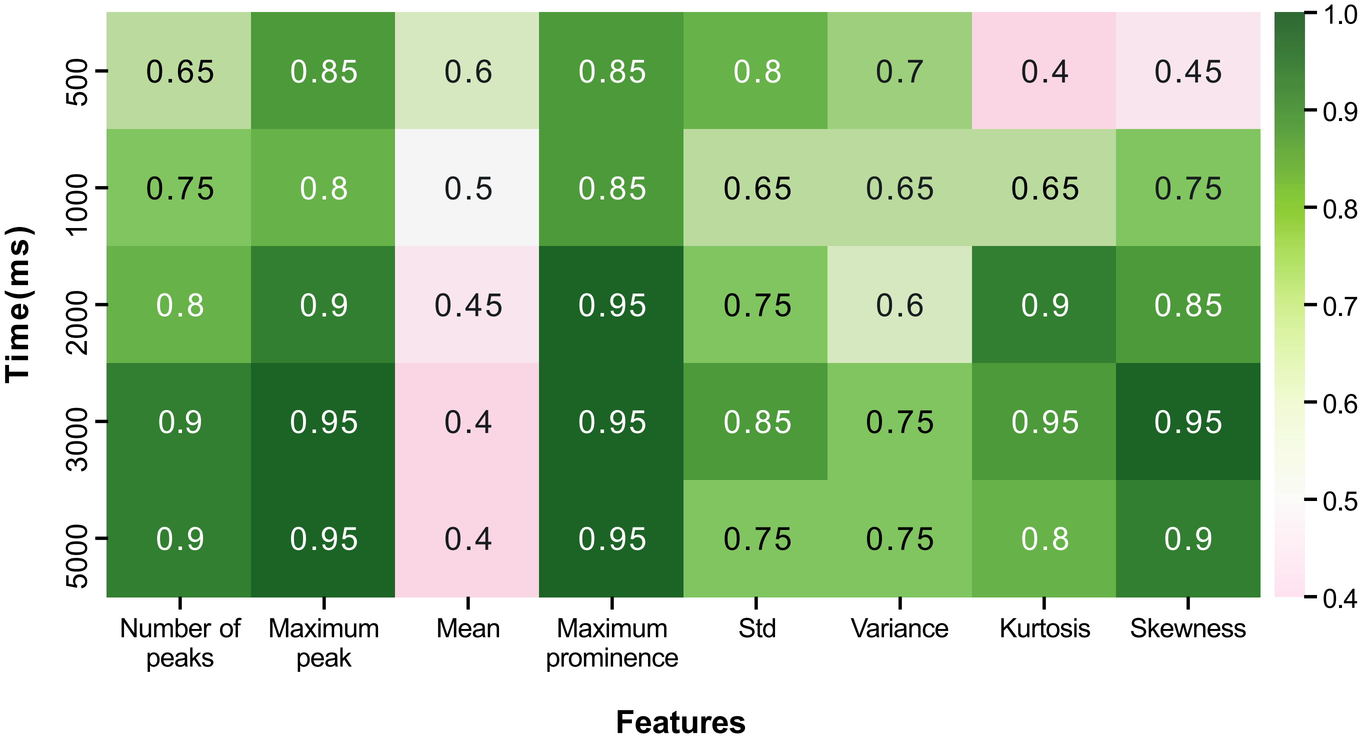


**Figure S2.12: Accuracy of 30 cm vs 80 cm discrimination for reproduced plumes.** Classification accuracy on a test set of 45 plumes for a linear discriminant trained on 135 plumes, using only one feature (columns) and one time window (rows) at a time. The split was stratified by distance such that equal numbers of 30 cm and 80 cm plumes were present in the training and the test set. Same analysis for the reproduced plume as previously shown for the recorded plumes in Figure S2.4.

### Figure S3.1

A

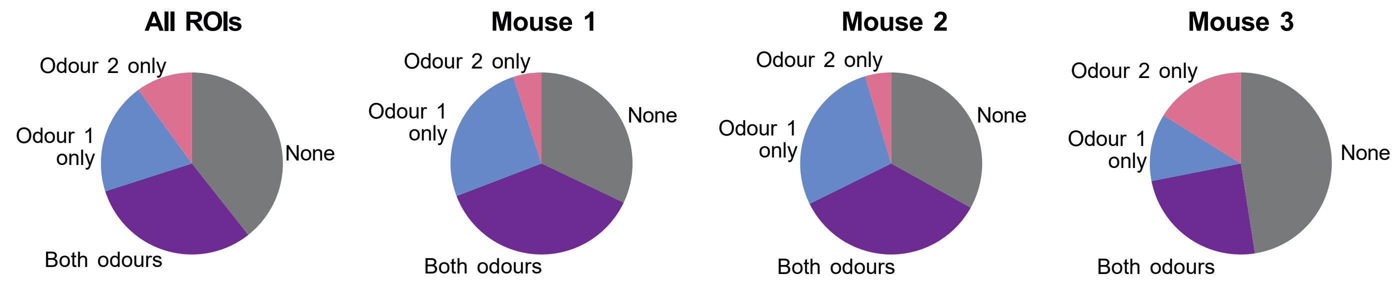


B

| | All | Mouse 1 | Mouse 2 | Mouse 3 |
|---|---|---|---|---|
| **Total** | 531 | 159 | 130 | 242 |
| **None** | 209 | 51 | 43 | 115 |
| **EB only** | 106 | 41 | 36 | 29 |
| **2H only** | 53 | 8 | 6 | 39 |
| **Both odours** | 163 | 59 | 45 | 59 |

C

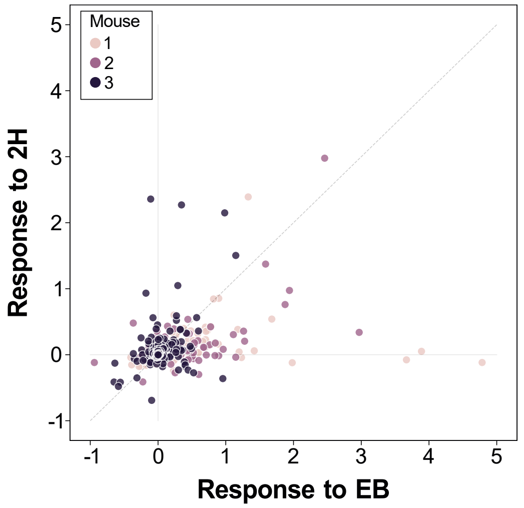


D

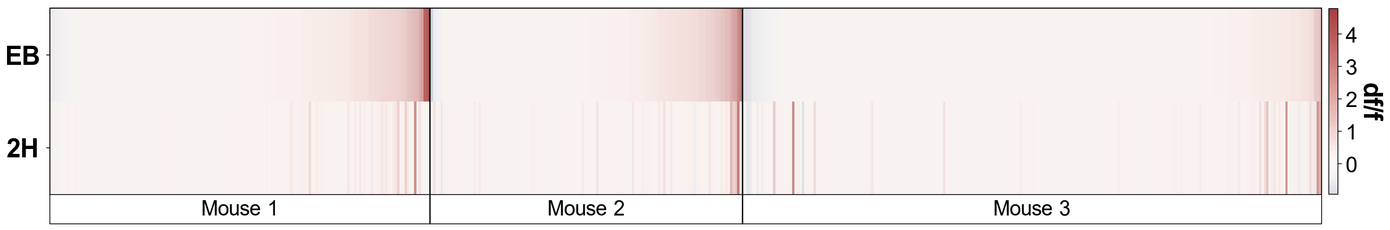


**Figure S3.1: Response profiles to EB and 2H of recorded cells. (A)** Proportion of ROIs responsive only to EB (Odour 1), 2H (Odour 2), both odours, or neither, for all mice pooled together (left) or for each mouse individually (right). A cell was classed as "responsive" if mean fluorescence during the response window for all trials of an individual odour was above mean + 3*s.d. of the period 3 s before stimulus onset. **(B)** Values used to calculate the pie-charts in A. **(C)** Odour preference (EB or 2H) for recorded cells, colour-coded by mouse. In this case, "response" is the mean value of the fluorescence trace over a 5 s sampling window from the trial initiation. **(D)** Mean response to individual odours (rows, top = EB, bottom = 2H) for all cells (columns), split by mouse, then ordered by increasing response to EB (Odour 1).

### Figure S3.2

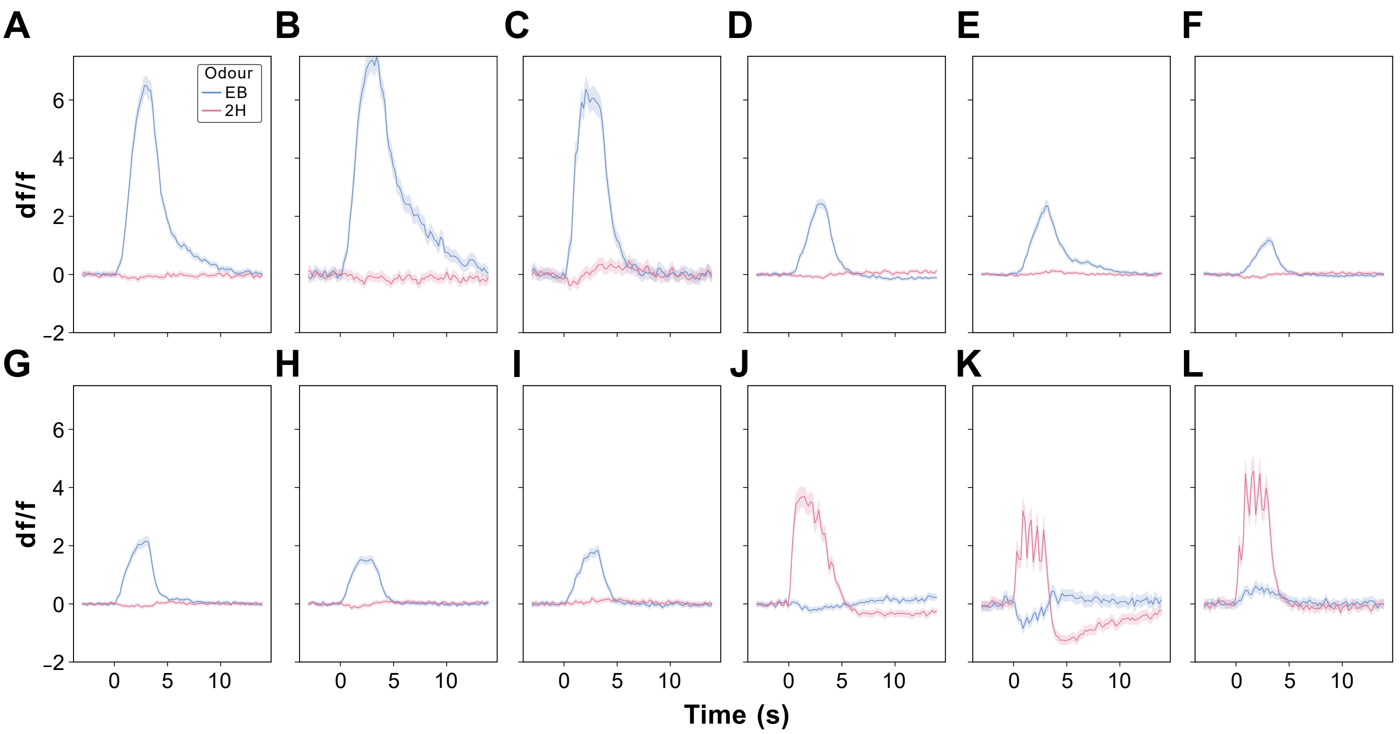


**Figure S3.2: Responses to individual odours from example cells.** Average fluorescence traces (mean ± s.e.m., n = 352 trials, except for mouse 2, for which n = 265 trials) for all trials of an individual odour, regardless of distance, for 12 example cells, split by odour: Odour 1 = EB, blue; Odour 2 = 2H, violet. Odour delivery period from 0-3 s. Cells in A-F were recorded from mouse 1, cells in G-I from mouse 2 and cells in J-L from mouse 3.

### Figure S3.3

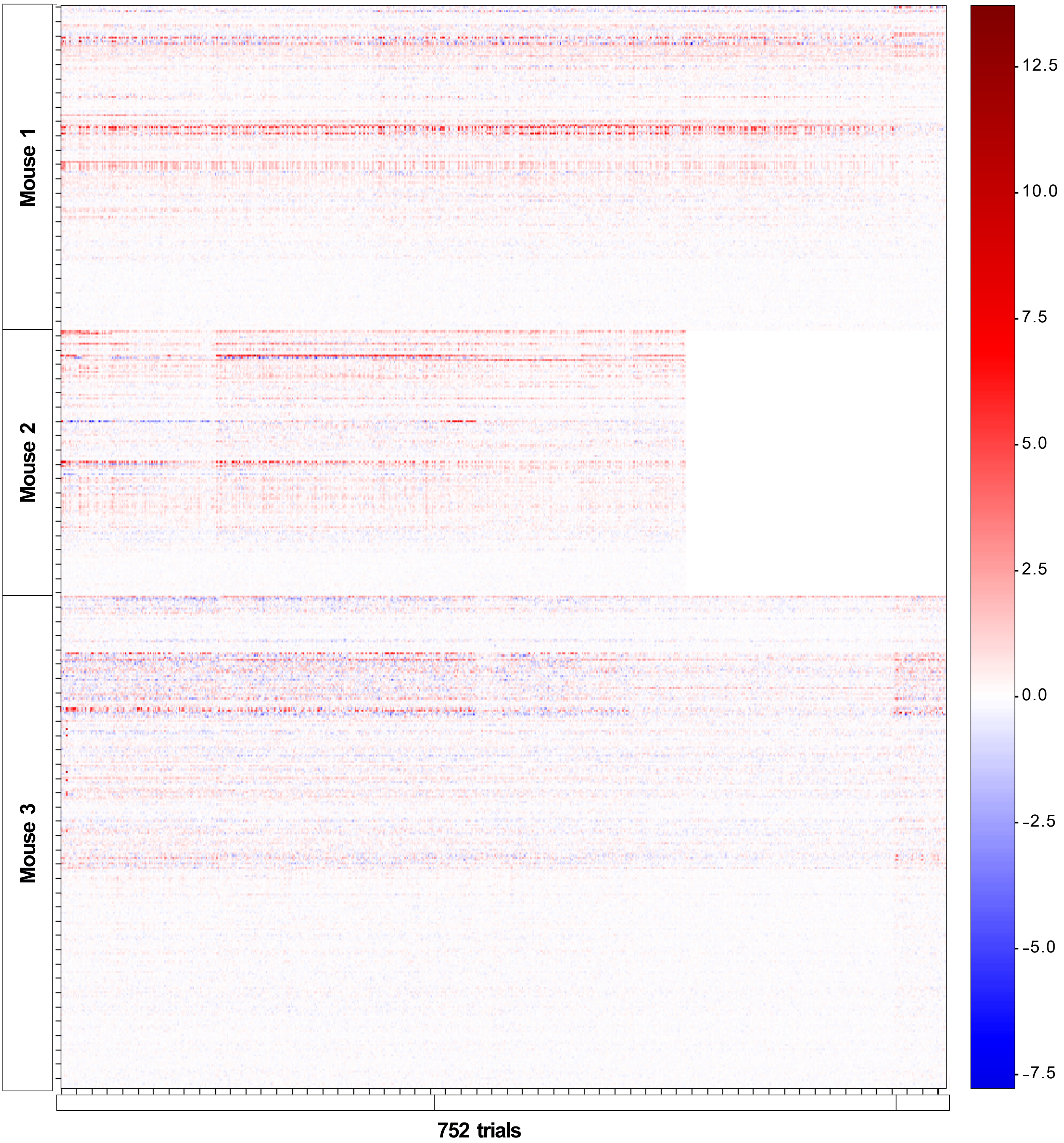


| Core plumes | Additional repeats | Additional odours |
|---|---|---|
| 40 plumes x 4 distances x 2 odours | 40 plumes x 4 distances x 2 odours + 1 plume x 4 distances x 2 odours x 8 repeats | 1 plume x 4 distances x 4 odours x 3 repeats |
| = 320 trials, randomised | = 384 trials, randomised | = 48 trials, randomised |

**Figure S3.3: Overview of experiment timeline and cell responses.** Heatmap of responses for each ROI (rows) for each trial (columns). Values represent mean df/f over the 5 s response window. ROIs are ordered by mouse and cell ID. Trial order is described in the table underneath. Mouse 2 did not reach the end of the experiment.

### Figure S3.4

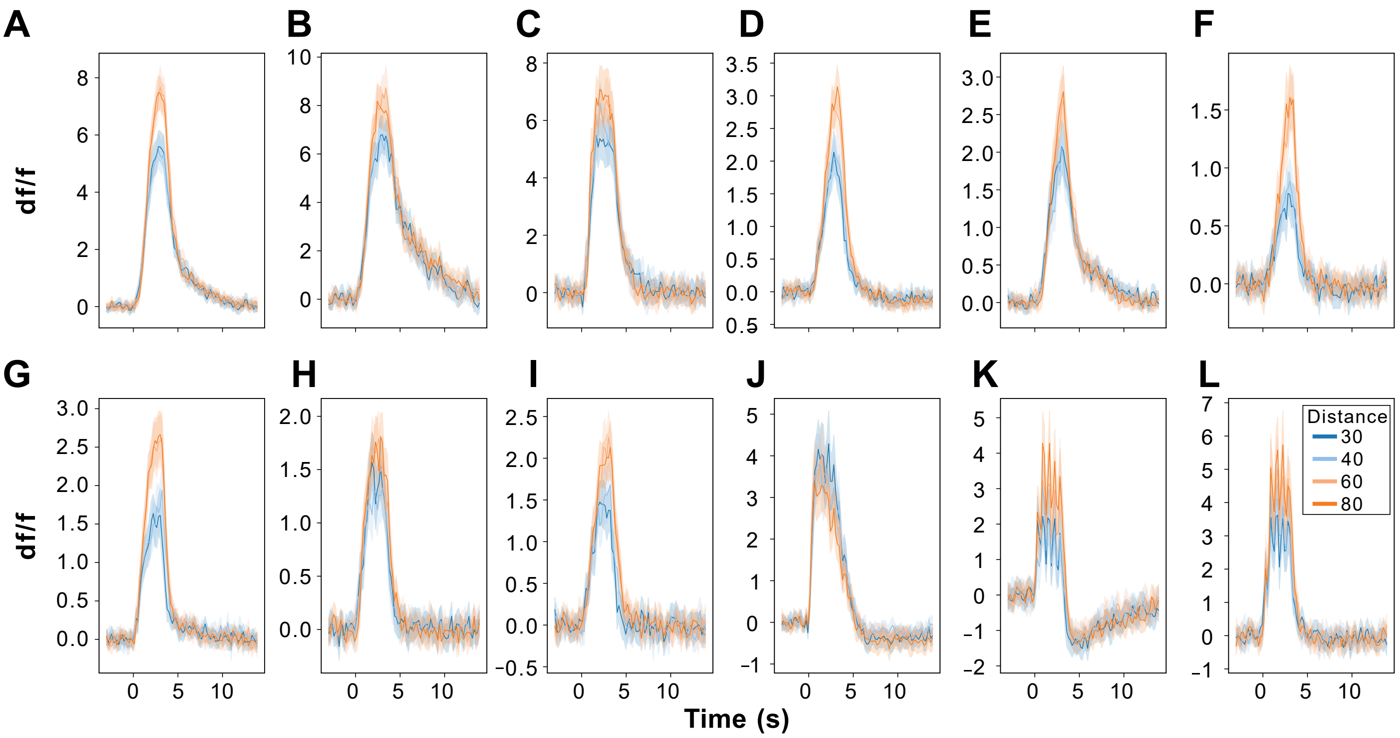


**Figure S3.4: Average responses of distance-sensitive cell-odour pairs.** Average fluorescence traces (mean ± s.e.m., n = 160 trials) for 12 example cells (same as Figure 3) colour-coded for the 4 distances (30 cm blue, 40 cm pale blue, 60 cm pale orange, 80 cm orange), for one of the two main odours only: Odour 1 (EB) for cells A-I and Odour 2 (2H) for cells in J-L. Odour stimulation (shaded area) was 3 s long and started at time = 0, which is also the onset of inhalation.

### Figure S3.5

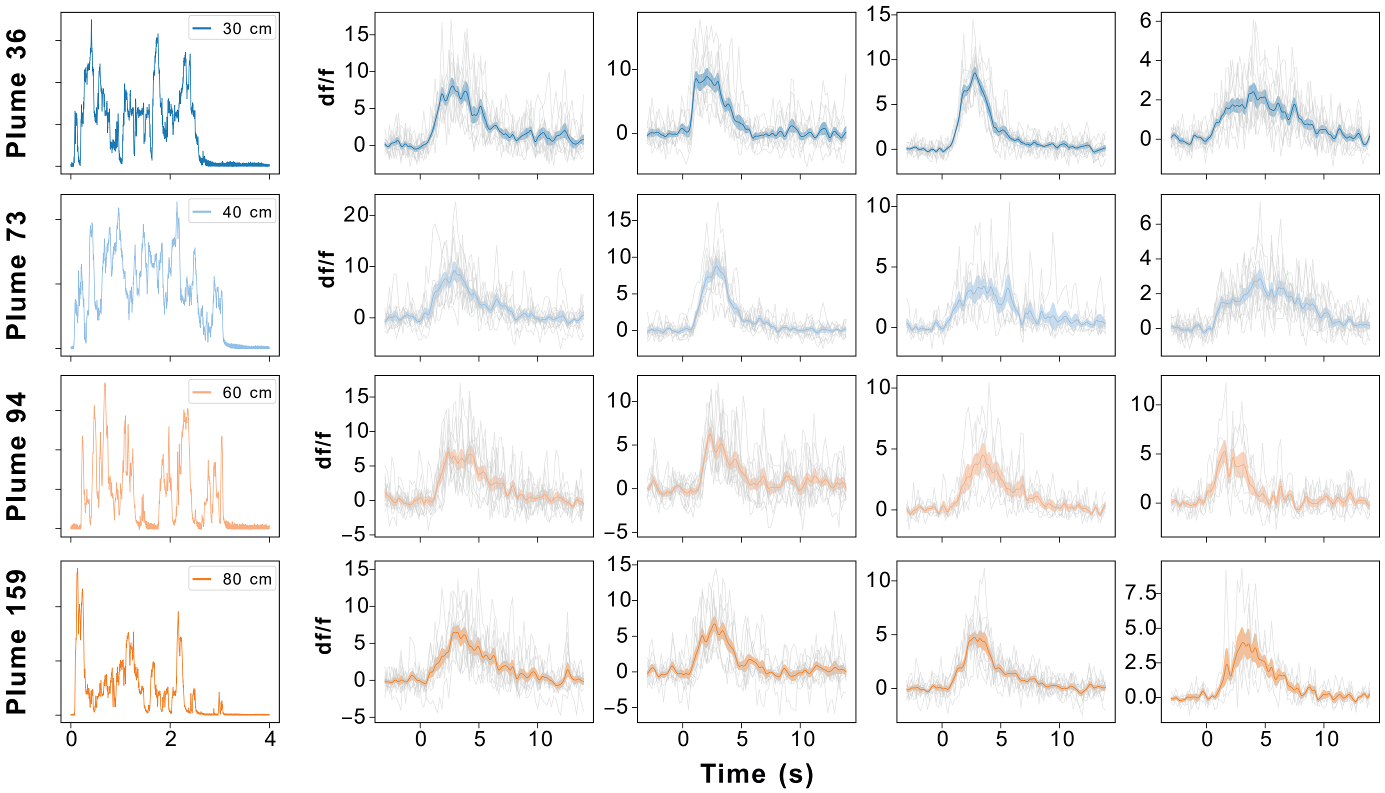


**Figure S3.5: Responses from example cells to plumes presented multiple times.** Example plume traces recorded from 4 distances (left column) that were presented for 10 repetitions and corresponding fluorescence traces of example cells (four remaining columns). Individual trials are shown in light grey, mean trace ± s.e.m colour-coded according to plume distance: (30 cm (dark blue), 40 cm (light blue), 60 cm (light orange) and 80 cm (dark orange).

### Figure S4.1

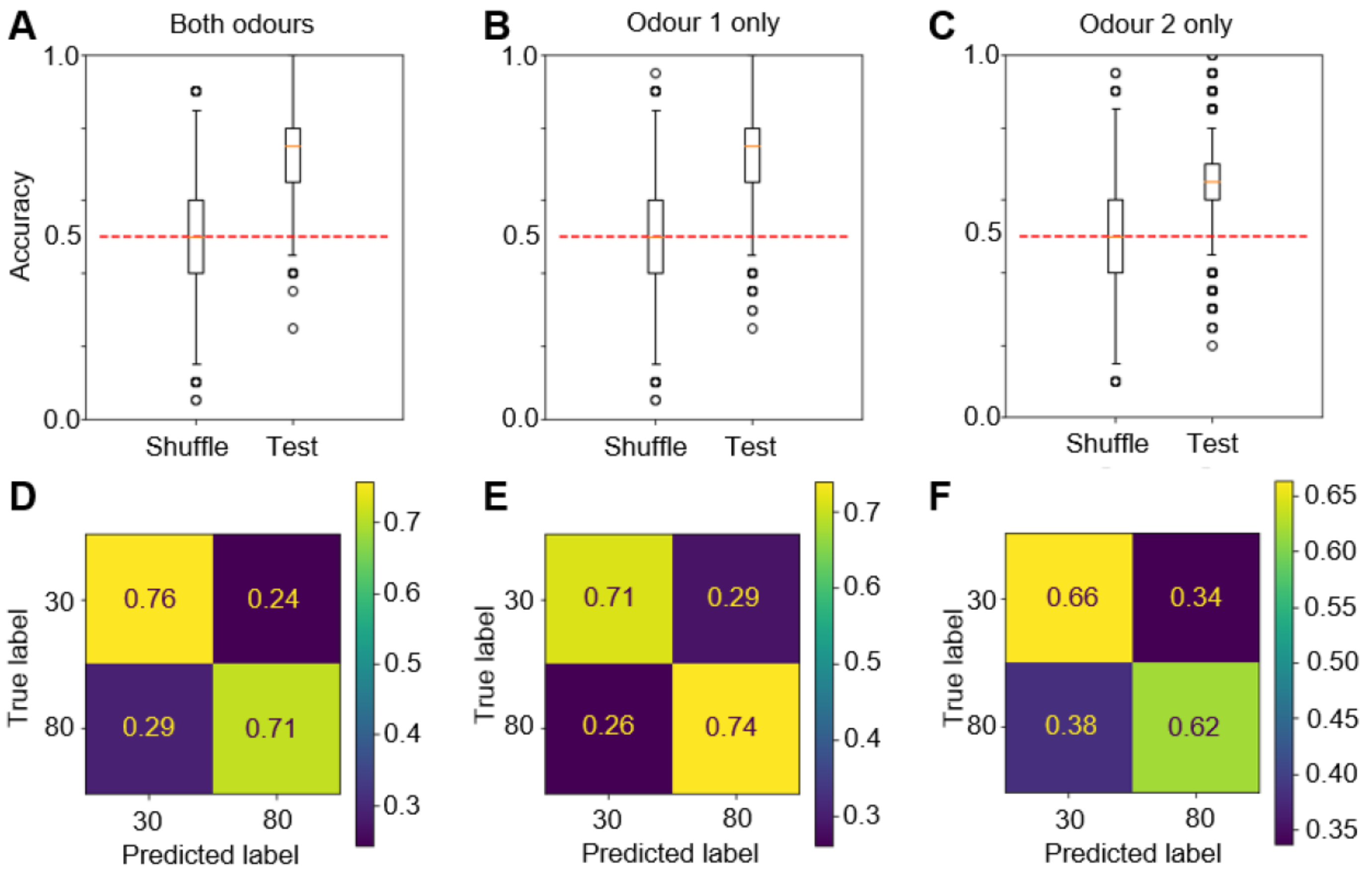


**G**

| **30 vs 80 Accuracy (mean±sd)** | **All mice** | **Mouse 1** | **Mouse 2** | **Mouse 3** |
|---|---|---|---|---|
| **Both odours** | **0.73±0.09 *** | 0.71±0.10 * | 0.59±0.10 * | 0.67±0.09 * |
| **Odour 1** | **0.73±0.09 *** | 0.65±0.10 * | 0.58±0.10 * | 0.62±0.09 * |
| **Odour 2** | **0.64±0.10 *** | 0.59±0.10 * | 0.54±0.10 * | 0.64±0.10 * |

**Figure S4.1: Population classifier results – binary classification 30 cm vs 80 cm. (A-C)** Accuracy of trained classifiers on the test set when trained on labelled (right) or shuffled data (left). Chance = 50%. Box indicates 25th-75th percentiles, thick line is median, whiskers are most extreme data points not considered outliers. Repeated 50000 times after a 75:25 training:test split out of a total of 80 trials (40 per distance) and 1062 cell-odour pairs (**A**) or 531 cells when individual odours were trained independently (**B,C**). **(D-F)** Average confusion matrices for the classifiers shown in A-C. **(G)** Table summarising the accuracy (mean±s.d.) obtained by training classifiers on data from all or individual mice, or all or individual odours. Stars indicate significant difference from the shuffled accuracy score, $p < 0.05$, Mann Whitney U test.

### Figure S4.2

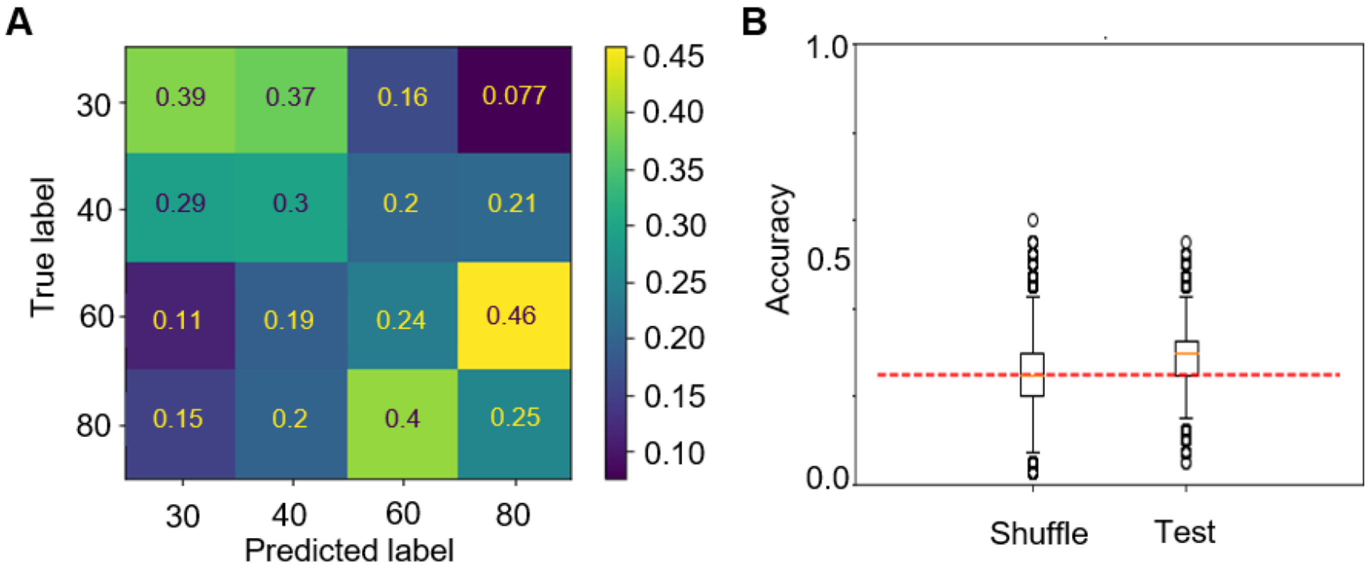


**C**

| 4 way classification Accuracy (mean±sd) | All mice |
|---|---|
| **Both odours** | **0.29±0.06 *** |
| Odour 1 | 0.30±0.06 * |
| Odour 2 | 0.28±0.06 * |

**Figure S4.2: Population classifier results – 4-way classification. (A)** Average confusion matrices for classifiers trained to classify trials into the 4 distances used. **(B)** Accuracy of trained classifiers on the test set when trained on labelled data (right) or shuffled data (left). Chance = 25%. Box indicates 25th-75th percentiles, thick line is median, whiskers are most extreme data points not considered outliers. Repeated 50000 times after a 75:25 training:test split out of a total of 160 trials (40 per distance) and 1062 cell-odour pairs **(C)** Table summarising the accuracy (mean ± s.d.) obtained by training classifiers on data from both or individual odours, from all mice. Stars indicate significant difference from the shuffled accuracy score, $p < 0.05$, Mann-Whitney U-test

### Figure S4.3

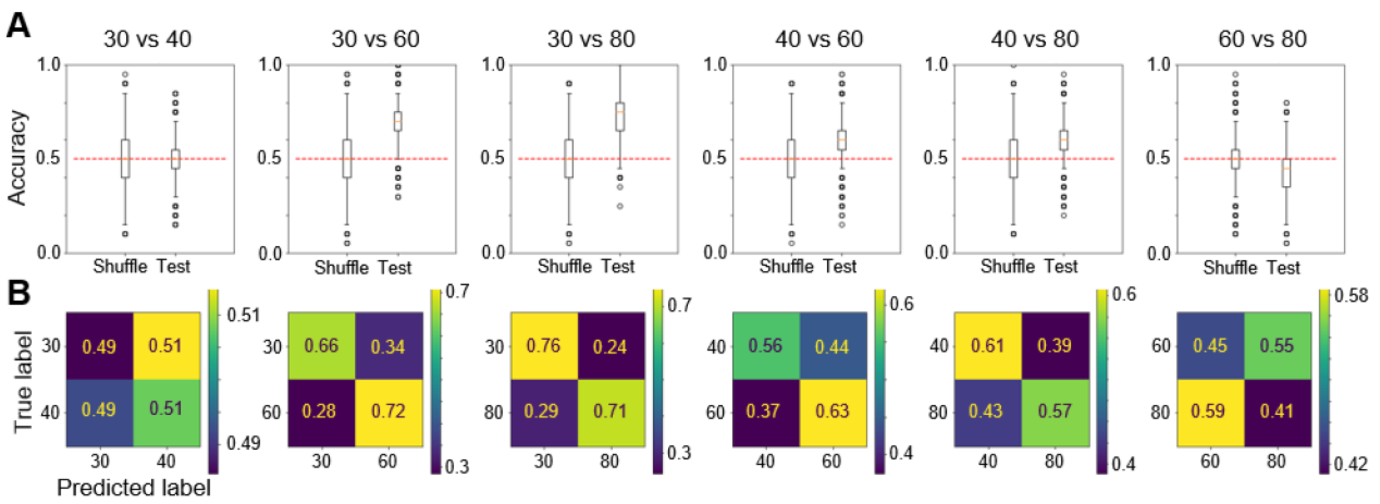


**C**

| | 30 vs 40 | 30 vs 60 | 30 vs 80 | 40 vs 60 | 40 vs 80 | 60 vs 80 |
|---|---|---|---|---|---|---|
| **Accuracy (mean±sd)** | 0.50±0.10 | 0.69±0.09 * | 0.73±0.09 * | 0.59±0.10 * | 0.59±0.09 * | 0.43 ± 0.10 |

**D**

| **Accuracy (mean±sd)** | **30** | **40** | **60** | **80** | |
|---|---|---|---|---|---|
| **30** | | 0.49±0.10 | 0.67±0.09 * | 0.64±0.10 * | Odour 2 |
| **40** | 0.51±0.09 * | | 0.55±0.10 * | 0.55±0.10 * | |
| **60** | 0.60±0.10 * | 0.56±0.10 * | | 0.44±0.10 | |
| **80** | 0.73±0.09 * | 0.62±0.09 * | 0.45±0.09 | | |
| | Odour 1 | | | | |

**Figure S4.3: Population classifier results – pairwise comparisons (A)** Each plot shows accuracy of trained classifiers on the test set when trained on labelled data (right) or on shuffled data (left), for a specific distance pair: 30vs40, 30vs60, 30vs80, 40vs60, 60vs80, 60vs80. Chance is at 50%. Box indicates 25th-75th percentiles, thick line is median, whiskers are most extreme data points not considered outliers. Repeated 50000 times after a 75:25 training: test split out of a total of 80 trials (40 per distance) and 1062 cell-odour pairs. **(B)** Average confusion matrices for the classifiers in A. **(C)** Table summarising the accuracy (mean±s.d.) of classifiers in **A**, data from all mice and both odours (n = 1062 cell-odour pairs). **(D)** Table summarising the accuracy (mean±s.d.) of classifiers similar to those in **A** but trained on data from individual odours (n = 531 cells), for Odour 1 (EB, blue) or Odour 2 (2H, violet). Stars indicate significant difference from the shuffled accuracy score, $p < 0.05$, Mann-Whitney U-test.

### Figure S4.4

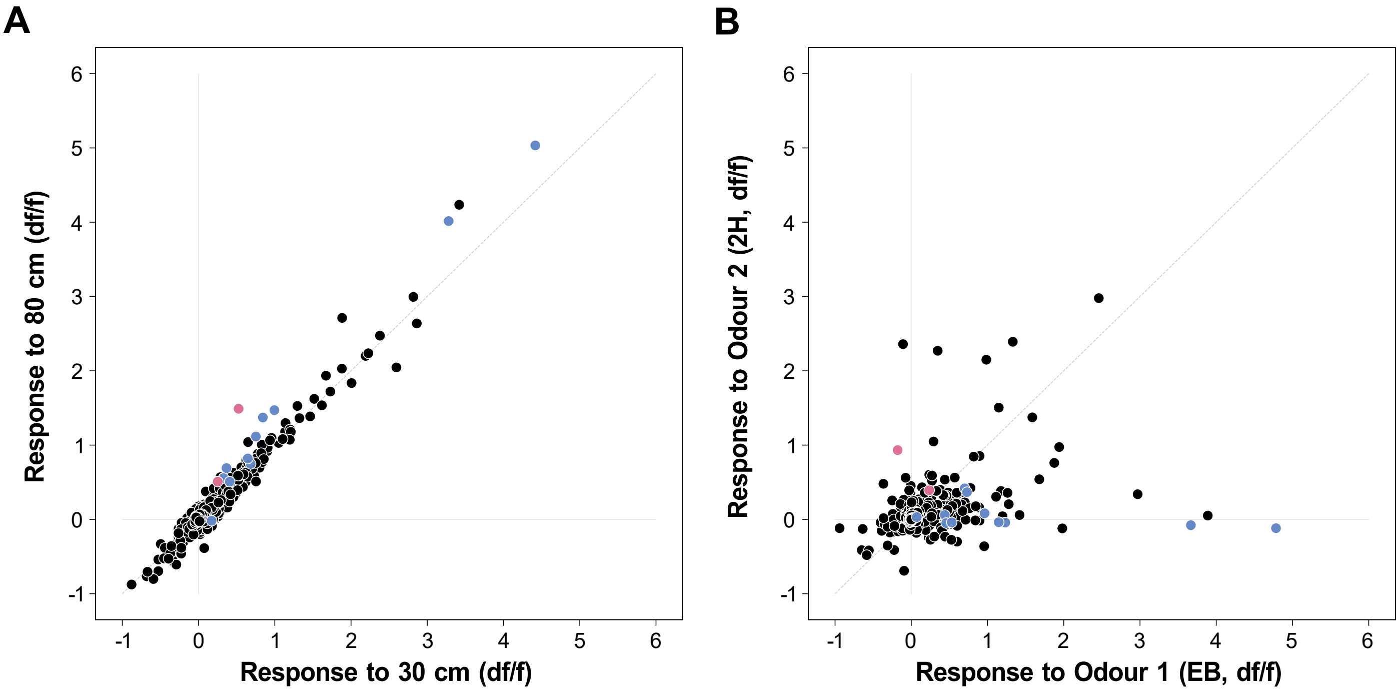


**Figure S4.4: Characterisation of distance sensitive cells. (A)** Responses to 30 cm vs responses to 80 cm plotted as mean response for all trials from those distances. Distance-sensitive cells colour-coded by the odour for which they are distance-sensitive: Odour 1 (EB) blue, Odour 2 (2H) violet. Cells above diagonal respond more strongly to 80 cm while cells below diagonal respond more strongly to 30 cm plumes. The majority of cells are only weakly responsive and/or show very small differences. **(B)** Odour preference as in Figure S3.1C, but with distance-sensitive cells colour-coded by the odour for which they were classified as distance-sensitive: Odour 1 (EB) blue, Odour 2 (2H) violet. Each dot is a cell. Cells that are distance-sensitive for Odour 1 respond strongly to Odour 1 but not Odour 2, while cells that are distance-sensitive for Odour 2 respond strongly to Odour 2 but not Odour 1. Cells that respond strongly to both odours are not distance-sensitive (black).

### Figure S4.5

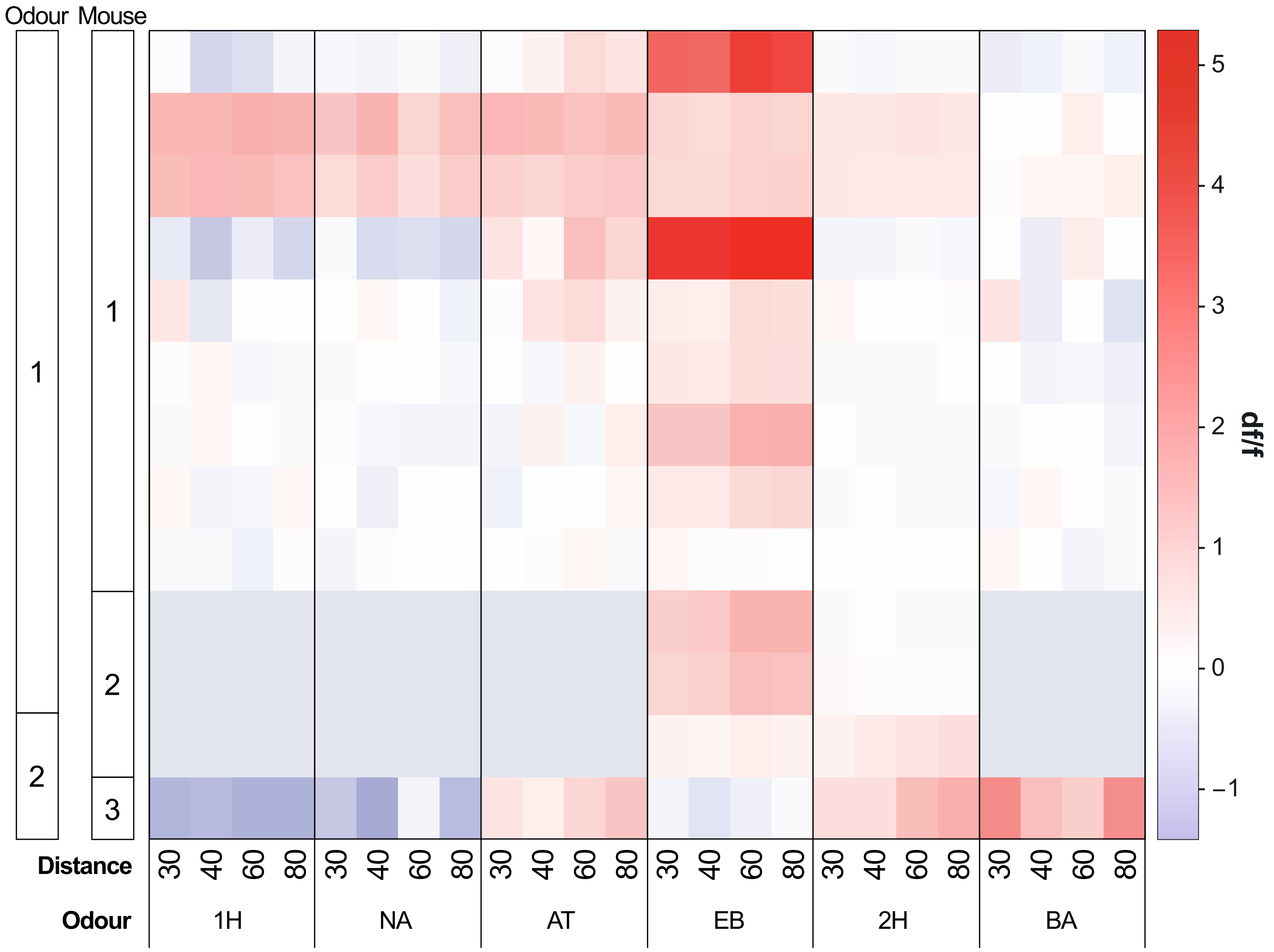


**Figure S4.5: Responses of distance-sensitive cells to odours and distances.** Heatmap of responses for each odour (big columns) further subdivided by distance (small columns) for all distance-sensitive cells (rows). Cells are ordered first by the odour for which they are distance sensitive (top: EB, bottom: 2H), then by mouse, then by cell ID. Grey areas indicate the missing trials with the additional 4 odours that were not presented for mouse 2.

### Figure S5.1

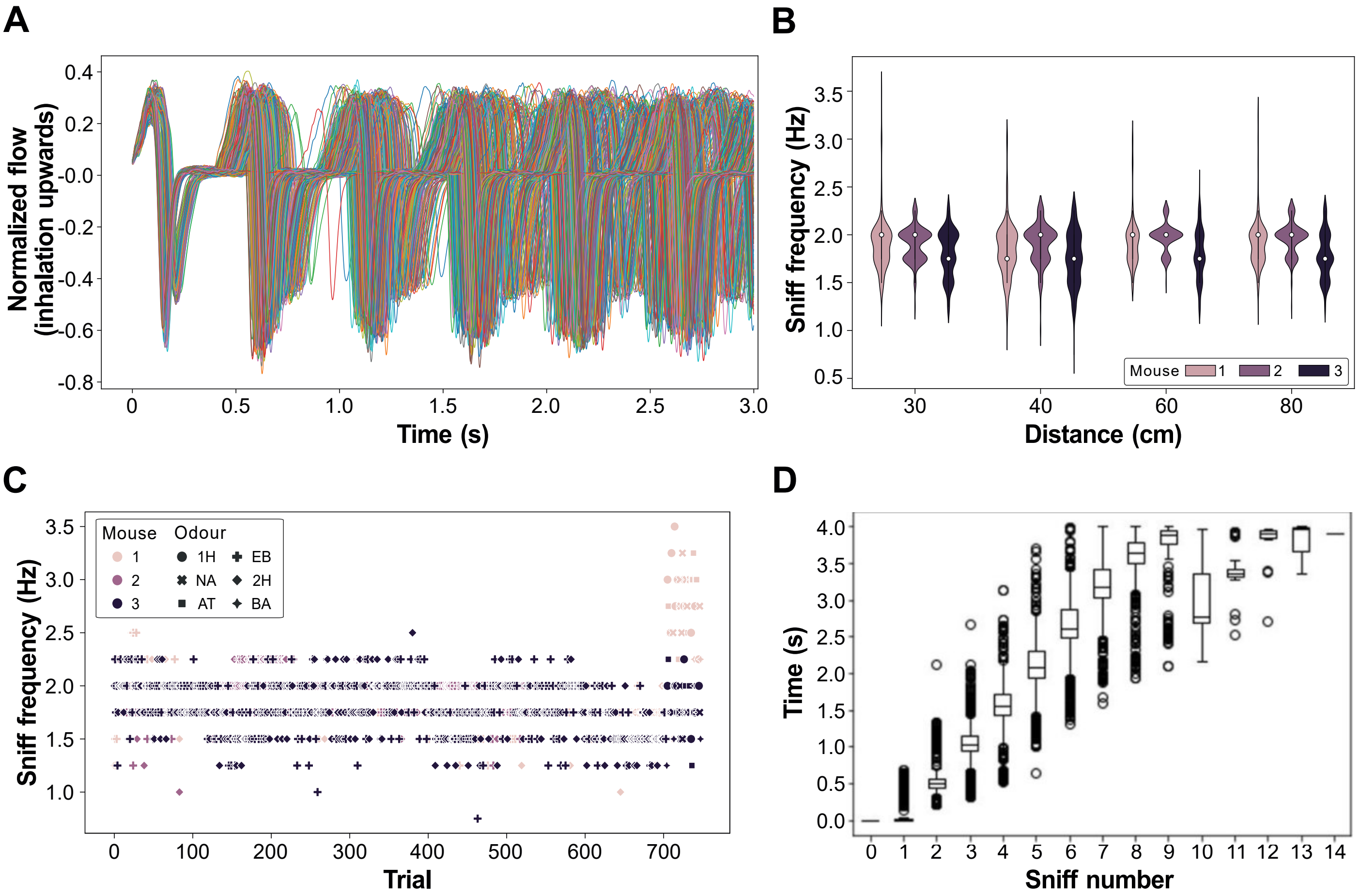


**Figure S5.1: Respiration stability across experiments. (A)** Respiration traces during odour presentation (3 s) as recorded with a flow sensor placed in front of the contralateral naris. All trials from a single mouse (mouse 2), each line is a trial (n = 532), inhalation pointing upwards. Traces normalised to the maximum amplitude during the experiment. Trials aligned to time = 0 ms which is the start of inhalation, as detected using a threshold crossing approach. The first sniff shows high synchrony across trials, with decrease in synchrony for subsequent sniffs. **(B)** Sniff (respiration) frequency for each trial, split by mouse (colours) and by distance (x-axis). **(C)** Sniff frequency (y-axis) for each trial (x-axis) for the length of the experiment. Each dot is a trial, colour-coded by mouse; different symbols indicate different odour identities. **(D)** Timing of sniff onsets for the 5s response window considered for the analysis of fluorescence traces. First sniff is always at 0 ms. Box indicates 25th-75th percentiles, thick line is median, whiskers are most extreme data points not considered outliers.

### Figure S5.2

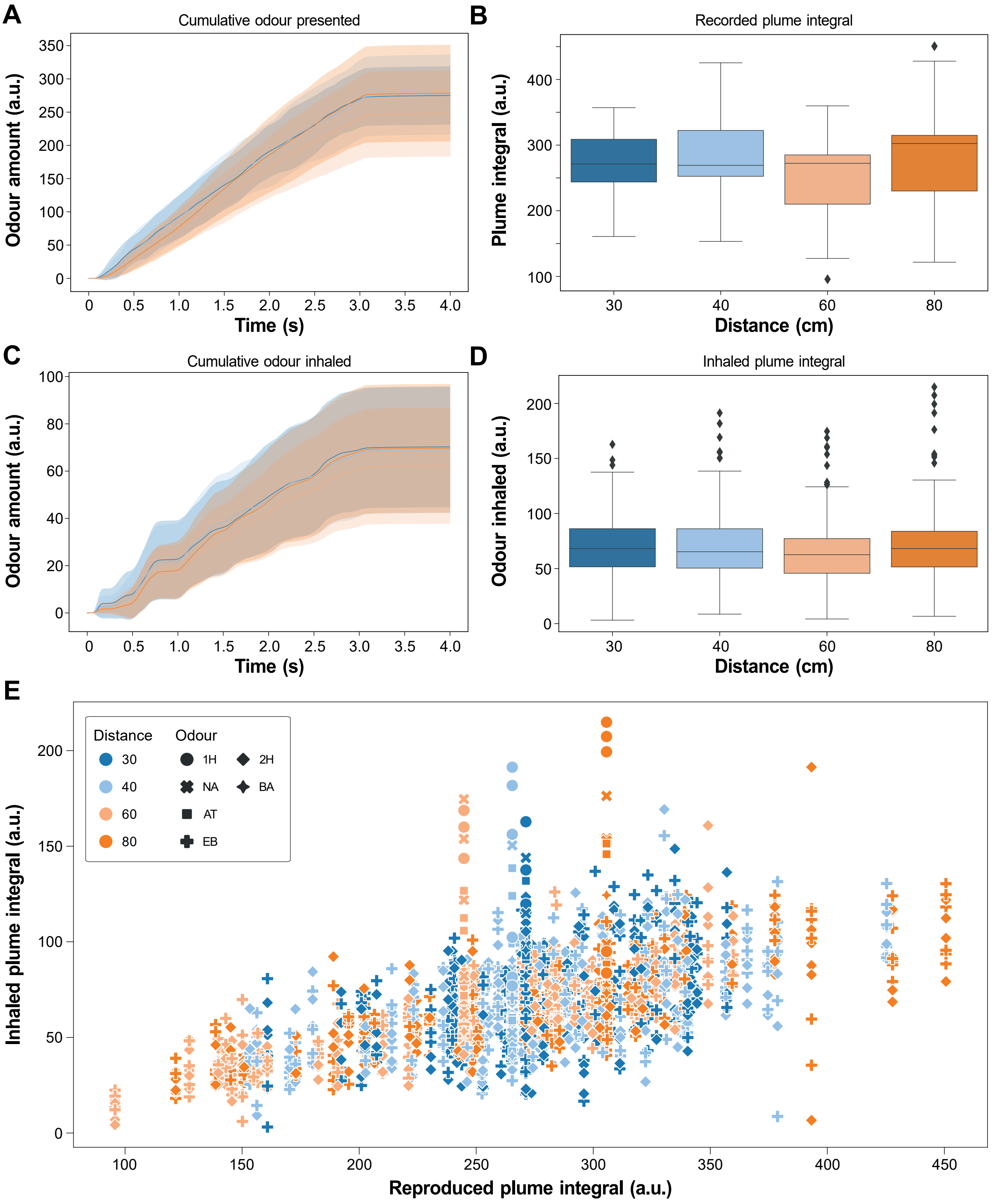


**Figure S5.2: Odour amount presented and inhaled. (A)** Odour amount (a.u.) reaching the animal calculated as the integral of the plume trace (PID signal) for time windows of increasing size, from the start of the trial, which is also the start of inhalation, until the corresponding point on the x-axis. The delay for the delivered odour to reach the animal is 65 ms and odour delivery lasts 3 s. Colour-coded for distance (blue: 30 cm, pale blue: 40 cm, pale orange: 60 cm, orange: 80 cm), mean ± s.d. **(B)** Box plot showing odour amount for the entire response window (5 s) since trial start for plumes for each distance. Box indicates 25th-75th percentiles, thick line is median, whiskers are most extreme data points not considered outliers. Trials from 60 cm have a lower amount of odour presented compared to all other distances, but no other differences reach significance levels ($p < 0.05$, Mann–Whitney U test). (**C)** Same as **A** but for the *inhaled plume* (Methods) (**D)** Same as **B** but for the *inhaled plume*. Trials from 60 cm have a lower amount of odour inhaled compared to all other distances, but no other differences reach significance levels ($p < 0.05$, Mann–Whitney U test). **(E)** Scatter plot of measures in **B** and **D** showing the relationship between the odour amount presented and inhaled. Colours represent distance categories; symbols represent different mice.

### Figure S5.3

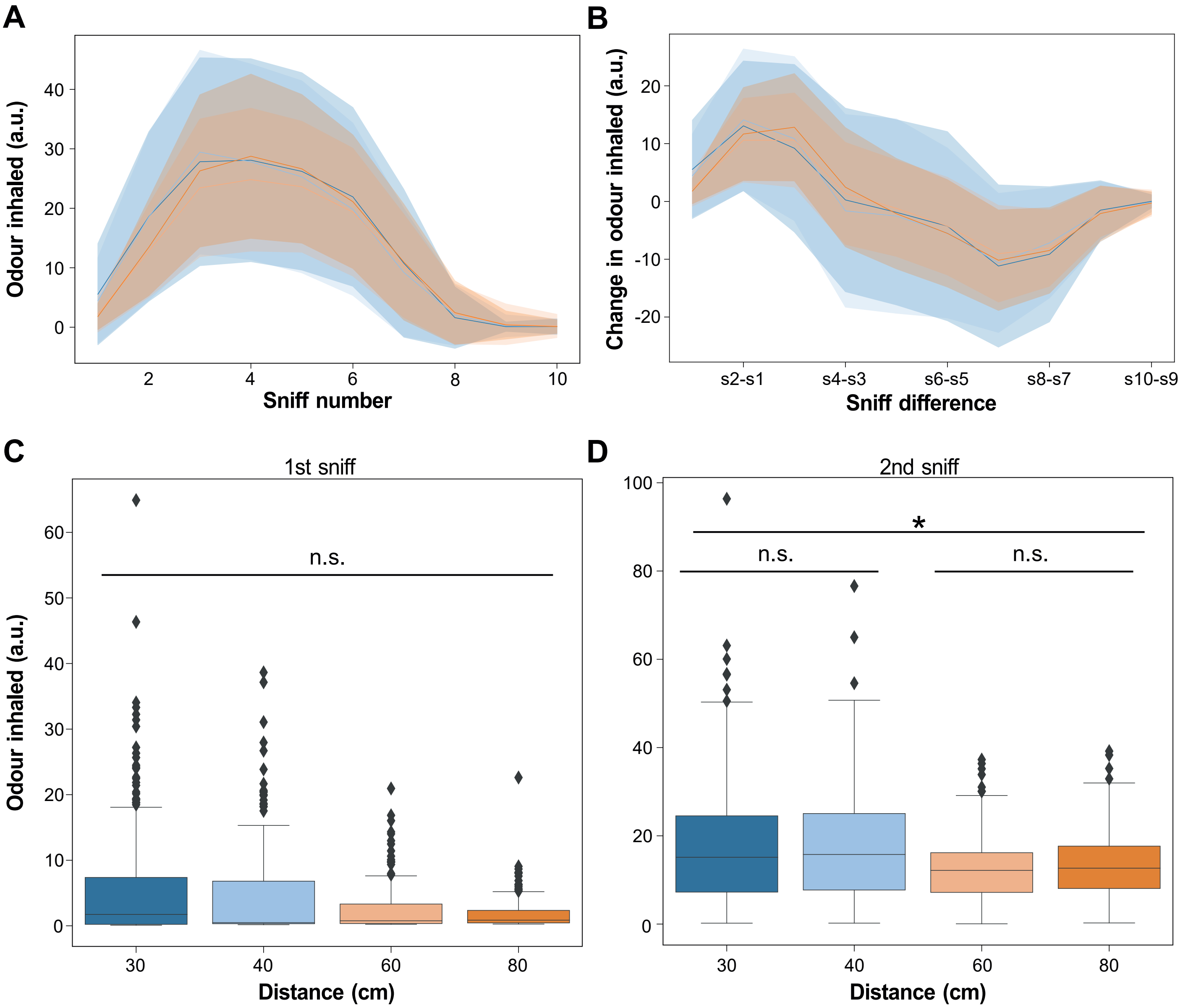


**Figure S5.3: Odour concentration in sniff time. (A)** Odour amount (a.u.) as inhaled by the mouse for each sniff calculated as the integral of the inhaled plume trace (PID signal) Colour-coded for distance (blue: 30 cm, pale blue: 40 cm, pale orange: 60 cm, orange: 80 cm), mean ± s.d. **(B)** Change in the amount of odour inhaled between consecutive sniffs, calculated as the difference between consecutive sniffs in **A.** Sniff number 1 corresponds to the odour inhaled in the first sniff, sniff number 2 corresponds to the difference between the second and the first sniff etc. **(C)** Box plot showing odour inhaled for the first sniff. Box indicates 25th-75th percentiles, thick line is median, whiskers are most extreme data points not considered outliers. No significant difference between groups (Kruskal-Wallis test). **(D)** Same as **C** but for the second sniff. More odour was inhaled during near trials (30 cm and 40 cm) than in far trials (60 cm and 80 cm), when performing pairwise comparisons ($p < 0.05$, Kruskal-Wallis test followed by post-hoc Mann-Whitney U test with Bonferroni correction). Differences within a distance group (30 cm vs 40 cm or 60 cm vs 80 cm) were not significant).
